# Chainsaw: protein domain segmentation with fully convolutional neural networks

**DOI:** 10.1101/2023.07.19.549732

**Authors:** Jude Wells, Alex Hawkins-Hooker, Nicola Bordin, Ian Sillitoe, Brooks Paige, Christine Orengo

## Abstract

**Motivation:** Protein domains are fundamental units of protein structure and play a pivotal role in understanding folding, function, evolution, and design. The advent of accurate structure prediction techniques has resulted in an influx of new structural data, making the partitioning of these structures into domains essential for inferring evolutionary relationships and functional classification.

**Results:** This manuscript presents Chainsaw, a supervised learning approach to domain parsing that achieves accuracy that surpasses current state-of-the-art methods. Chainsaw uses a fully convolutional neural network which is trained to predict the probability that each pair of residues is in the same domain. Domain predictions are then derived from these pairwise predictions using an algorithm that searches for the most likely assignment of residues to domains given the set of pairwise co-membership probabilities. Chainsaw matches CATH domain annotations in 78% of protein domains versus 72% for the next closest method. When predicting on AlphaFold models expert human evaluators were twice as likely to prefer Chainsaw’s predictions versus the next best method.

**Availability and Implementation:** Code implementation of Chainsaw is available at github.com/JudeWells/chainsaw.

## 1 Introduction

Protein domains are generally defined as self-stabilizing units composed of several secondary structural elements that pack together to form a hydrophobic core. From an evolutionary perspective, the protein domain is the level at which homology and functional groups are understood. Structural protein domain databases such as CATH (1), SCOP (2), SCOPe (3) and ECOD (4) are essential for advancing scientific understanding of the protein universe. In 2022 DeepMind released the AlphaFold models for over 200 million proteins, increasing the number of available structures by multiple orders of magnitude (5). These databases present opportunities to discover novel domains, infer evolutionary links and generate functional hypotheses but an essential first step towards these goals is to parse these 200 million structures into constituent domains with high accuracy.

Existing protein domain boundary prediction techniques can be broadly classified into two categories: sequence-based approaches and structure-based approaches. As expected, approaches utilizing structural input outperform those relying solely on sequence information (6). With the advent of high-quality predicted structures from AlphaFold2 (7), obtaining a 3D structure is no longer a significant constraint. To integrate the 200 million AlphaFold models into protein domain databases, it is logical to exploit the predicted structure as an input for enhancing domain boundary prediction. Historically, most structure-based approaches have used unsupervised, heuristic algorithms applied to contact maps or pairwise residue distances. These approaches are grounded in the physical intuition that the density of contacts is higher within domains than between domains (8; 9; 10; 11; 12; 13). Although unsupervised methods can be effective, it is challenging to hand-design a heuristic that encompasses all cases. Other methods (14; 15) augment the unsupervised approach with the ability to match against a library of known domains, using sequence or structure comparisons. For example, DPAM (Domain Parser for AlphaFold Models) (15) uses a fixed formula to assess prospective splits as a function of three inputs: inter-residue distance, AlphaFold’s Predicted Aligned Error (PAE) and predicted domain co-membership via matching against a library of known domains. Comparison-based methods are well suited to segmenting proteins containing only known domains but may underperform on proteins containing domains which are not easily recognised by comparison tools or which are not included in existing databases.

The growth of protein structure databases complete with domain annotations presents an opportunity to instead recast the domain segmentation problem as a supervised learning task. Deep learning models have the potential to capture complex structural relationships and exploit these to achieve higher accuracy than heuristic unsupervised methods. Previously proposed supervised domain segmentation methods have mostly relied on sequence inputs and consequently struggled to match the performance of unsupervised methods which segment a known or predicted structure directly (6). Other supervised approaches have relied on a per-residue boundary classification approach (6; 16; 17). Somewhat similar to our proposed approach, Eguchi et al. train a convolutional neural network (CNN), herein EguchiCNN, to do image-segmentation on protein structures represented by 2D distance maps (18). The EguchiCNN architecture was primarily designed for the more specific task of classifying domain regions into one of the 38 CATH architectures. However, part of the pipeline includes a domain segmentation predictor which uses the same CNN architecture as their architecture classification model. This approach treats the domain segmentation problem as a multiclass classification of residues where each domain constitutes a separate class. Limitations of this approach are that it can only handle proteins up to 512 residues in length and it can only detect a maximum of 8 domains. A recent supervised approach called Merizo (19) uses a transformer architecture with invariant point attention to directly cluster residues into domains based on both sequence and structure inputs. This method was shown to perform better than UniDoc (13), SWORD (10), DeepDom (17) and EguchiCNN (18). However, Merizo was not trained on any single-domain CATH proteins, as such we find that it tends to over-split single-domain proteins (Section 2: Results).

In this work, we introduce Chainsaw, a supervised learning approach to protein domain segmentation. Instead of predicting domain boundaries directly or considering each domain as a separate class, Chainsaw relies on a 2D convolutional neural network trained to estimate the probability that pairs of residues belong in the same domain. Domain boundaries are derived from these pairwise co-membership probabilities using a greedy algorithm that searches for the most likely assignment of residues to domains given the predicted probabilities (Section 4.4). Formulating the supervised learning problem as a classification task at the level of pairs of residues rather than as a boundary prediction task at the level of individual residues has three notable advantages. Firstly, it makes the prediction of discontinuous domains more straightforward. Second, it sets no limit on the number of domains that can be predicted. Finally, it improves the class imbalance problems associated with residue classification. Unlike methods such as EguchiCNN, Chainsaw can handle inputs of any size without cropping or padding. We show that Chainsaw achieves better domain parsing accuracy when compared with supervised methods Merizo and EguchiCNN as well as other leading unsupervised structure-based domain parsers (UniDoc, PUU and SWORD) on held-out test sets of domain annotations from CATH and the Critical Assessment of Structure Prediction (CASP) competitions. We further evaluate Chainsaw on a random sample of AlphaFold models and find fewer domain prediction errors than the next best method. In a blind side-by-side human comparison of 200 AlphaFold models, we find the Chainsaw domain parsing to be preferable to UniDoc in roughly twice as many cases. Finally, we show that Chainsaw combined with Foldseek can be used to infer functional annotations in previously uncharacterized proteins.

## 2 Results

### 2.1 Protein domain segmentation with fully convolutional neural networks

The curated domain annotations in databases such as CATH provide a rich source of signal for training deep learning methods to segment protein structures into their constituent domains. We therefore first considered how to exploit such annotations as training data by formulating domain segmentation as a supervised learning problem. In particular, we sought to avoid some of the problems with training a network to solve a residue-level binary classification task of identifying residues at the boundary between domains. These problems include handling of discontinuous domains, severe class imbalance of boundary to non-boundary residues and sensitivity to small changes in the boundary. The latter is a problem given that in many cases multiple adjacent residues could equally be considered to be the ‘correct’ boundary. In addition, for training a neural network the domain label target representation should have some desirable properties such as being invariant to indexing or ordering of the domains and the dimensionality of the label should not depend on the number of domains. Our solution is to define a classification task over pairs of residues, from which domain assignments for individual residues can be recovered. To derive this pairwise task, we start by observing that the domain assignments of a protein of length *L* can be uniquely represented by an *L* × *L* matrix **A**, whose entries *a*_*i j*_ are 1 if residues i and j are in the same domain and 0 otherwise. For domains with residues that are continuous in sequence, this will result in an adjacency matrix that takes the form of a block-diagonal matrix, (see Figure 2a). Domains which are discontinuous in sequence result in blocks occurring in the off-diagonal; (see Figure 2b). Given the matrix **A**, residue-level domain assignments can be recovered by interpreting the matrix **A** as the adjacency matrix of a graph and partitioning it into a set of *K* connected components, where *K* is the number of domains. Importantly, this procedure works identically for domains that are continuous or discontinuous in sequence. Therefore, supposing we were able to produce a perfect predictor of the matrix **A** of pairwise domain co-membership between residues in a given structure, we could then unambiguously read off residue-level domain assignments for residues in either continuous or discontinuous domains. As a training label for a neural network, this representation has the advantage of being permutation invariant with respect to domain labels (there is no indexing suggesting an ordering of the domains) and the dimensionality is determined by the number of residues alone rather than being dependent on the number of domains.

**Figure 1.**
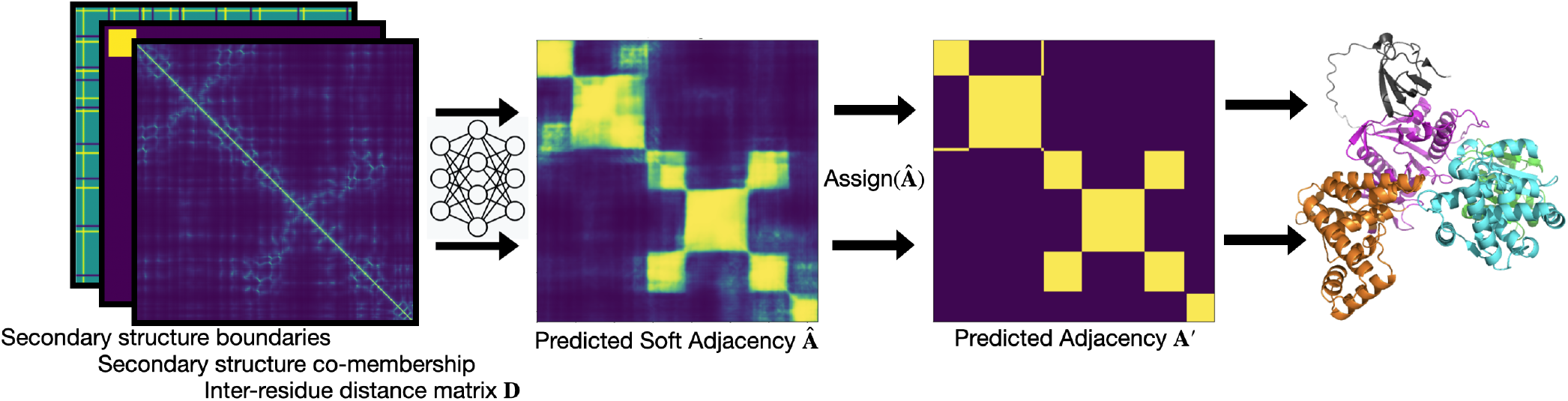
Chainsaw method overview.

**Figure 2.**
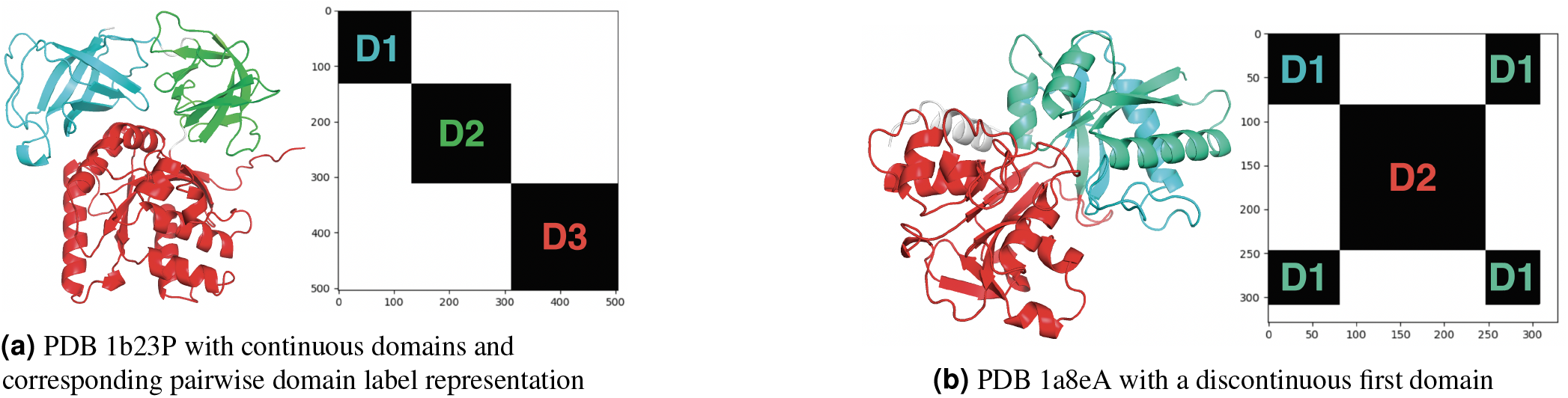
Domain assignments represented with binary pairwise co-membership matrices.

The main component of Chainsaw is a convolutional neural network (Figure 3) that is trained to predict the matrix **A** for a given input structure, by predicting whether each pair of residues belongs in the same domain. The network takes as input a set of pairwise representations derived from the protein’s 3D structure, consisting of the pairwise *α*-carbon residue distances and predicted secondary structure segments (Section 4.2), and outputs estimated probabilities that pairs of residues belong in the same domain. The final domain assignments are then determined by a search algorithm that maximises the likelihood of the assignments under the predicted pairwise co-membership probabilities (Section 4.4). As training data, we obtained a set of PDB structures with associated domain annotations from CATH, which were converted into the corresponding matrices **A** to serve as targets in a binary classification task for residue pairs (Methods).

**Figure 3.**
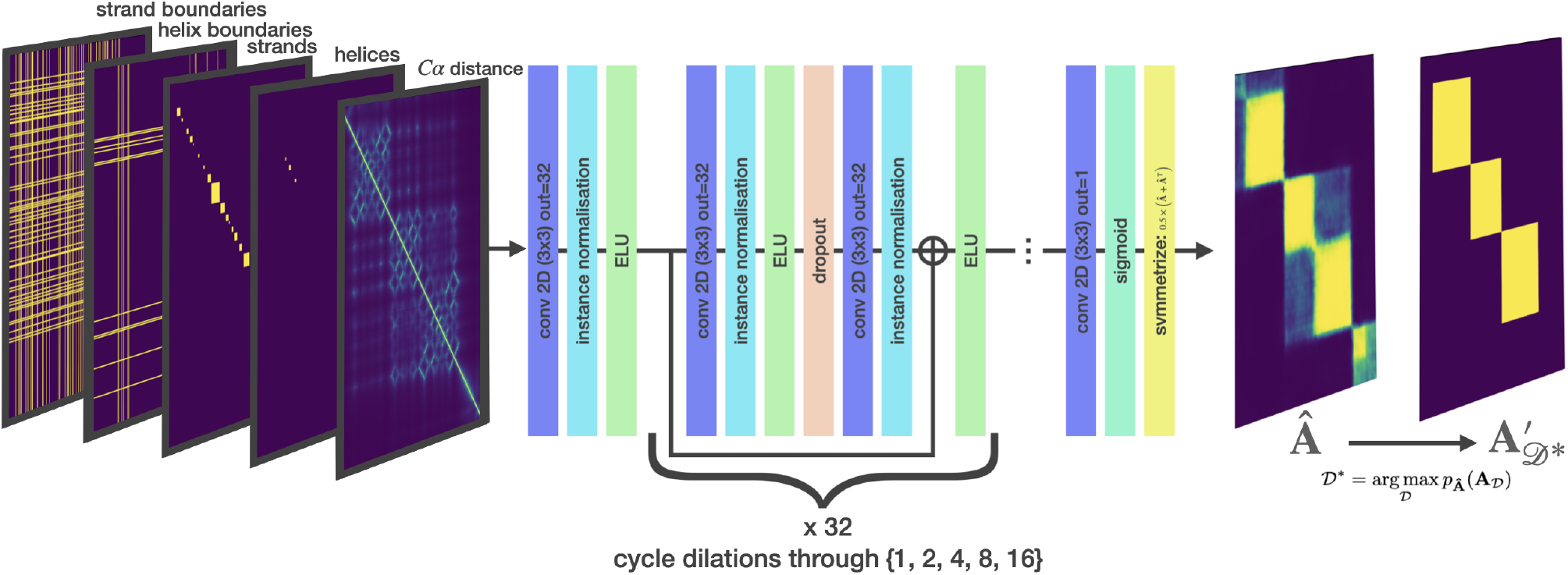
Chainsaw network diagram.

### 2.2 Chainsaw performance on experimental PDB structures

We benchmarked Chainsaw against three unsupervised methods, UniDoc, PUU and SWORD and two supervised methods, Merizo and EguchiCNN. Both Merizo and UniDoc were previously shown (19) to significantly outperform previous supervised learning methods EguchiCNN (18) and DeepDom (17). It should be noted that EguchiCNN was trained for the more specific task of classifying domains into superfamilies with generic segmentation as a pre-processing step and DeepDom is a structure-free predictor. We constructed a benchmark dataset of 1365 protein chains from the PDB using CATH domain annotations as ground truth. Our test dataset was constructed to ensure an equal balance of single and multi-domain proteins, this is motivated by findings in our other work (20) where we find that applying structure-based domain parsers to the AFDB suggests the following proportions: 3% zero-domain, 42% single-domain and 55% multi-domain. Using our test dataset we first compare Chainsaw against two unsupervised methods, UniDoc and PUU and one supervised method, EguchiCNN (Figure 4a). We note that EguchiCNN may have been trained on some proteins from our test set which could artificially inflate its performance. We cannot present results for SWORD on this benchmark as it failed to output results on a significant proportion of the test set. We assess domain prediction accuracy via three metrics: the intersection-over-union (IoU) (19), the proportion of correctly parsed domains (domain-level IoU >= 0.8) and the domain boundary distance score (21). Calculation details for all metrics are provided in Section A. Chainsaw achieves an average intersection-over-union (IoU) score of 0.88 vs 0.84 for the next-best competitor method UniDoc (Figure 4a). If we restrict our attention to multi-domain proteins only, the performance gap increases, with Chainsaw scoring 0.83 versus 0.76 for UniDoc. To compare against Merizo, we create a subset of the test data for which there is no domain in a CATH superfamily that also occurs in the Merizo training data (n=208). On this subset, Chainsaw achieves an average IoU of 0.91 vs 0.83 for Merizo (figure 4b). On this dataset, we find Merizo over-splits single domain structures in 28% of cases versus 10% for Chainsaw, plausibly because Merizo was trained solely on multi-domain proteins in CATH. We note that if we only compare methods on multidomain proteins the performance of Chainsaw and Merizo is the same, though the small sample size (n=52) limits the power of our analysis. To further test whether the performance differences between Chainsaw and Merizo can be attributed entirely to differences in the training data, we separately trained a Chainsaw model from scratch using the Merizo training data and evaluated it on the Merizo test data (which only contains multi-domain proteins). Following this approach we observe that Chainsaw still outperforms Merizo, suggesting that the differences are not solely driven by the difference in training data (figure 4c). As a final test, to see how Chainsaw performs on domain annotations which are not from CATH, we also evaluated a model on domain annotations from CASP6 (n=63) and find that the Chainsaw model still outperforms UniDoc, PUU, SWORD2 and Merizo (Supplementary Table 3).

**Figure 4.**
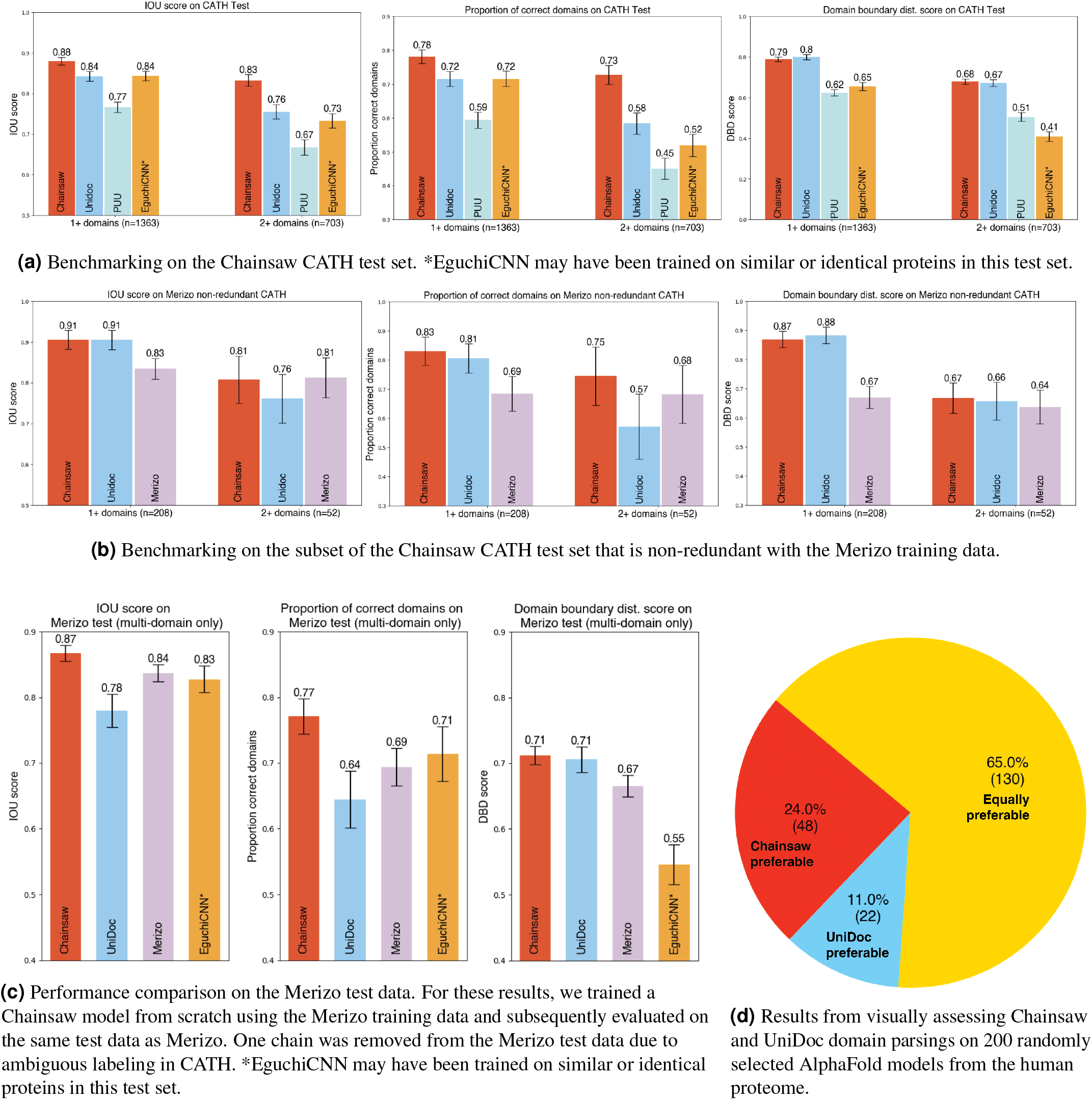
Assessing performance on CATH annotated PDB structures and AlphaFold models. Bar plots show 95% CI.

### 2.3 Chainsaw performance on AlphaFold models

We sought to evaluate Chainsaw’s performance on predicted structures in the AlphaFold database (AFDB). This is challenging because we lack a ground-truth domain assignment on AlphaFold models. One approach is to map CATH annotations from PDBs to AlphaFold models with matching sequences. Following this approach we observed no significant change in the performance of Chainsaw when predicting on AlphaFold models as opposed to their corresponding PDB files (Supplementary Table 2). However, this evaluation approach only considers AFDB structures in the PDB and are, therefore, typically well-modelled. In general, AFDB structures are notably different from experimentally resolved structures in the PDB. The most significant difference concerning domain parsing is the presence of long segments of residues with no apparent secondary structure (see figures 5b and 5h for examples). To evaluate performance on AlphaFold models, a random sample of 200 human protein structures was taken from the AlphaFold database. A naive sampling would have resulted in relatively few large proteins, so to mitigate this, we sampled structures equally from binned protein lengths to ensure representation of all sizes. Chainsaw and UniDoc predictions were assessed visually in a blind side-by-side comparison to look for domain segmentation faults. Faults considered were: under-splitting (see figures 5a, 5b, 5c, 5d), over-splitting (see figures 5e, 5f, 5i), incorrect boundaries (see figures 5g, 5c), missing domains, and falsely identifying domains (see figure 5h). UniDoc cannot predict non-domain residues. This would result in very poor performance when run on AlphaFold models which contain many poorly modelled segments with no predicted secondary structure. To improve UniDoc’s predictions, residues with low predicted confidence by AlphaFold (predicted local distance difference test < 70) were removed from the domain predictions. In our evaluation protocol evaluators compare the number and severity of segmentation faults to judge which segmentation is preferable. We also allow the option to judge the domain parsings of equal quality. We find that the Chainsaw domain parsing was preferable in 24% of cases, UniDoc was preferable in 11% of cases and both parsings were of equal quality in 65% of cases (Figure 4d). Figure 5 showcases a selection of these judgements. Judgements and domain parsing images for all 200 assessed AlphaFold models are shown in the Supplementary Material.

**Figure 5.**
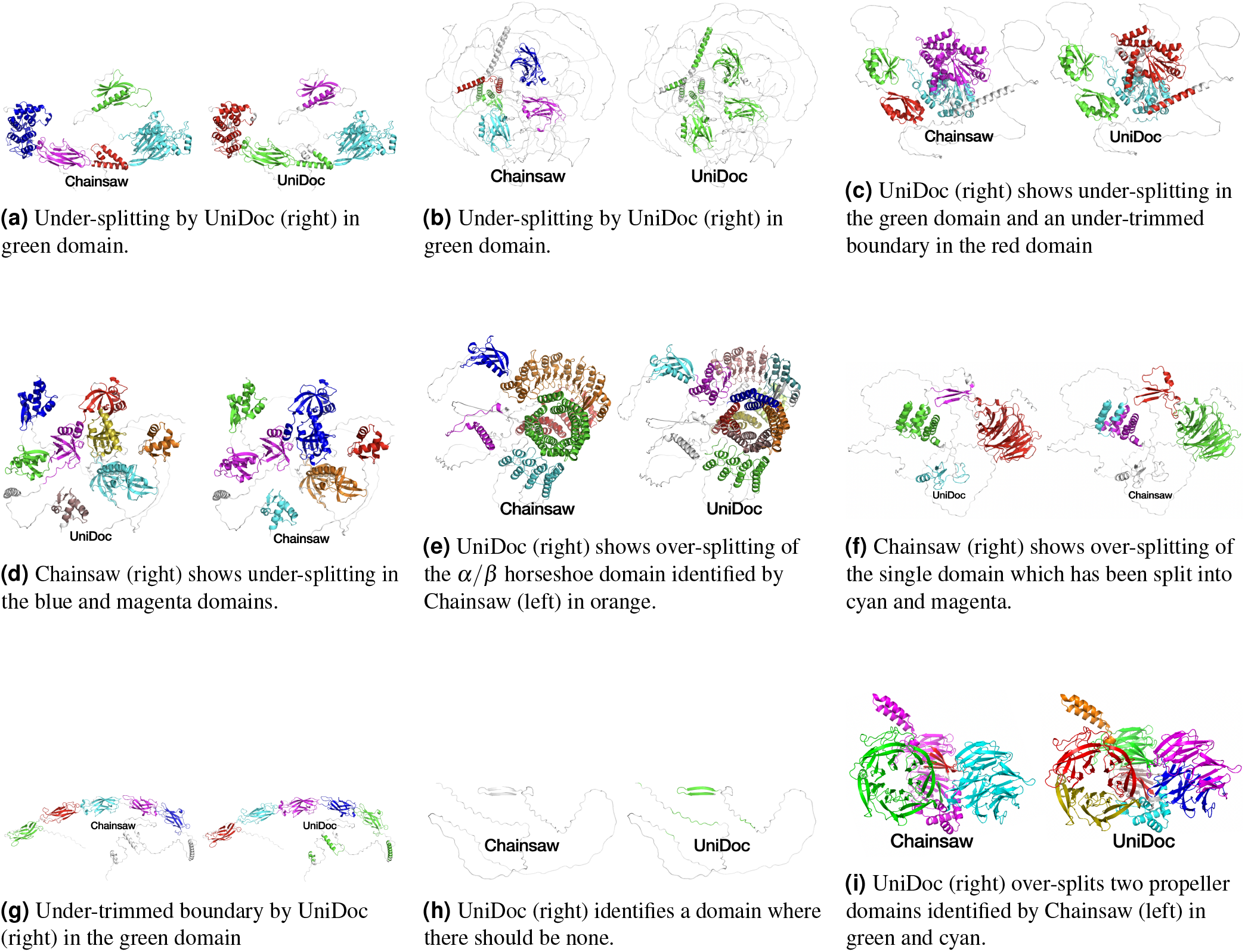
A selection of judgements from the blind comparison of UniDoc and Chainsaw on 200 AlphaFold human structures. Domain segmentations and judgements for all 200 proteins can be found in the supplementary material.

### 2.4 Inference times

Chainsaw’s inference time is 0.6s on CPU (0.2s on GPU) (Table 1). The inference time for each model was calculated by measuring prediction times on a set of PDBs where the mean sequence length was 164 residues and the max sequence length was 500 residues. The CPU times were generated on an 8-core M1 MacBook Pro. The GPU times were generated on an Nvidia RTX 4090.

**Table 1.**
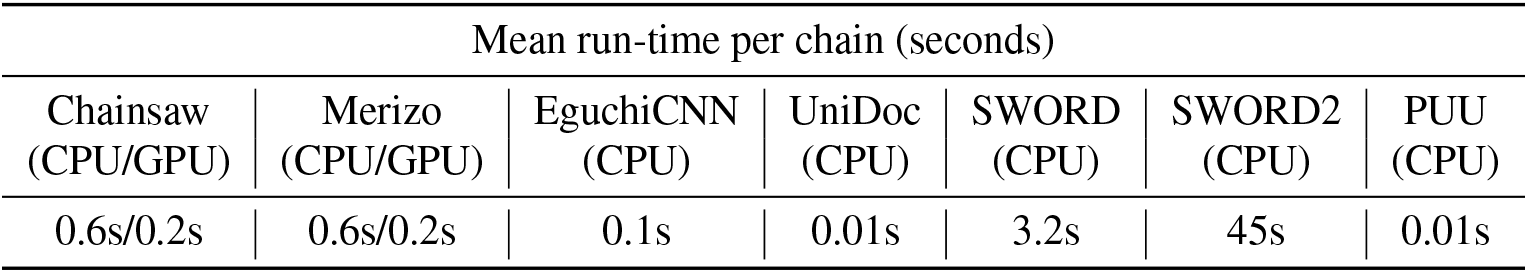
Model inference times.

| Mean run-time per chain (seconds) |  |  |  |  |  |  |
| --- | --- | --- | --- | --- | --- | --- |
| Chainsaw<br>(CPU/GPU) | Merizo<br>(CPU/GPU) | EguchiCNN<br>(CPU) | UniDoc<br>(CPU) | SWORD<br>(CPU) | SWORD2<br>(CPU) | PUU<br>(CPU) |
| 0.6s/0.2s | 0.6s/0.2s | 0.1s | 0.01s | 3.2s | 45s | 0.01s |

**Table 2.** Comparing Chainsaw performance on AlphaFold models versus experimental structures from the PDB.

| Dataset | Chainsaw NDO Score |  |
| --- | --- | --- |
|  | on AlphaFold | on PDB |
| CATH n. dom. $\geq 1$ | 0.93 | 0.94 |
| CATH n. dom. $\geq 2$ | 0.91 | 0.90 |
| CATH n. dom $\geq 3$ | 0.94 | 0.95 |

**Table 3.** Comparison of methods CASP 6 dataset.

|  | Chainsaw | UniDoc | PUU | SWORD2 | Merizo |
| --- | --- | --- | --- | --- | --- |
| Intersection over union | **0.89** | 0.87 | 0.88 | 0.88 | 0.84 |
| Proportion correct domains | **0.78** | 0.74 | 0.72 | 0.77 | 0.69 |

### 2.5 Using Chainsaw predictions for downstream tasks

We employed Chainsaw to generate domain predictions for three downstream tasks. The first is identifying domains in the PDB that have not yet been annotated in the CATH protein domain database. The second is identifying domains in the AFDB that can be matched to representative domains found in CATH, finally, we show that Chainsaw can be used to infer novel functional annotations for previously uncharacterized proteins. In each of these cases, we first generate domain predictions using Chainsaw and subsequently use the Foldseek (22) structure and sequence matching algorithm to match predicted domains against a library of known CATH domains (clustered at 60% sequence identity). Figure 6 shows three examples where Chainsaw combined with FoldSeek can identify domains in PDB structures which appear to be good candidates to bring into the CATH domain database. For our second task, we generate Chainsaw domain predictions for the entire proteome of *Leishmania infantum*, a parasite responsible for the neglected tropical disease infantile visceral leishmaniasis. We find that around 16% of Chainsaw predicted domains in this proteome can be matched with high confidence (Foldseek value < 1e-10) to an existing CATH S60 domain. This amounts to 2,567 predicted domains where we can infer homology to existing domains. Following the same procedure for UniDoc we note that the UniDoc+Foldseek pipeline matches a similar number of predicted domains (2,451), however 28% of Chainsaw+Foldseek domain matches are not discovered using the UniDoc+Foldseek pipeline and 24% of UniDoc+Foldseek domain matches are not discovered by the Chainsaw+Foldseek pipeline. A ‘domain match’ is defined here as identifying the same CATH S60 representative domain in a given AlphaFold model. This suggests that Chainsaw and UniDoc are complementary methods which can be combined to recover more correct domain parsings than either method individually. We note, however, that the overlap between methods would likely be greater if we matched against CATH domains clustered at 35% sequence identity or considered the top-k Foldseek matches instead of only the top one. For the final analysis, we show that the Chainsaw+Foldseek pipeline can infer functional annotations for *Leishmania infantum* proteins that are uncharacterized. We start by considering the subset of the *Leishmania infantum* proteome where UniProt lists the protein as ‘uncharacterized’ and there are no Gene Ontology (GO) annotations (23) associated with the protein. This amounts to 3280 out of 7924 *Leishmania infantum* proteins. Of these we find 413 proteins which have a Chainsaw predicted domain which matches against a CATH S60 representative domain with a Foldseek e-value of less than 1e-5. Of these matched proteins, 396 are matched to a CATH S60 domain with a functional annotation from Pfam (24) or GO. Four of these are shown as case studies in Figure 7. We note that this technique can detect structural homology in multi-domain proteins where the sequence identity is frequently less than 15% and for each of the showcased examples (Figure 7) we checked that there were no Pfam, CATH FunFam or Gene3D matches returned when running the sequences through InterPro Scan.

**Figure 6.**
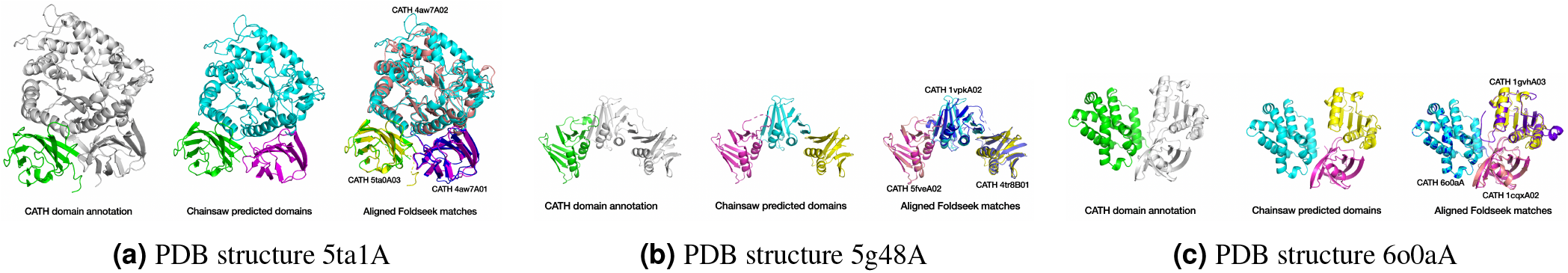
Example PDB structures where Chainsaw combined with FoldSeek can identify two additional domains which have not yet been annotated by CATH. In each case, the single CATH-identified domain is shown on the left, Chainsaw’s domain prediction is shown in the middle and the matched CATH domains aligned with the Chainsaw predictions are shown on the right.

**Figure 7.**
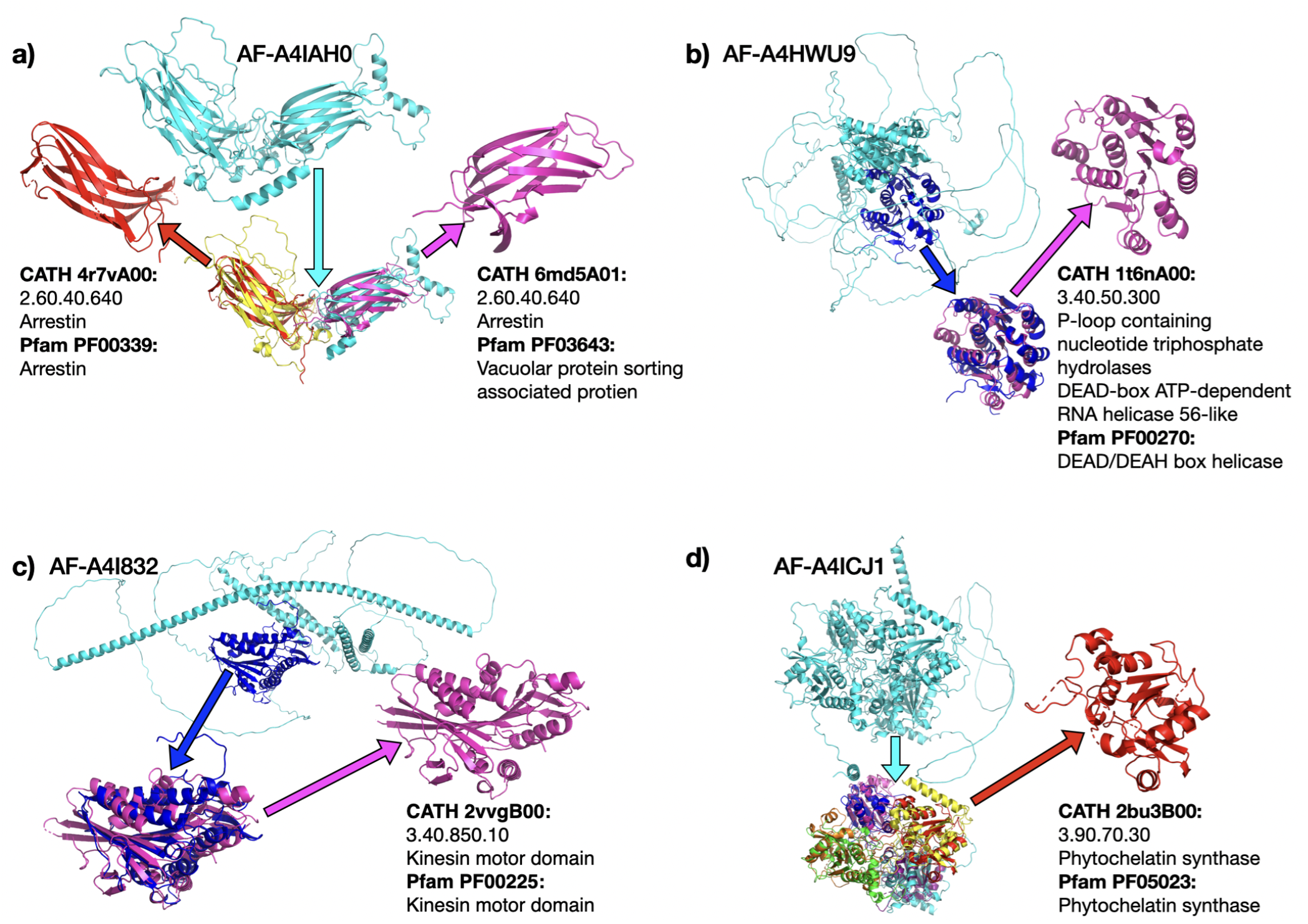
Uncharacterized proteins (no Pfam or Go annotations) from the *Leishmania infantum* proteome were parsed with Chainsaw. The predicted domains were subsequently searched against the CATH S60 domain representative structures using Foldseek. We show four examples where we can infer novel functional annotations via structural homology with representative CATH domains. Sequence identity for the matches above ranges from 11 to 16 per cent, which indicates why these homologous relationships were not detected with sequence-only methods. Figure **d** shows that Chainsaw correctly parses the structure into four phytochelatin synthase domain repeats. This protein is common to multiple pathogenic organisms and has been considered a potential drug target due to the fact it has no human homolog (25).

## 3 Discussion

This study presents Chainsaw, a supervised learning approach to protein domain prediction, we leverage a residual convolutional neural network to estimate the probability that pairs of residues are in the same domain and combine this with an algorithm for converting the pairwise probabilities into domain assignments. Our approach outperforms state-of-the-art structure-based methods on both annotated PDB structures and AlphaFold models. Nonetheless, in our analysis of performance on AlphaFold models we observe a significant proportion (11%) of cases where UniDoc domain parsings are preferable to Chainsaw and a majority of cases (65%) where the two predictions are of equal quality. In cases where the parsing is of equal quality a significant proportion of these are proteins that could be parsed in multiple ways, with alternative segmentations by each method appearing equally valid. For these reasons, we conclude that a sensible strategy for domain identification involves using an ensemble of domain prediction methods. This approach enables confidence measures derived from model consensus and a diversity of possible predictions. The possibility of multiple valid domain segmentations within a single structure motivates future work to extract multiple predictions from Chainsaw. We observe that the Chainsaw neural network confidence score is correlated with prediction accuracy (Section 4.5) and we see some hints that uncertainty in the output is indicative of alternative valid assignments. This insight opens up the potential for adapting the domain assignment algorithm to yield multiple assignments from a single network prediction. An advantage of Chainsaw when compared with models such as DPAM (15) is the lack of dependencies on databases of known domains. We demonstrate a proof-of-concept showing that domain prediction methods can be combined with structure and sequence matching algorithms to systematically identify domains and homology relationships in large databases of predicted structures such as AFDB. We further show that this approach can detect homologs where sequence-based methods cannot (Figure 7). A natural extension of this work is to apply these techniques at scale and develop approaches for detecting novel domains. It is important to note that while we have trained Chainsaw solely using domain annotations from the CATH database, there are several other widely used domain classification resources available, including SCOP (26) Pfam (24) and ECOD (4). The benchmarking we conducted primarily focuses on CATH annotated domains however, we are encouraged to see that Chainsaw has good performance on domain annotations from CASP, as well as on un-annotated structures in the AlphaFold database.

## 4 Methods

### 4.1 Datasets

Following a similar approach to Merizo (19), we generated a train-validation-test split on CATH annotated PDB files ensuring that no CATH superfamily is represented in more than one of the splits. We only include PDBs composed of domains belonging to CATH classes 1, 2 and 3. We note that the CATH superfamily is one level stricter than the S35 (35% sequence identity) clusters within the CATH hierarchy (27). We opted not to use the same splits as Merizo, as these splits only contained multi-domain chains and we know that approximately 45% of chains in the AlphaFold Database have one or zero domains (20). After splitting the CATH superfamily codes into train, test and validation we represent each PDB chain by a tuple which contains all of the CATH S35 codes of its constituent domains. For each unique tuple of CATH S35 codes we take one PDB chain to represent that cluster in the validation and test sets. PDB chains which contained irregular amino acids or were missing *α*-carbon atoms were removed, two additional chains 2v495 and 3vkgB were removed from the test set due to incorrect and incomplete CATH annotations. For the training data we take one representative PDB file for each tuple of CATH S95 (95% sequence identity) clusters. To account for the additional training data redundancy introduced by this choice, during training we sample chains to train on in a redundancy-aware manner, by making use of CATH’s sequence-identity-based clustering of domains. As such, one epoch is defined as one pass through all of the S60 sequence clusters (clustered at 60% sequence identity). We additionally experiment with varying the relative frequency with which single and multi-domain chains are sampled as training datapoints. Our final approach sampled multi-domain proteins with probability 0.65 and single domain with probability 0.35. Hyper-parameter selection was based on performance on the validation set, final performance is reported on the non-redundant test set. To show that our method is not overfitting to CATH assignments and to test that the approach generalizes to alternative domain assignments we evaluate an additional test set using the domain assignments from CASP 6 (28) (Supplementary Table 3).

### 4.2 Input features

The Chainsaw neural network takes a 3D structural representation of a protein, such as a PDB file. From the 3D structure, we generate five feature channels comprised of a residue pairwise distance matrix and four channels representing predicted secondary structure using STRIDE (29). The pairwise residue distance matrix is an *L* × *L* matrix **D** where element *d*_*i j*_ is the distance, in angstroms, between the *α*-carbon atoms of residues *i* and *j*. The predicted secondary structure is represented in two formats. The first is a co-membership matrix **C** where element *c*_*i j*_ is 1 if residues *i, j* are in the same secondary structure component, 0 otherwise. The second format indicates which residues occur at the start and end of secondary structure components with the first residue of a secondary structure component indicated with 1 and the last residue −1. Each of the secondary structure representations is instantiated independently for helices and strands resulting in four secondary structure feature channels in total.

### 4.3 Network architecture and training objective

We formulate the supervised learning problem as a 2D to 2D task: transforming the 2D input features into a pairwise probability matrix which expresses the probability that pairs of residues are in the same domain. As such, a fully convolutional architecture with skip connections is a natural choice. We use a modified version of the trRosetta architecture (30), a model originally developed for protein structure prediction. trRosetta is a residual network, whose blocks employ convolutional layers with progressively increasing dilation rates to achieve a wide receptive field (figure 3). We enforce a symmetric-output constraint by adding the transpose of the final layer to itself to give the final symmetrised **Â**. We truncate the trRosetta architecture to 31 blocks and reduce the number of filters from 64 to 32 but otherwise follow the original model in all details (30). The learning objective is to minimize the binary-cross entropy of the predicted residue pairwise domain co-membership matrix (which we call the soft adjacency matrix) and the true adjacency matrix. One advantage of this pairwise representation in both inputs and outputs lies in its SE(3) invariance, signifying that the representation remains unaltered under rigid transformations of the 3D structure, which consist of rotations, translations and reflections within the original coordinate space. An important theoretical motivation for using this representation is that a unique configuration of points in 3D space, up to rigid transformation, is fully specified by the 2D pairwise distance representation, (see theorem 1). To put this another way, the 2D representation captures all relevant spatial geometry while ignoring nuisance factors such as the arbitrary selection of the coordinate system that is used to represent the 3D structure.

### 4.4 Domain assignment algorithm

The output of the neural network is an *L* × *L* matrix **Â**, with entries *â*_*i j*_ representing the probability that residue *i* and residue *j* belong to the same domain. A further processing step is required to resolve the uncertainty in the neural network’s prediction to derive a final domain assignment. Let **A**_*𝒟*_ be the binary matrix associated with a given domain assignment *𝒟*. To identify the most likely domain assignment *𝒟* ^∗^ given a predicted set of co-membership probabilities **Â**, we seek to find the *𝒟* with maximum probability under **Â**,

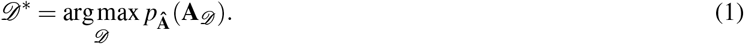

The probability of a domain assignment given the predicted values **Â** is given by the product of the entries

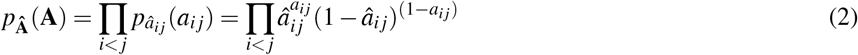

where each *â*_*i j*_ represents the probability *a*_*i j*_ = 1. The representation **A**_*𝒟*_ has useful properties as a label for supervising the neural network (Section 2.1), however, at the stage where we search for the optimal assignment it is preferable to work with a low-rank factorization of **A**_*𝒟*_, where each residue’s domain assignment is represented as a one-hot encoded vector. To perform the maximisation, we use a greedy algorithm, which exploits the fact that **A**_*𝒟*_ has a low-rank structure induced by the domain assignment *𝒟*. Let **v**_1_,…, **v**_*K*_ be a sequence of binary indicator vectors for each of the *K* (predicted) domains, let **V**_*D*_ be the matrix whose columns are the **v**_*i*_, hence **V**_*𝒟*_ is a binary matrix with dimensions *L* × *K*. The requirement that no residue can be assigned to more than one domain ensures that the rows of this matrix are *K*-dimensional one-hot vectors indicating domain assignments for each residue. Given this matrix of domain assignments, **V**_*𝒟*_, the elements *a*_*i j*_ of the adjacency matrix **A**_*𝒟*_ are generated as

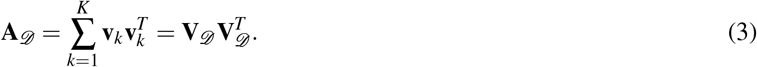

Thus our maximisation problem becomes to find a set of *K* vectors, such that the probability of domain assignment induced by the vectors is maximised

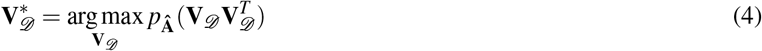

where the optimal number *K* of domains is itself unknown *a priori* and is therefore determined jointly with the domain assignments. The overall DomainAssigner(**Â**) procedure is defined in Algorithm 1. Note that the overall computation time is kept low by maximizing the log probability

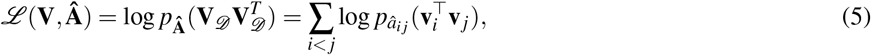

which decomposes into a sum of log probabilities over individual entries in the adjacency matrix. This means that the change in probability by changing a single residue *j* to be assigned to domain *k*,

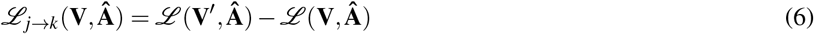

where **V**^′^ has been modified only in r ow *j*, can be computed in runtime which is linear with the number of residues.

Our domain assignment algorithm proceeds as follows. First **V**_*𝒟*_ is initialised as an *L* × *K*_*init*_ matrix of zeros, where *K*_*init*_ is an initialisation for the number of domains (*K*_*init*_ = 4). For each residue in turn, we score each of the K possible domain assignments using the score *p*_**Â**_ (**V**_*𝒟*_ **V**_*𝒟*_^*T*^), and assign the residue to the domain which produces the maximal score, as long as the maximal score is greater than the score under the current assignment. In case of ties, the first domain is selected. After a complete pass through the sequence, the process is iterated a number of times to allow for corrections in initial assignments. As soon as some residue is assigned to the *K*^*th*^ domain, we add an extra column of zeros, corresponding to an ‘overflow’ domain which can be subsequently assigned to. This allows the algorithm to predict an arbitrary number of domains. The procedure is summarised in Algorithm 1. We note that the algorithm is incentivised to predict the correct number of domains: given a perfect predictor of **A**, then for any ground truth adjacency matrix **A**_*D*_, all assignments other than the correct assignment are guaranteed to receive lower scores, including assignments which over- or under-predict the number of domains. Indeed for a perfect predictor, our algorithm is guaranteed to recover the correct domain assignment. Not all residues have to be assigned to a domain. The algorithm only makes an assignment if the score of the best possible domain assignment is greater than the score under a null assignment, in which the residue is not assigned to any domain. Since each residue is initialised with a null assignment (corresponding to a row of zeros in **V**), residues for which no better assignment is found will maintain a null assignment throughout the procedure.

#### Algorithm 1 DomainAssigner(Â)

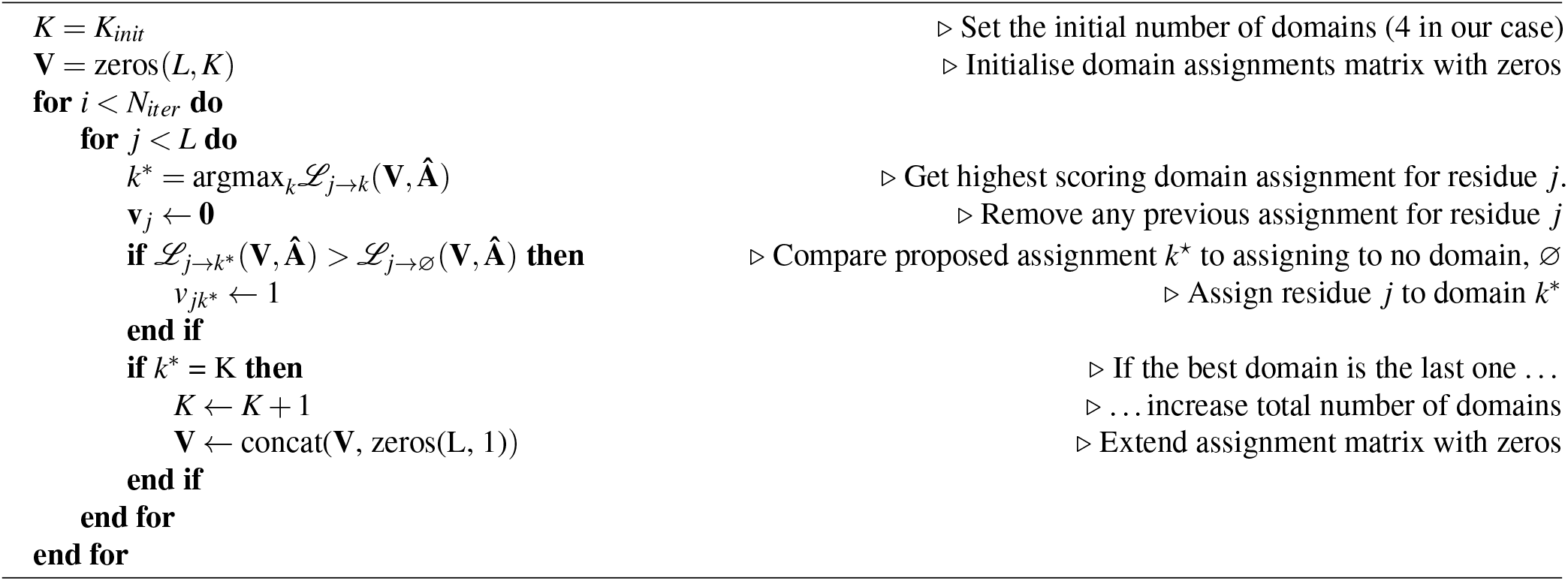

### 4.5 Uncertainty quantification

A natural approach to uncertainty quantification is to consider the output of the neural network **Â** as an *L* × *L* multivariate Bernoulli distribution. Then we can consider the output of the final assignment **A**^′^ as an observation from the **Â** distribution and calculate the likelihood (normalised by the number of residues). Figure 9 compares instances where the model has high confidence (figure 9a) with other cases where the model confidence is lower suggesting that alternative domain assignments may be valid (figure 9b).

**Figure 8.**
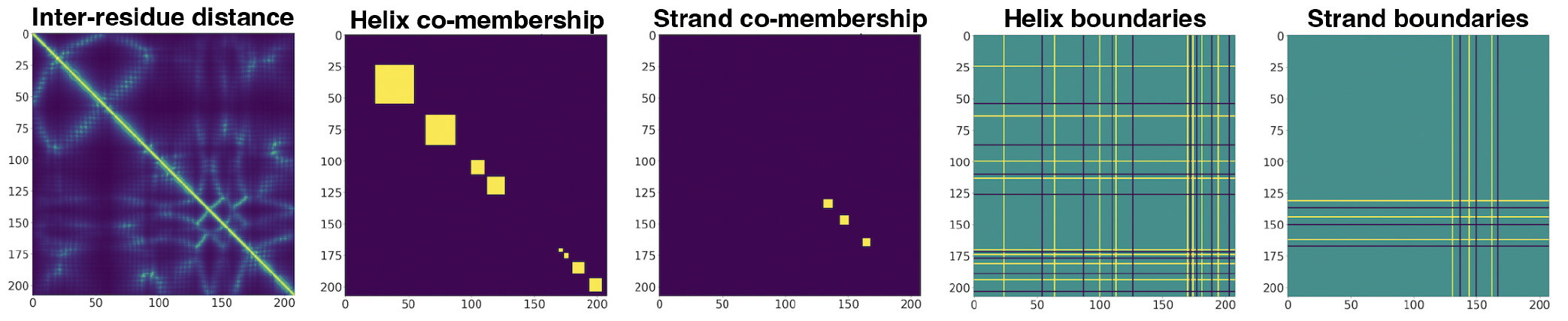
Input features for the Chainsaw neural network.

**Figure 9.**
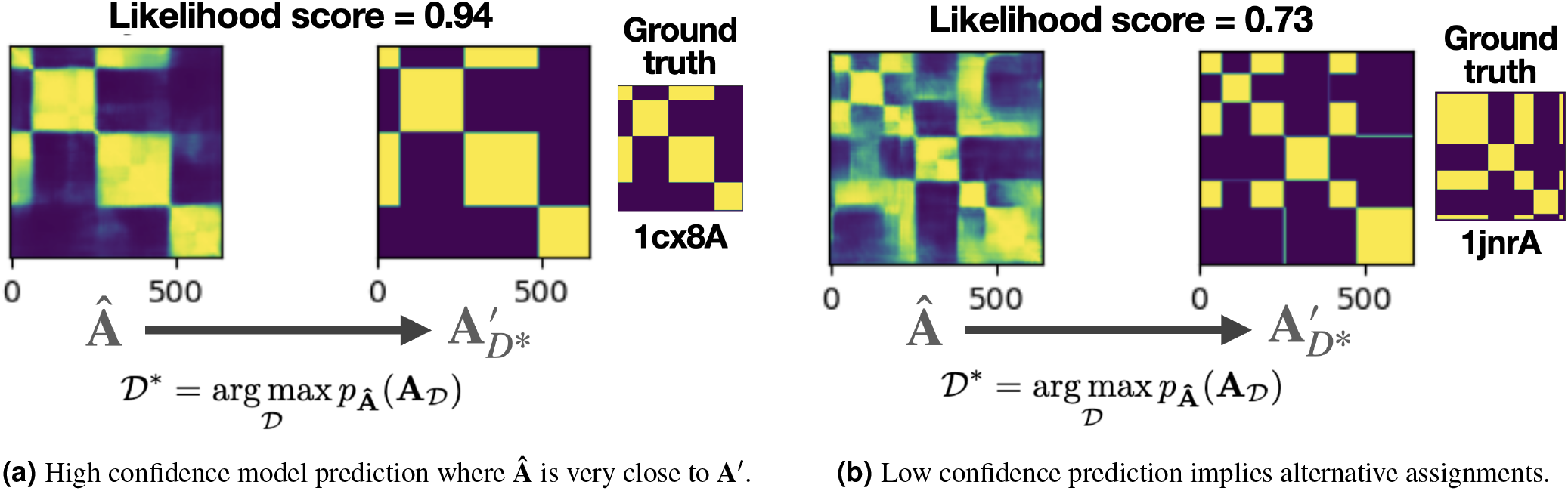
Chainsaw generates a confidence score which is the residue-averaged likelihood of **A**^′^ under **Â** we find that this confidence measure has a good correlation with ground-truth accuracy.

We observe that Chainsaw’s confidence score has a good correlation with the ground truth accuracy. The confidence score achieves a Spearman’s correlation score of 0.68 with the IoU score when measured on the Chainsaw CATH test set. On this test set the confidence score ranged from 0.51 to 1.0. Using a confidence score cutoff of 0.85 will increase the probability that a domain is predicted correctly (IoU score > 0.8) from 0.78 to 0.9 at the expense of introducing a 5% chance that a correctly predicted domain is discarded.

## Supporting information

Comparison of Chainsaw and UniDoc domain parsings on 200 AF models from human proteome

## 6 Acknowledgements

Jude Wells acknowledges the receipt of studentship awards from the Health Data Research UK-The Alan Turing Institute Wellcome PhD Programme in Health Data Science (Grant Ref: 218529/Z/19/Z). Alex Hawkins-Hooker was supported by the EPSRC Grant EP/S021566/1.

## 7 Supplementary material

### A Assessment metrics

#### A.1 Intersection over union

Following the approach of Merizo (19), for a given protein chain we compute the average intersection over union (IoU) between paired sets of predicted and ground-truth residues assigned to each domain. As a first step, each ground-truth domain is paired with a predicted domain such that the sum of all intersections over unions is maximised while respecting the following constraints: Each ground-truth domain (represented as a set of residue indices) *T*_*i*_ can have, at most, one paired predicted domain *P*_*i*_. Second, each predicted domain can be assigned at most once. No IoU is computed for the sets of residues that are labelled as, or predicted to be non-domain residues. To generate a final score for the whole chain each domain-level IoU is weighted by the number of residues in the ground-truth domain:

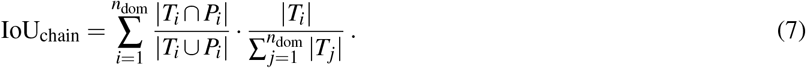

We additionally calculate the proportion of correctly parsed domains as the proportion of ground-truth domains where the domain-level IoU is 0.8 or greater.

#### A.2 Domain boundary distance score

The domain boundary distance score was introduced to assess domain boundary predictions in CASP 7. A detailed description of how the score is calculated is provided in (21). At a high level: each boundary is scored independently. Predicting within 1 residue of the true boundary scores 8 points, within 2 residues scores 7 and so on until the distance is 9 residues or more, at which point the score is 0. Each boundary score is divided by 8 so that scores per boundary are between 0 and 1. The final boundary distance score for the entire chain is then calculated as the sum of individual predicted boundary scores divided by the total number of domain boundaries. In order to ensure that over-prediction is penalized, the number of domain boundaries comes from the maximum number of domains in the target or the number of domains in the prediction (21).

### B Proof that pairwise 2D distances specifies 3D points up to isometry

#### Theorem 1.

*Let* **x**_0_,…, **x**_*n*_ *and* **y**_0_,…, **y**_*n*_ *be tuples representing points in the Euclidean space* 𝔼^3^. *Assume that for every pair of indices i, j* ∈ {0,…, *n*}, *we have*

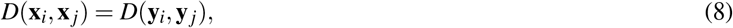

*where D denotes the distance function on* 𝔼^3^:

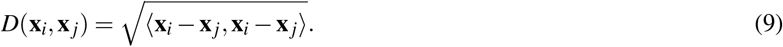

*and* ⟨·, ·⟩ *denotes the inner-product of two vectors. Then, there exists a unique bijective isometry g*: 𝔼^3^ → 𝔼^3^ *such that g*(**x**_*i*_) = **y**_*i*_ *for each i*.

*Proof*. **Step 1: Translation to Origin**. First, choose **x**_0_ and **y**_0_ and translate all points **x**_0_,…, **x**_*n*_ and **y**_0_,…, **y**_*n*_ in 𝔼^3^ such that **x**_0_ = **y**_0_ = **0**. These translations preserve distances, so

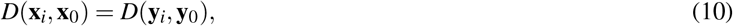

and therefore:

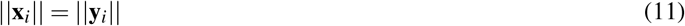

where || · || denotes the L2-norm of a vector.

**Step 2: Equality of Distance Implies Equality of Inner Product**.

From the definition of the distance function, we have

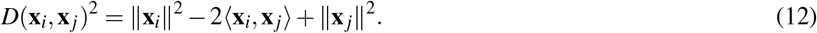

Using Equations (8) and (11), we deduce

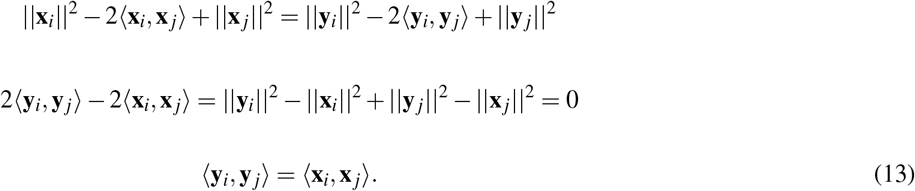

**Step 3: Identifying Basis Vectors**.

Without loss of generality, choose vectors **x**_1_, **x**_2_, and **x**_3_ that form a basis for 𝔼^3^.

#### Lemma 1.

*The Gram matrix of a set of vectors is non-singular if and only if the set of vectors are linearly independent*.

Let **G** be the 3 × 3 Gram matrix of **x**_1_, **x**_2_, **x**_3_, defined by

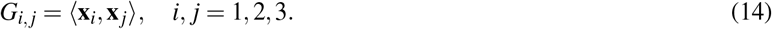

Since **y**_1_, **y**_2_, **y**_3_ share the same Gram matrix **G** (due to eq. 13), they also form a basis for 𝔼^3^.

#### Lemma 2.

*If two sets of basis vectors have the same Gram matrix, there exists an isometry that maps one set to the other*.

Let **T** be a (3 × 3) matrix which applies the orthogonal transformation *g*, mapping **x**_1_, **x**_2_, **x**_3_ to **y**_1_, **y**_2_, **y**_3_.

**Step 4: Extension to All Vectors**.

Any vector **x** in 𝔼^3^ can be expressed as a linear combination of our basis vectors:

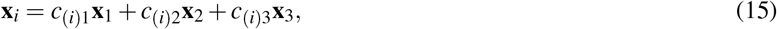

for some *c*_(*i*)1_, *c*_(*i*)2_, *c*_(*i*)3_ ∈ ℝ.

A unique solution for the values *c*_(*i*)1_, *c*_(*i*)2_, *c*_(*i*)3_ can be obtained using only the inner products of **x**_1_, **x**_2_, **x**_3_ with **x**_*i*_ We can write this as a system of linear equations in matrix form **Gc**_(*i*)_ = **b**, where

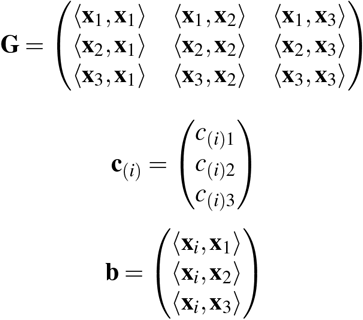

Since **x**_1_, **x**_2_, **x**_3_ form a basis, the matrix **G** is invertible. Therefore, the system of equations has a unique solution for **c**_(*i*)_. Given that ⟨**y**_*i*_, **y** _*j*_⟩ = ⟨**x**_*i*_, **x** _*j*_⟩ we see that matrix **G** and vector **b** will give rise to the same solutions for *c*_(*i*)1_, *c*_(*i*)2_, *c*_(*i*)3_ satisfying the equation

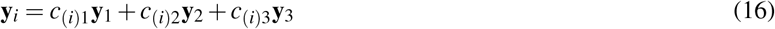

Finally, we note that the orthogonal transformation **T** maps every vector **x**_*i*_ to **y**_*i*_

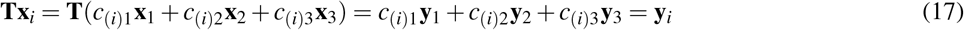

□
which completes the proof.

### C NDO score

The NDO score is a score with a maximum of 1 which represents the proportion of residues that have been assigned to the correct domain. To understand the calculation, the score can be decomposed into scores for each predicted domain and scores for each true domain. Each predicted domain 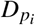 gets an un-normalised score 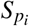 which is the number of residues in the maximum intersection over all intersections with the ground-truth domains 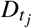 minus the sum of the size of all its other intersections with ground-truth domains (all those that do not include the maximum intersection):

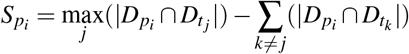

Similarly, each ground-truth domain, 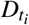, gets an un-normalised score, 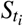, which is the number of residues in the maximum intersection over all intersections with the predicted domains minus the sum of all its other intersections with predicted domains:

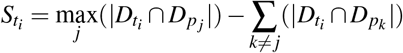

The individual unnormalised predicted domain scores and ground truth domain scores are summed together, divided by two (to account for counting residues in both the true domain scores and predicted domain scores), before being divided by the maximum unnormalised score 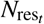 which equals the number of residues assigned to domains in the ground-truth assignment:

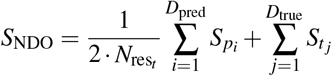

### D Additional results

#### D.1 Predicting CATH domain annotations mapped to AlphaFold models

When predicting domain boundaries on AlphaFold models as opposed to PDB structures we do not have a ground-truth set of labels. To overcome this problem we use SIFTS (31) to map PDB structures to their corresponding predicted structures in the AFDB. This enables us to map a subset of the CATH test set domain annotations onto AlphaFold models. Using this approach we compare Chainsaw’s performance on 1039 samples from the CATH test set. Table 2 shows that we do not see any significant decrease in performance when predicting on AlphaFold models as opposed to PDB structures. However, it is important to note that this subset of AlphaFold models is not representative of the AFDB as a whole because it only contains sequences that have experimental PDB structures and have been annotated by CATH. In contrast with the AFDB overall we observe that this subset of AlphaFold models has better modelled structure and contains few proteins with long regions of non-secondary structure residues.

#### D.2 Performance on CASP 6 domain annotations

To evaluate Chainsaw on CASP 6, we trained a separate model from scratch to ensure no homologous domains were in the training data.

#### D.3 Ablation analysis: secondary structure features

**Figure 10.**
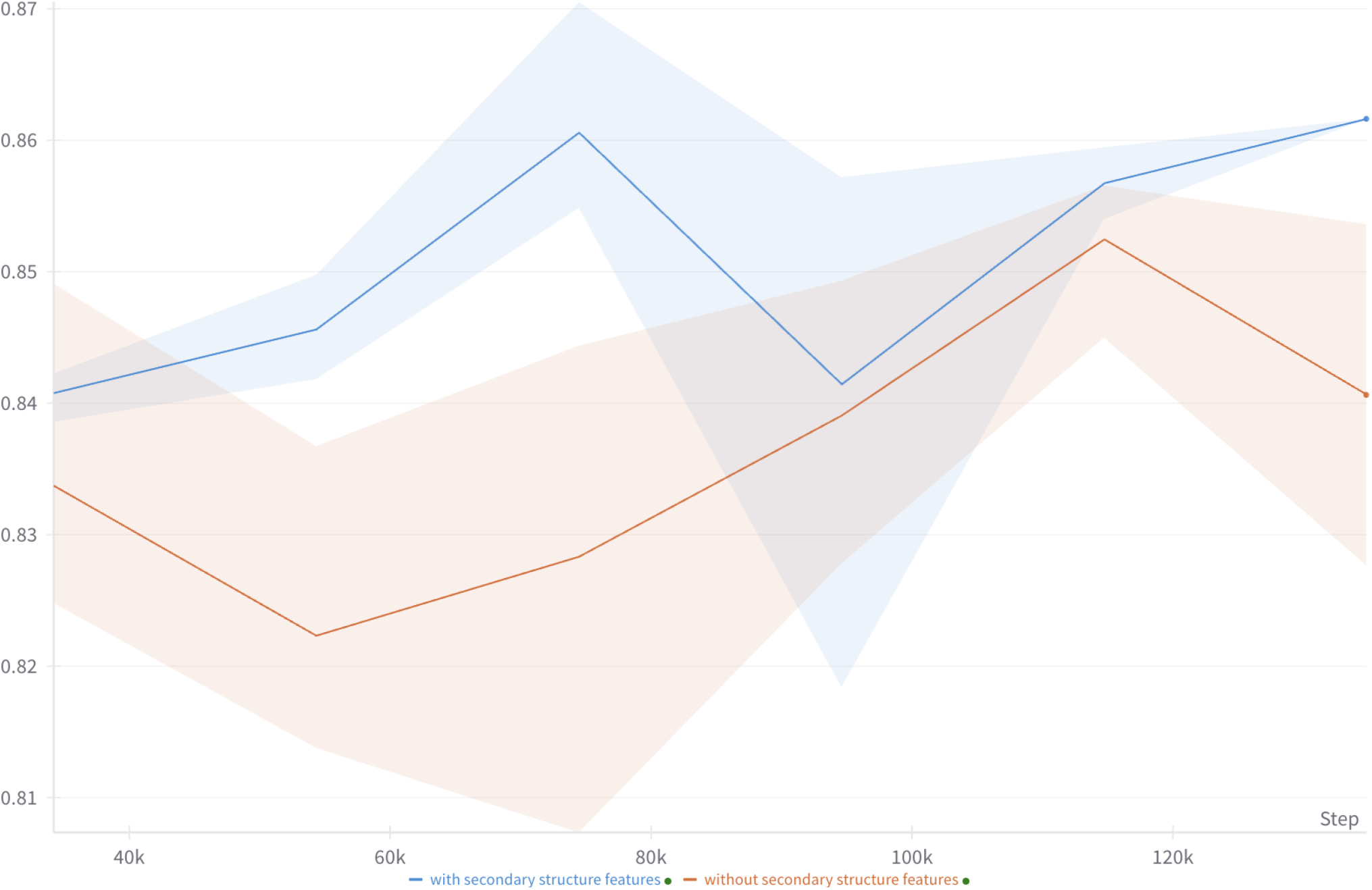
Results from training six Chainsaw models, three with secondary structure features included and three without (distance matrix only). We show the grouped mean IoU score on the validation data. The shaded range covers the minimum and maximum values at each epoch.

#### D.4 Ablation analysis: alpha distances versus beta distances

**Figure 11.**
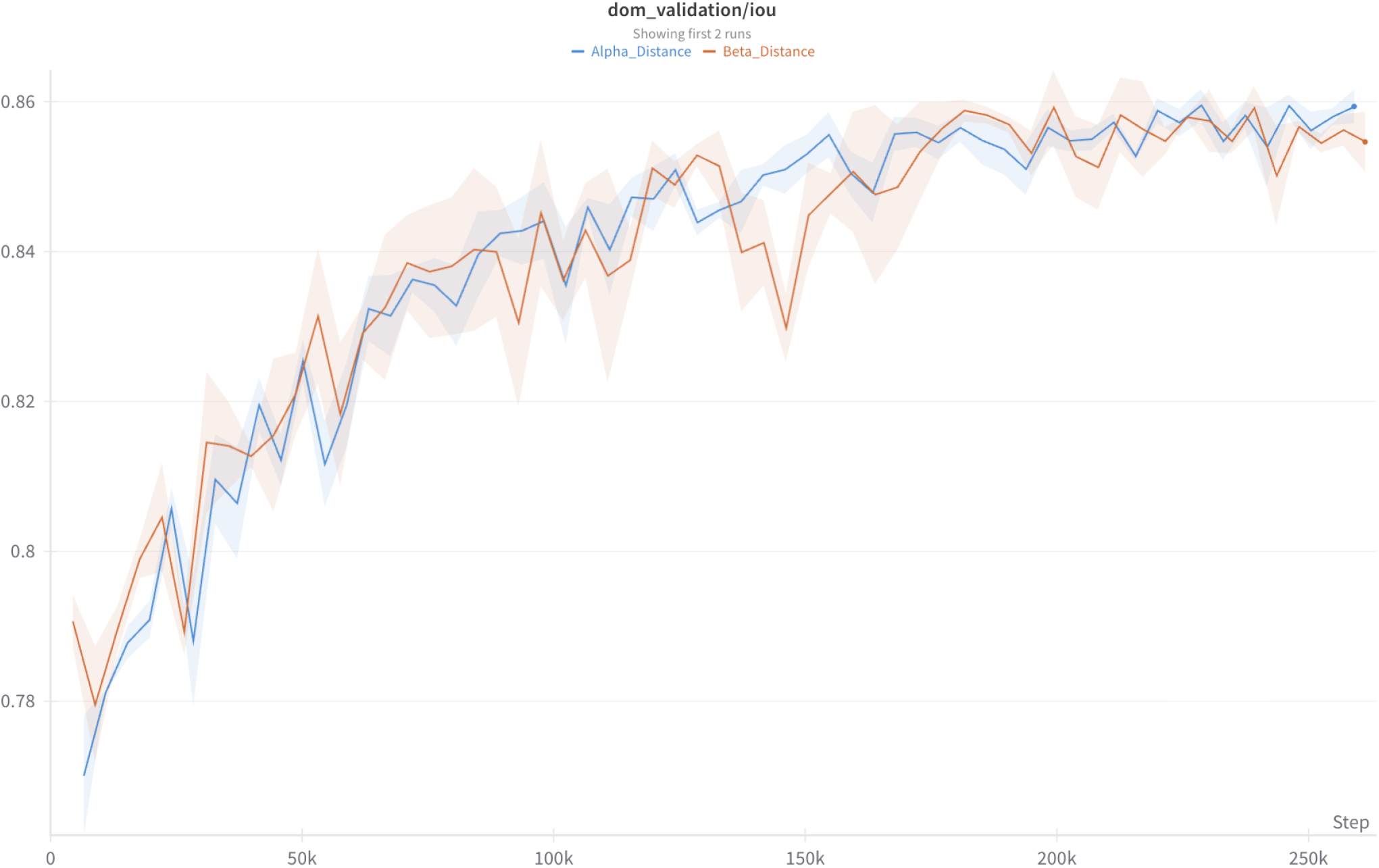
Results from training six Chainsaw models, three with *α*-carbon distances and three with *β*-carbon distances. The shaded range covers the minimum and maximum values at each epoch.

#### D.5 Model confidence and accuracy

**Figure 12.**
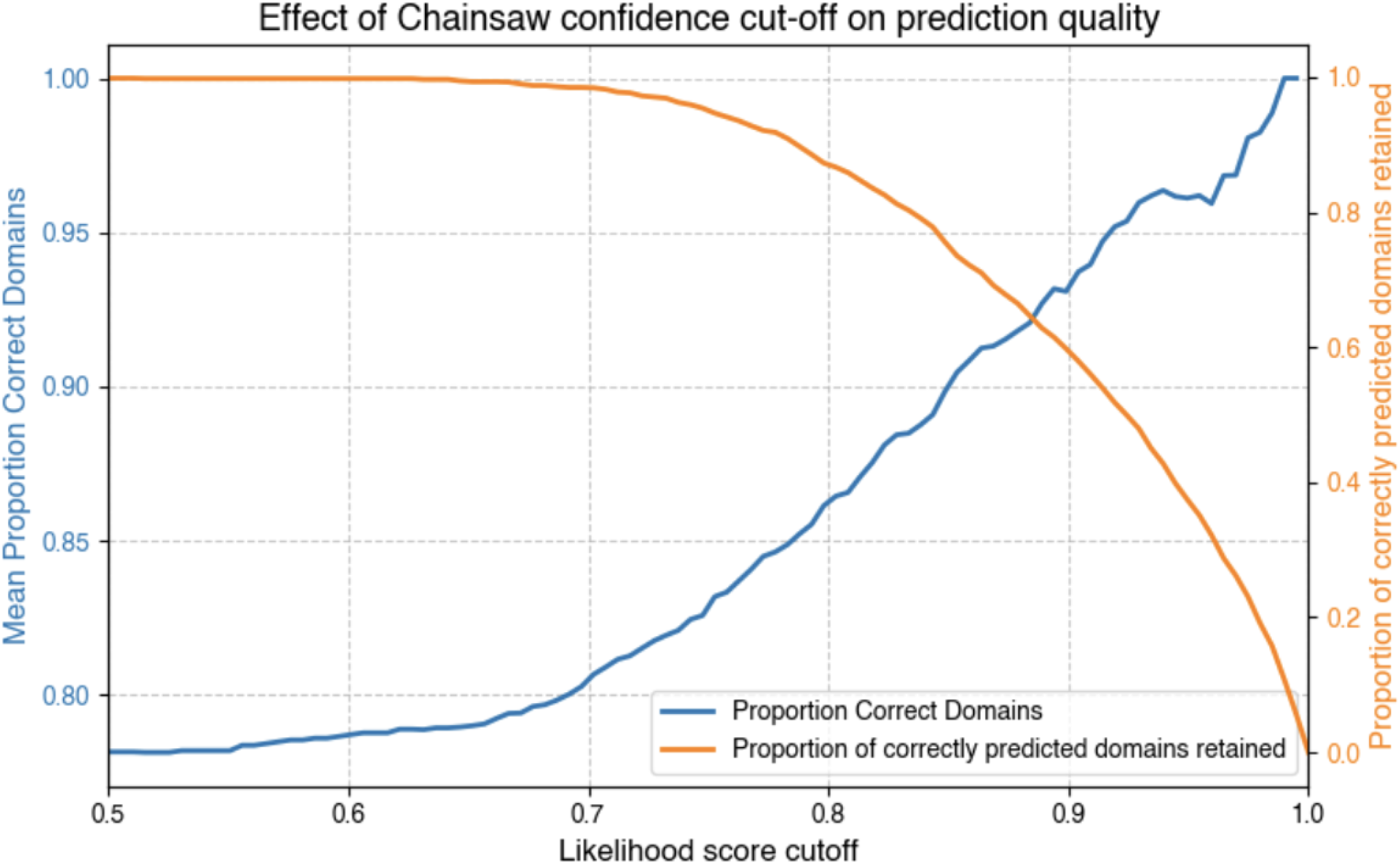
The Chainsaw confidence score reflects the likelihood of **A**^′^ (final assignment) under **Â** (output of neural network). We find that this is correlated with the accuracy of the predictions and can therefore be used as a filter to increase the precision of Chainsaw domain predictions albeit at the expense of recall.

**Figure 13.**
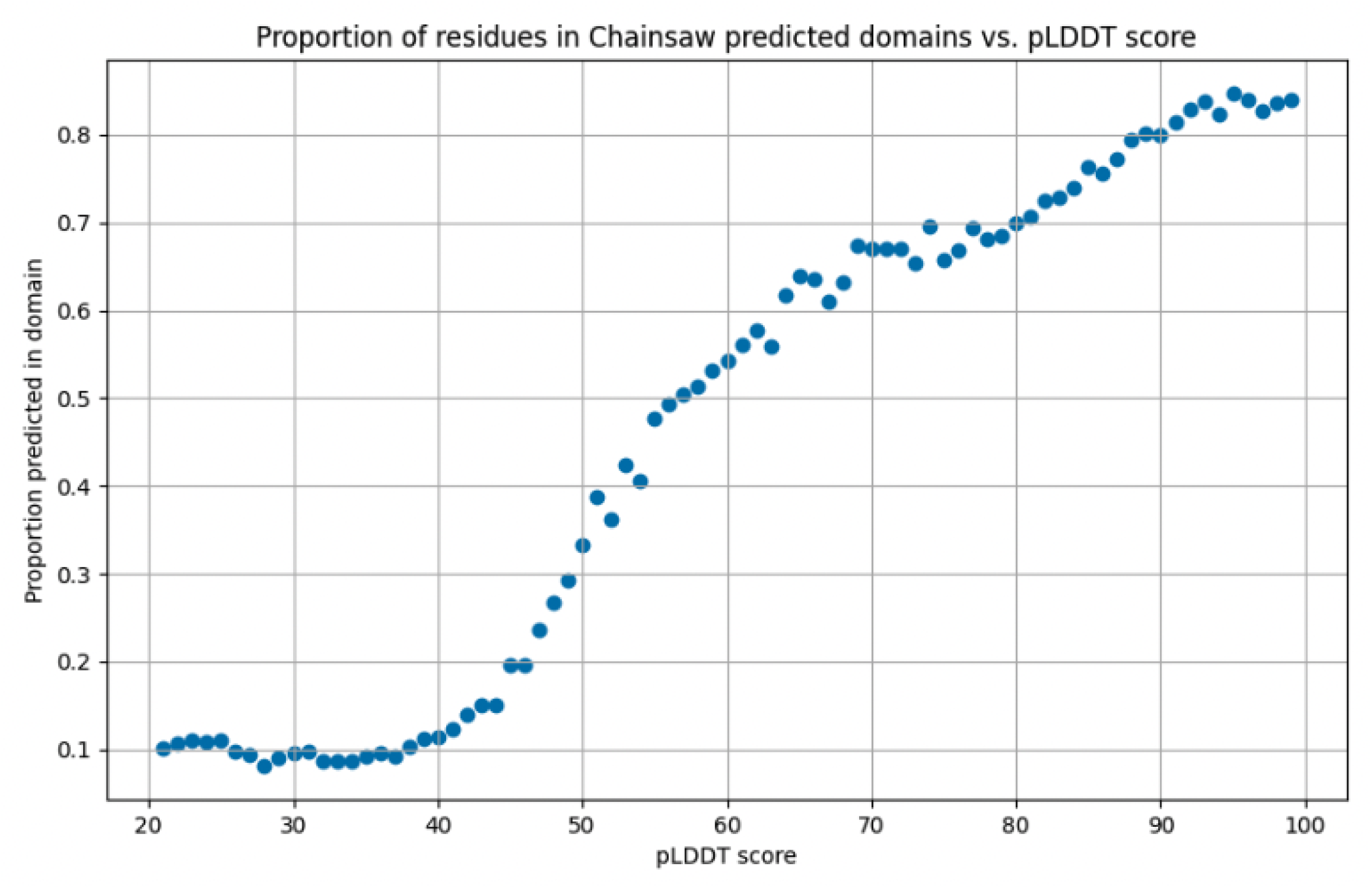
We calculate the proportion of residues that are predicted to be in a domain for each binned pLDDT score. Results were generated for 200 random AFDB models from the human proteome.

