## Supplementary material for "Chainsaw: protein domain segmentation with fully convolutional neural networks": Comparison of Chainsaw and UniDoc domain parsings on 200 AF models from human proteome: chainsaw_vs_unidoc_200_AF_human_blind_comparison_judgements.pdf

### 1 Chainsaw vs. UniDoc 200 human blind comparison

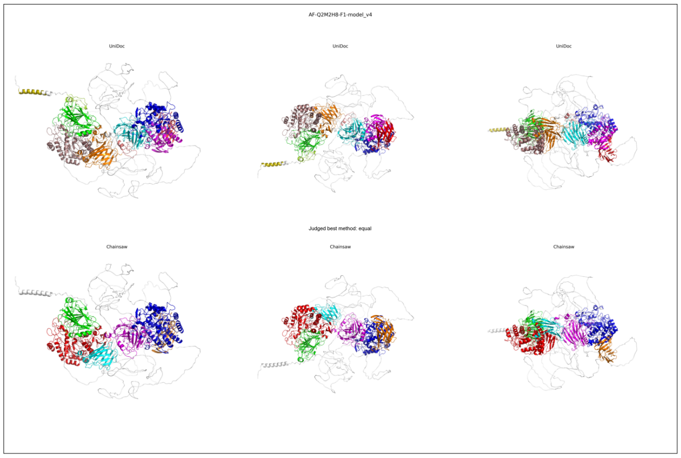


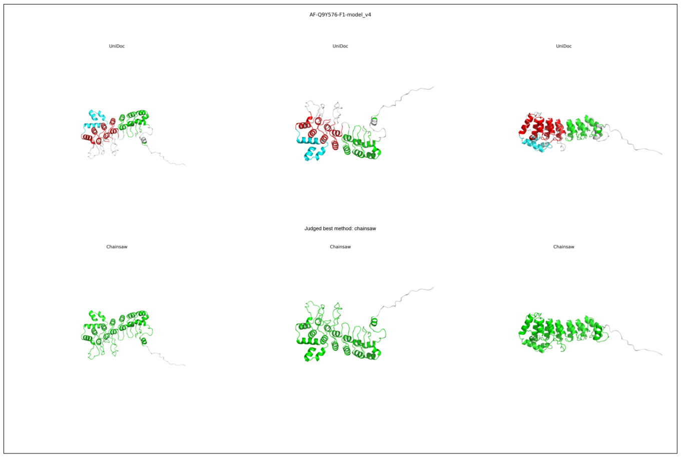


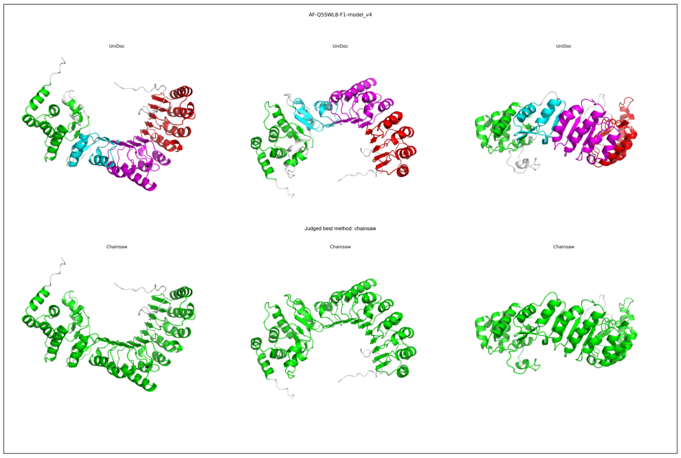


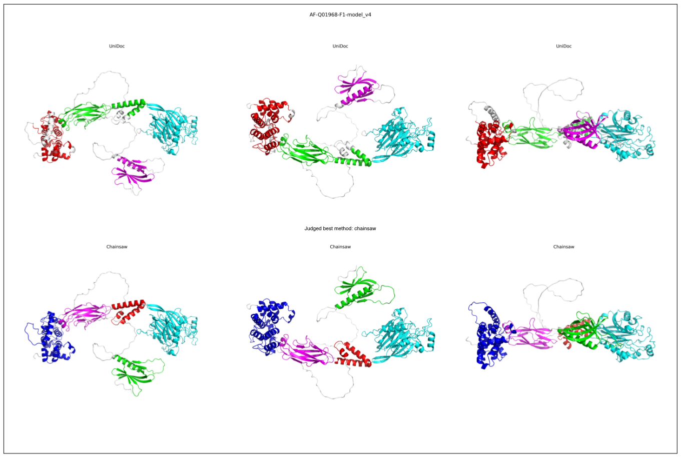


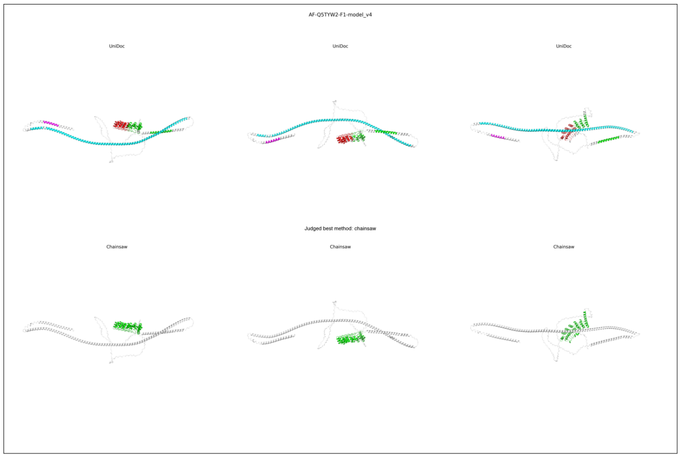


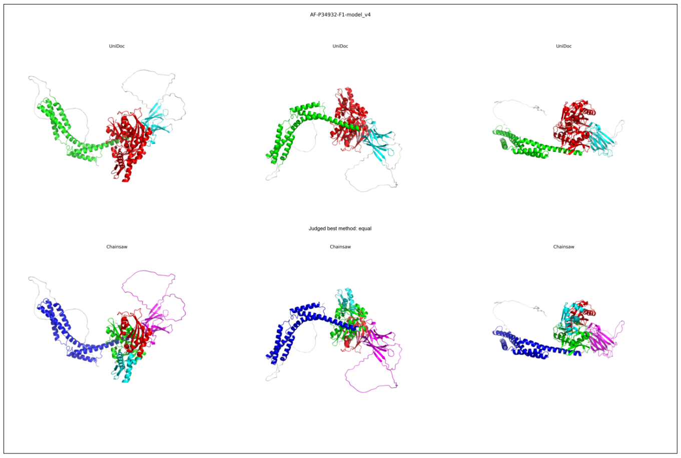


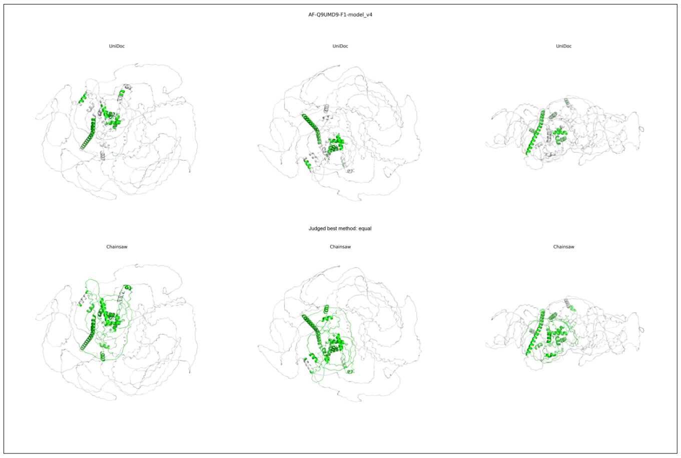


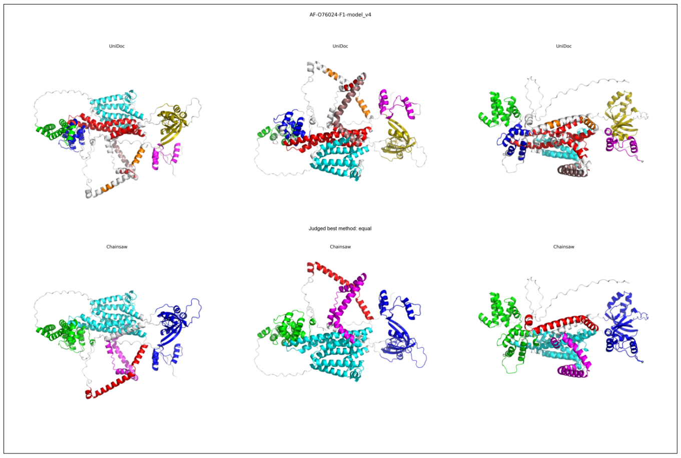

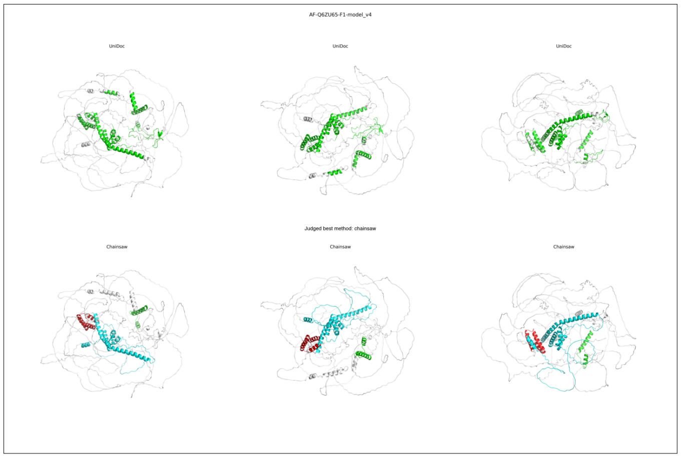
AF-Q6ZU65-F1-model_v4
UniDoc
UniDoc
UniDoc
Judged best method: chainsaw
Chainsaw
Chainsaw
Chainsaw

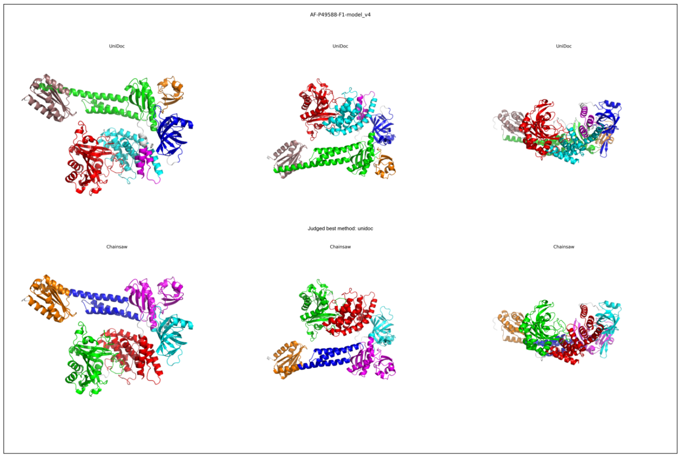
AF-P49588-F1-model_v4
UniDoc
UniDoc
UniDoc
Judged best method: unidoc
Chainsaw
Chainsaw
Chainsaw

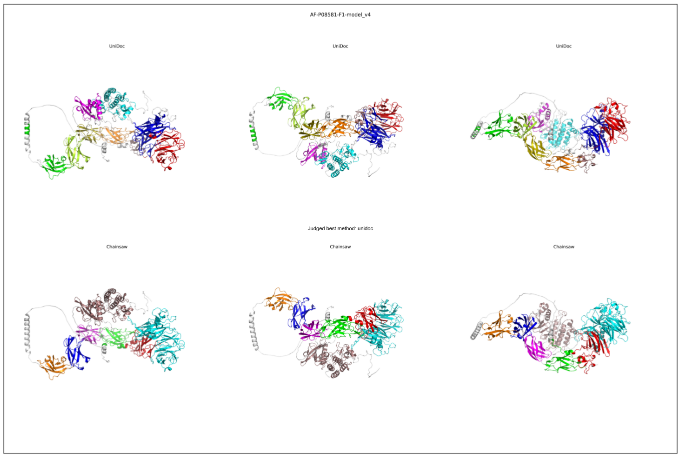
AF-P08581-F1-model_v4
UniDoc
UniDoc
UniDoc
Judged best method: unidoc
Chainsaw
Chainsaw
Chainsaw

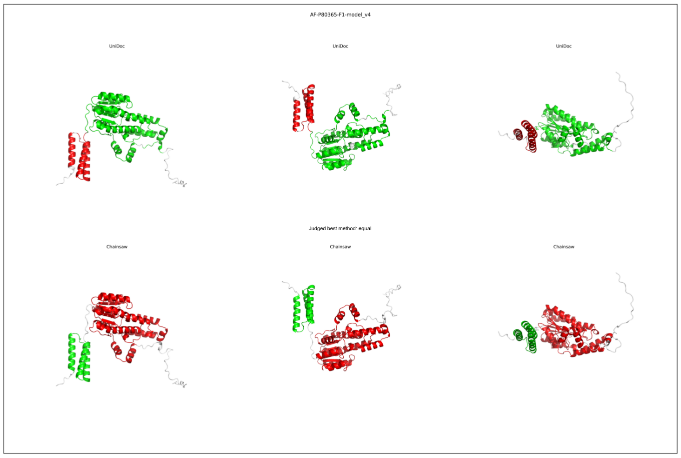
AF-P80365-F1-model_v4
UniDoc
UniDoc
UniDoc
Judged best method: equal
Chainsaw
Chainsaw
Chainsaw

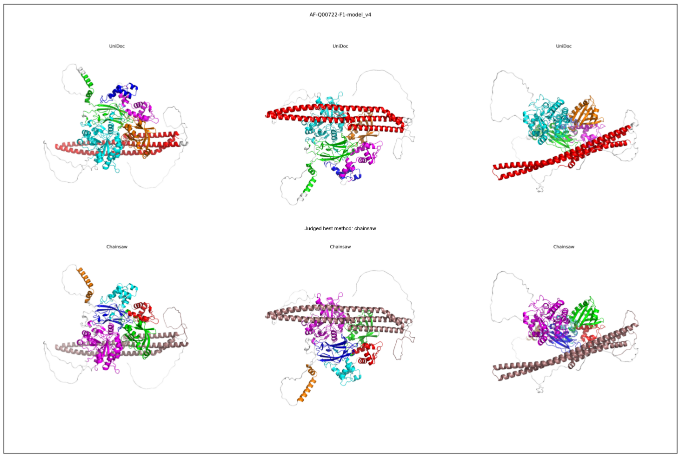
AF-Q00722-F1-model_v4
UniDoc
UniDoc
UniDoc
Judged best method: chainsaw
Chainsaw
Chainsaw
Chainsaw

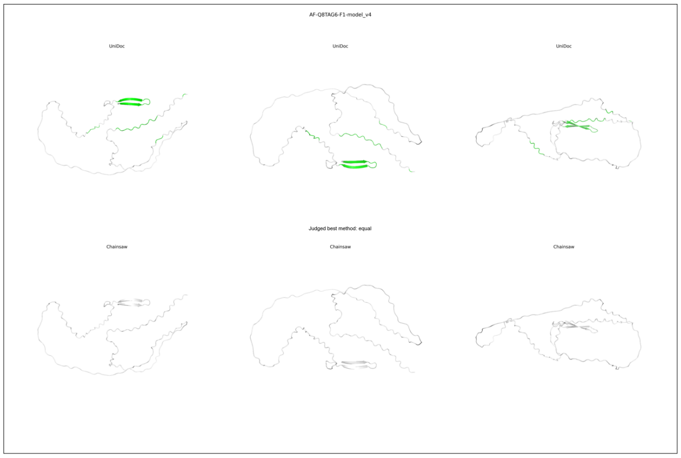
AF-Q8TAG6-F1-model_v4
UniDoc
UniDoc
UniDoc
Judged best method: equal
Chainsaw
Chainsaw
Chainsaw

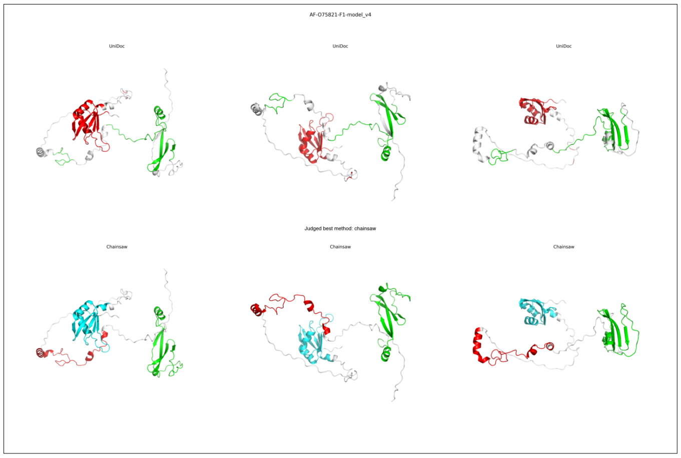
AF-O75821-F1-model_v4
UniDoc
UniDoc
UniDoc
Judged best method: chainsaw
Chainsaw
Chainsaw
Chainsaw

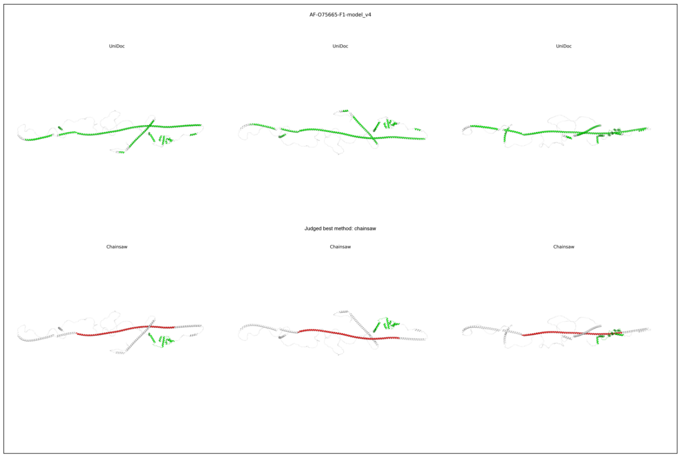
AF-O75665-F1-model_v4
UniDoc
UniDoc
UniDoc
Judged best method: chainsaw
Chainsaw
Chainsaw
Chainsaw

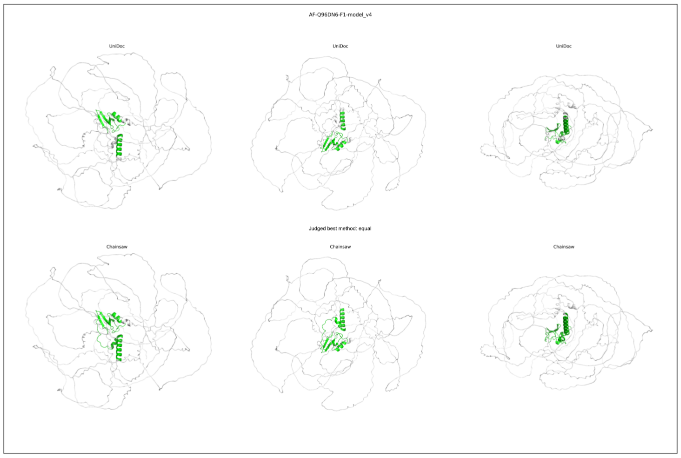
AF-Q96DN6-F1-model_v4
UniDoc
UniDoc
UniDoc
Judged best method: equal
Chainsaw
Chainsaw
Chainsaw

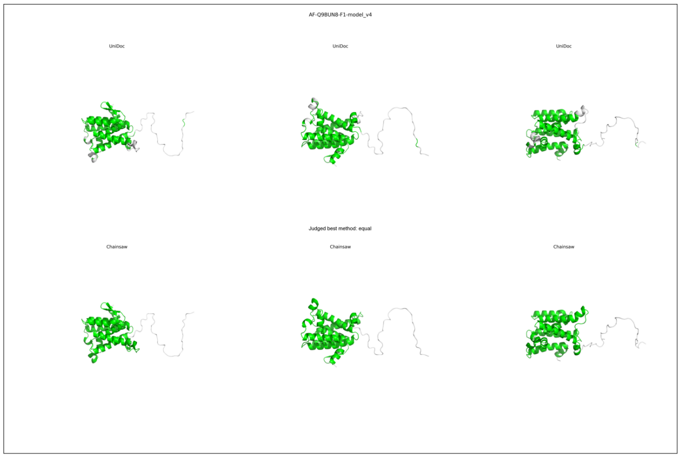
AF-Q9BUN8-F1-model_v4
UniDoc
UniDoc
UniDoc
Judged best method: equal
Chainsaw
Chainsaw
Chainsaw

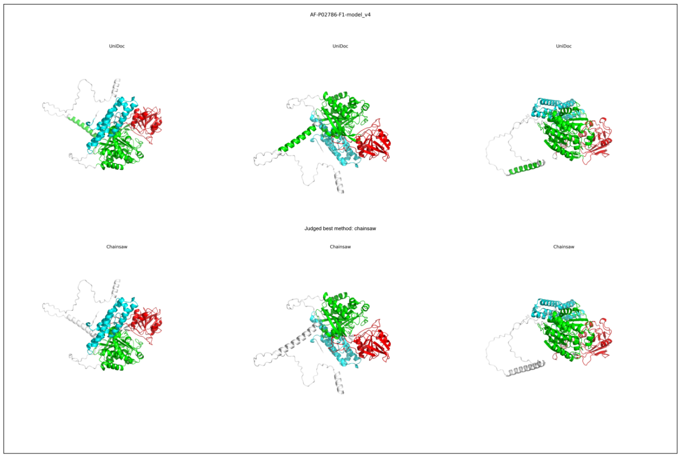
AF-P02786-F1-model_v4
UniDoc
UniDoc
UniDoc
Judged best method: chainsaw
Chainsaw
Chainsaw
Chainsaw

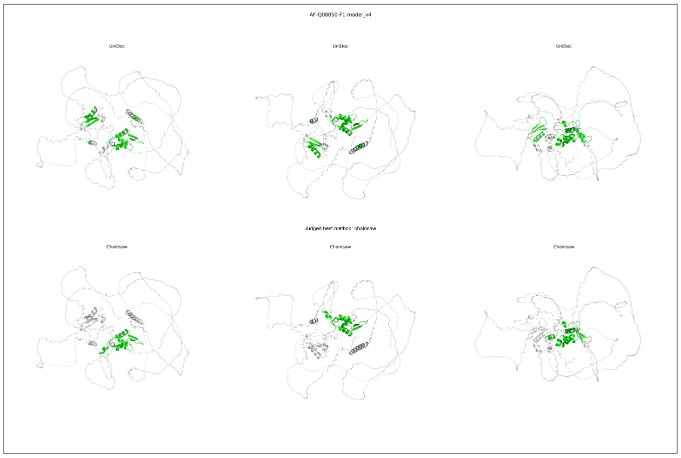
AF-Q08050-F1-model_v4
UniDoc
UniDoc
UniDoc
Judged best method: chainsaw
Chainsaw
Chainsaw
Chainsaw

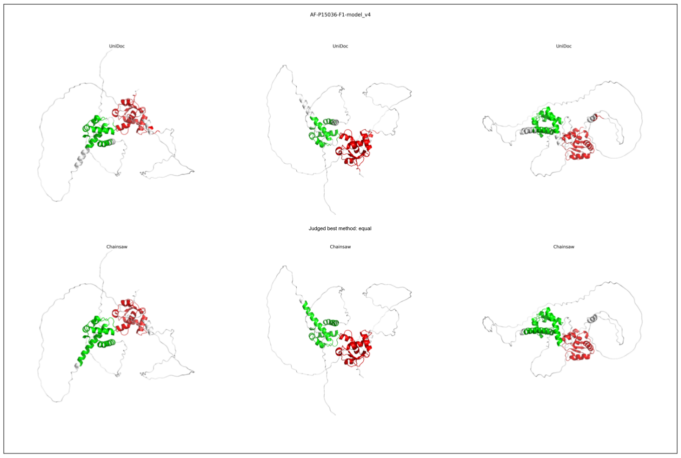
AF-P15036-F1-model_v4
UniDoc
UniDoc
UniDoc
Judged best method: equal
Chainsaw
Chainsaw
Chainsaw

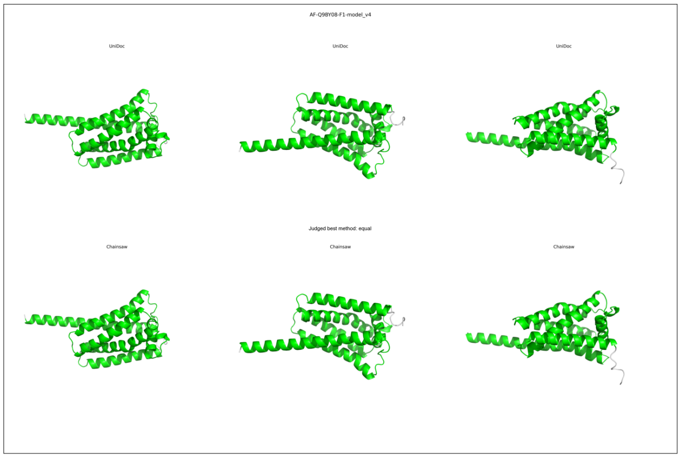
AF-Q9BY08-F1-model_v4
UniDoc
UniDoc
UniDoc
Judged best method: equal
Chainsaw
Chainsaw
Chainsaw

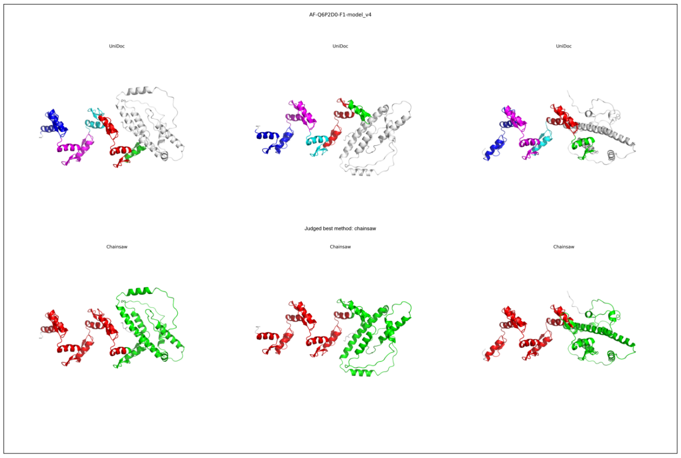
AF-Q6P2D0-F1-model_v4
UniDoc
UniDoc
UniDoc
Judged best method: chainsaw
Chainsaw
Chainsaw
Chainsaw

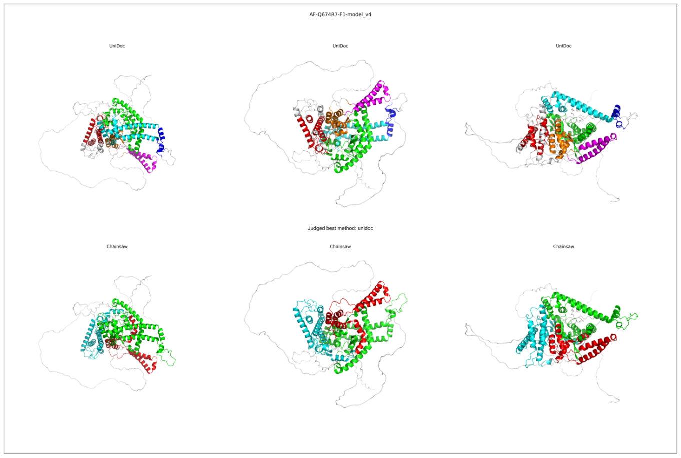
AF-Q674R7-F1-model_v4
UniDoc
UniDoc
UniDoc
Judged best method: unidoc
Chainsaw
Chainsaw
Chainsaw

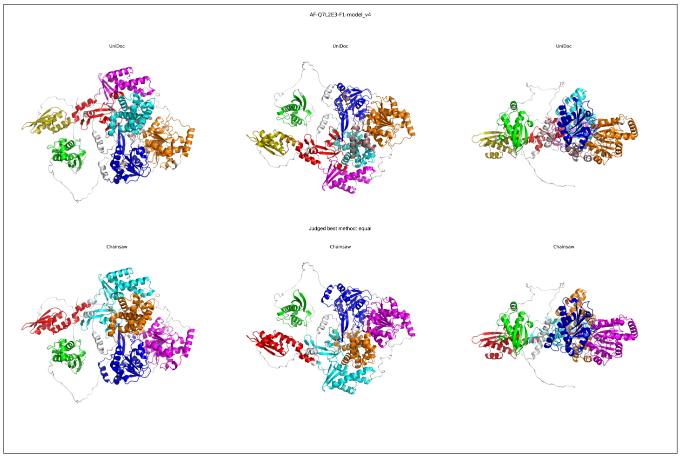
AF-Q7L2E3-F1-model_v4
UniDoc
UniDoc
UniDoc
Judged best method: equal
Chainsaw
Chainsaw
Chainsaw

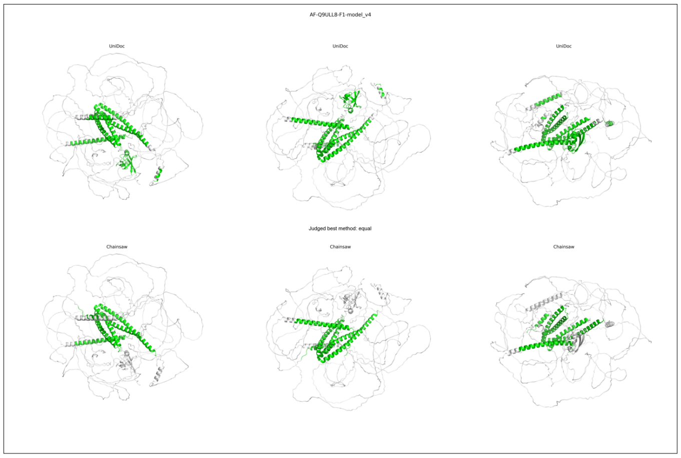
AF-Q9ULL8-F1-model_v4
UniDoc
UniDoc
UniDoc
Judged best method: equal
Chainsaw
Chainsaw
Chainsaw

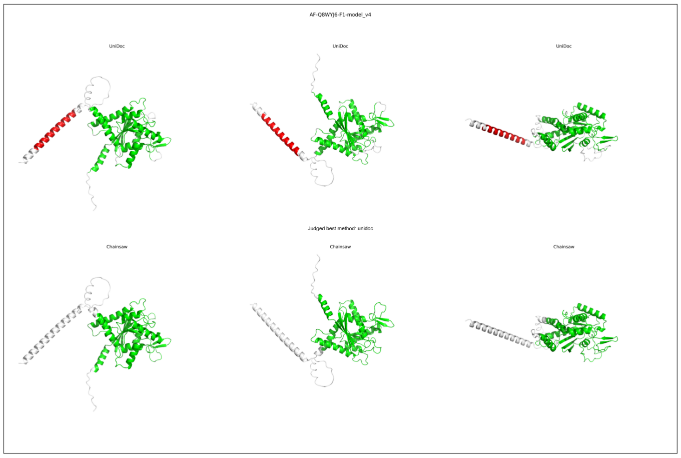
AF-Q8WYJ6-F1-model_v4
UniDoc
UniDoc
UniDoc
Judged best method: unidoc
Chainsaw
Chainsaw
Chainsaw

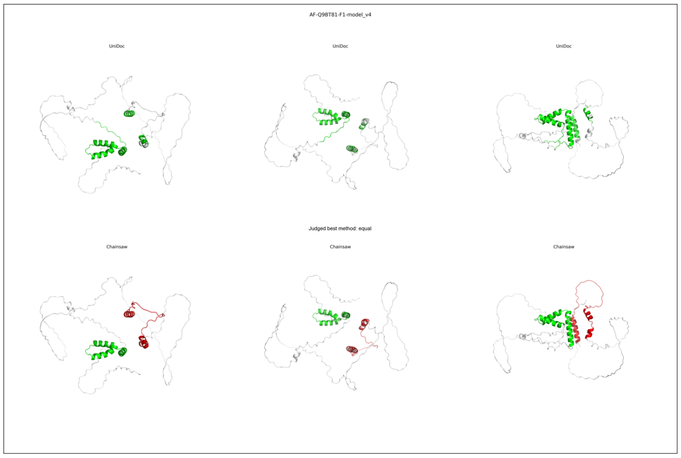
AF-Q9BT81-F1-model_v4
UniDoc
UniDoc
UniDoc
Judged best method: equal
Chainsaw
Chainsaw
Chainsaw

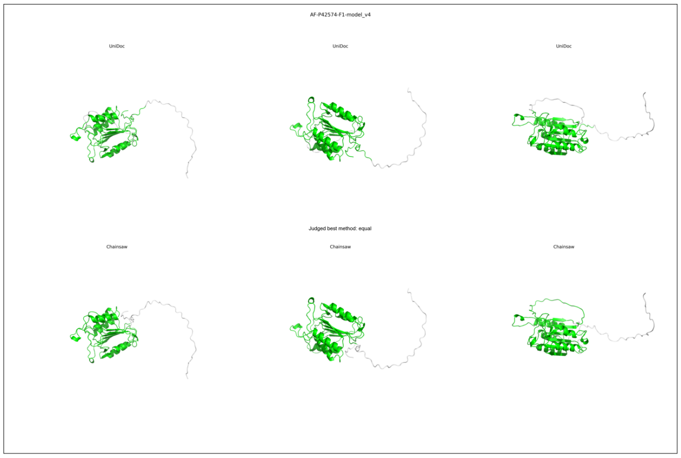
AF-P42574-F1-model_v4
UniDoc
UniDoc
UniDoc
Judged best method: equal
Chainsaw
Chainsaw
Chainsaw

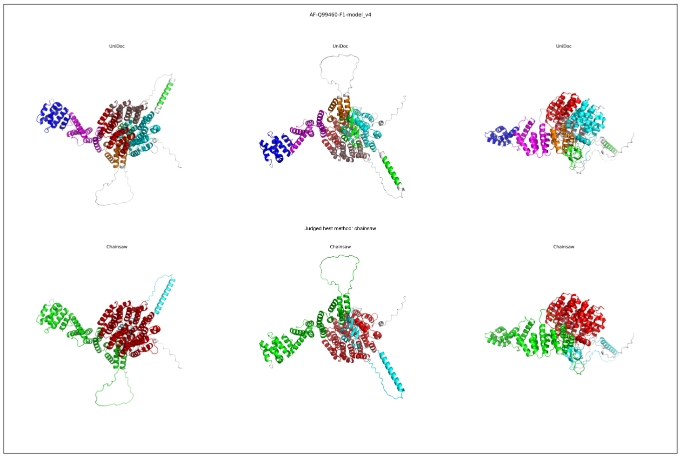
AF-Q99460-F1-model_v4
UniDoc
UniDoc
UniDoc
Judged best method: chainsaw
Chainsaw
Chainsaw
Chainsaw

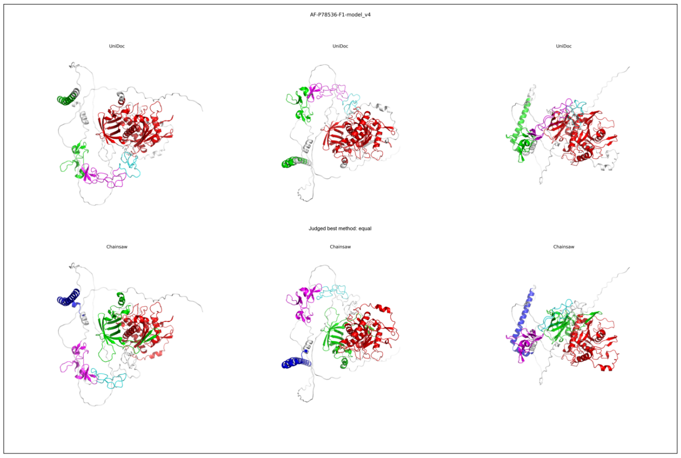
AF-P78536-F1-model_v4
UniDoc
UniDoc
UniDoc
Judged best method: equal
Chainsaw
Chainsaw
Chainsaw

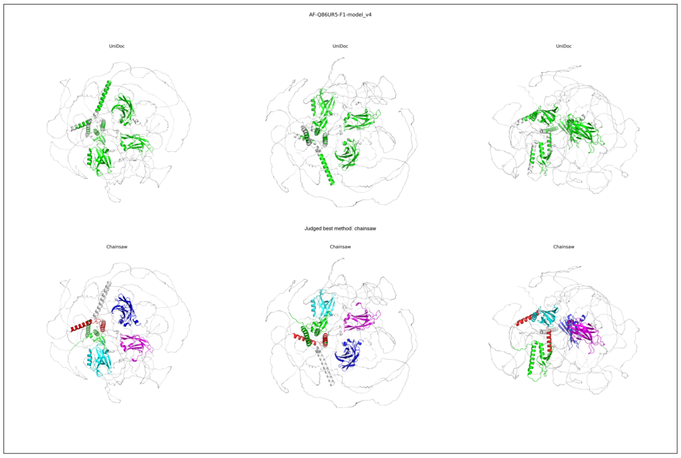
AF-Q86UR5-F1-model_v4
UniDoc
UniDoc
UniDoc
Judged best method: chainsaw
Chainsaw
Chainsaw
Chainsaw

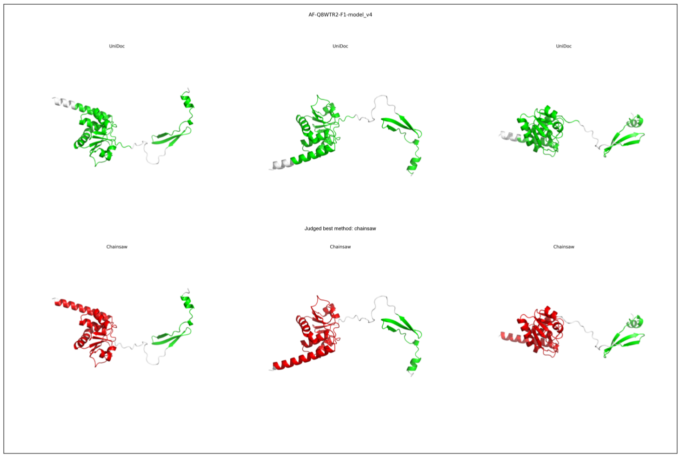

AF-Q8WTR2-F1-model_v4
UniDoc
UniDoc
UniDoc
Judged best method: chainsaw
Chainsaw
Chainsaw
Chainsaw


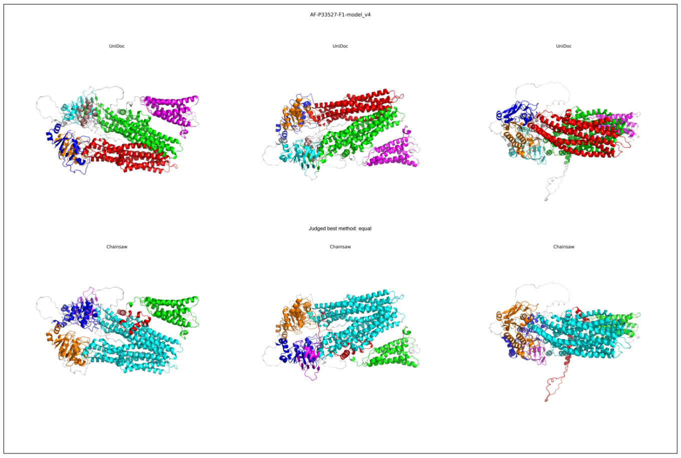

AF-P33527-F1-model_v4
UniDoc
UniDoc
UniDoc
Judged best method: equal
Chainsaw
Chainsaw
Chainsaw


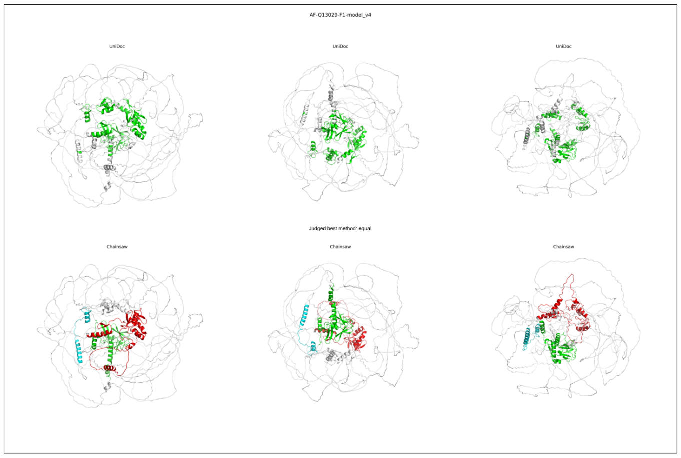

AF-Q13029-F1-model_v4
UniDoc
UniDoc
UniDoc
Judged best method: equal
Chainsaw
Chainsaw
Chainsaw


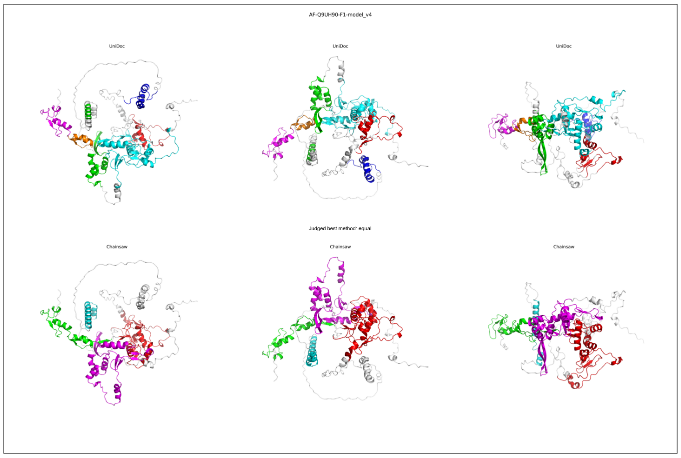

AF-Q9UH90-F1-model_v4
UniDoc
UniDoc
UniDoc
Judged best method: equal
Chainsaw
Chainsaw
Chainsaw


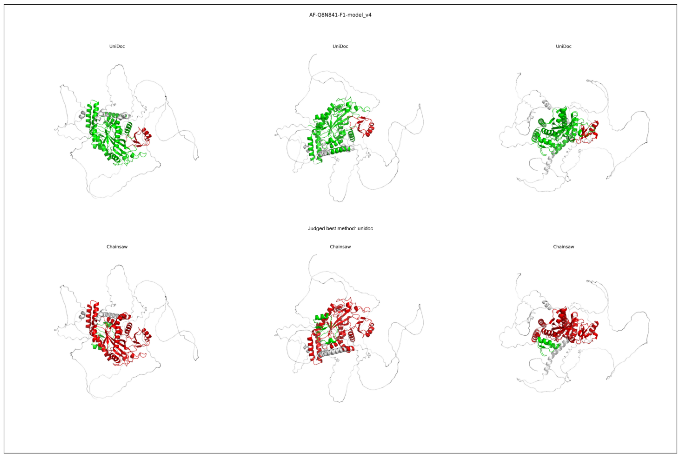

AF-Q8N841-F1-model_v4
UniDoc
UniDoc
UniDoc
Judged best method: unidoc
Chainsaw
Chainsaw
Chainsaw


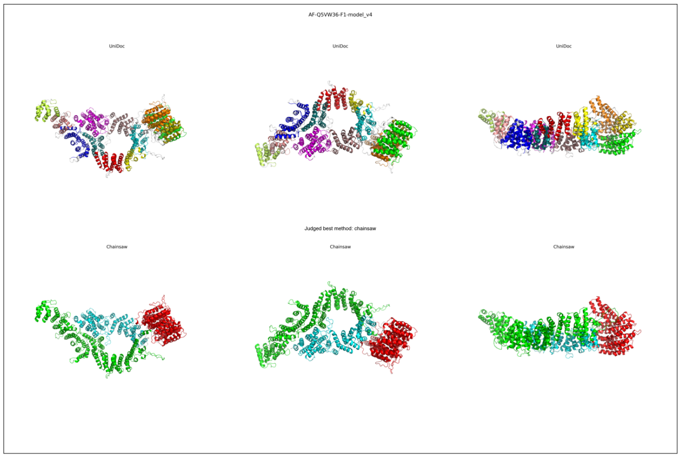

AF-Q5VW36-F1-model_v4
UniDoc
UniDoc
UniDoc
Judged best method: chainsaw
Chainsaw
Chainsaw
Chainsaw


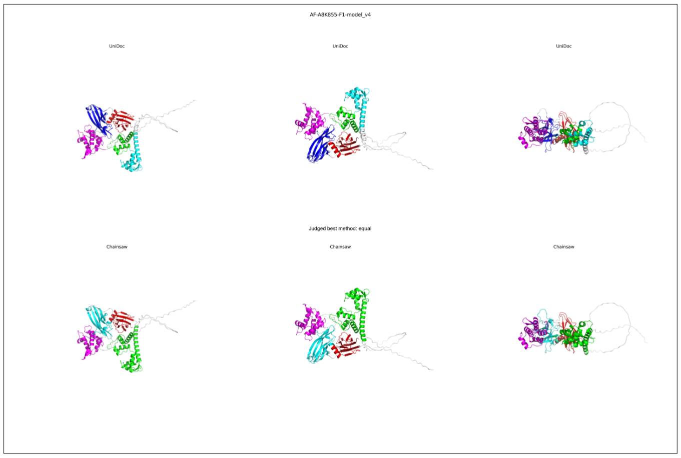

AF-A8K855-F1-model_v4
UniDoc
UniDoc
UniDoc
Judged best method: equal
Chainsaw
Chainsaw
Chainsaw


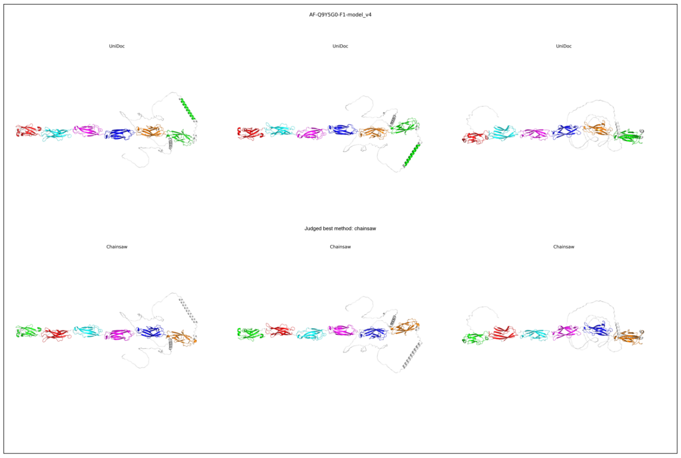

AF-Q9Y5G0-F1-model_v4
UniDoc
UniDoc
UniDoc
Judged best method: chainsaw
Chainsaw
Chainsaw
Chainsaw

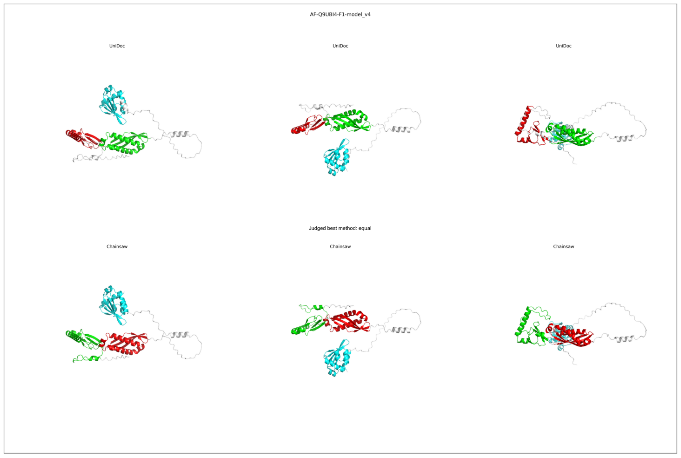
AF-Q9UBI4-F1-model_v4
UniDoc
UniDoc
UniDoc
Judged best method: equal
Chainsaw
Chainsaw
Chainsaw

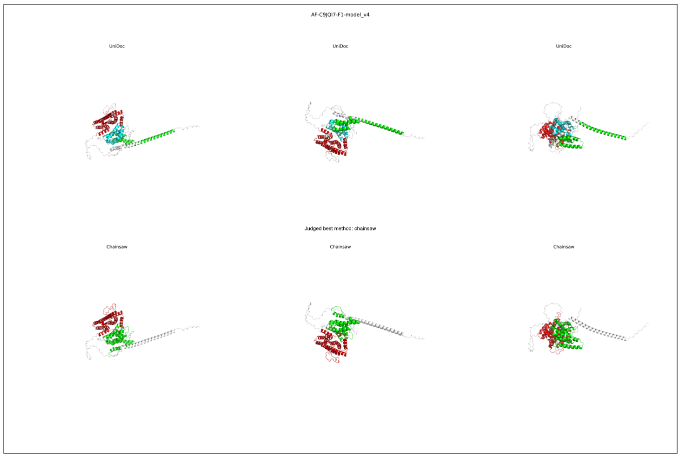
AF-C9JQI7-F1-model_v4
UniDoc
UniDoc
UniDoc
Judged best method: chainsaw
Chainsaw
Chainsaw
Chainsaw

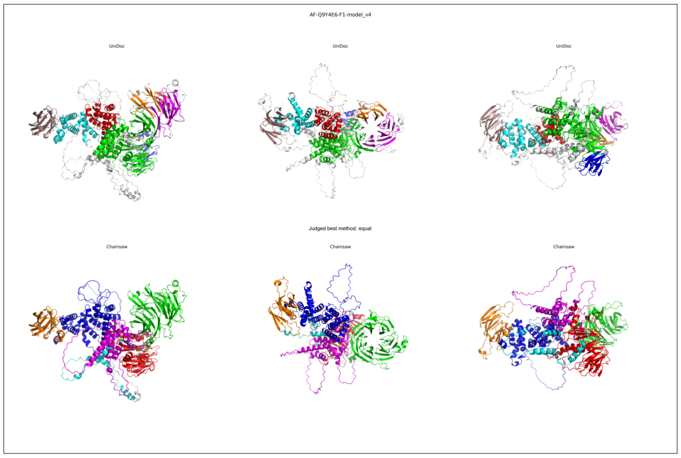
AF-Q9Y4E6-F1-model_v4
UniDoc
UniDoc
UniDoc
Judged best method: equal
Chainsaw
Chainsaw
Chainsaw

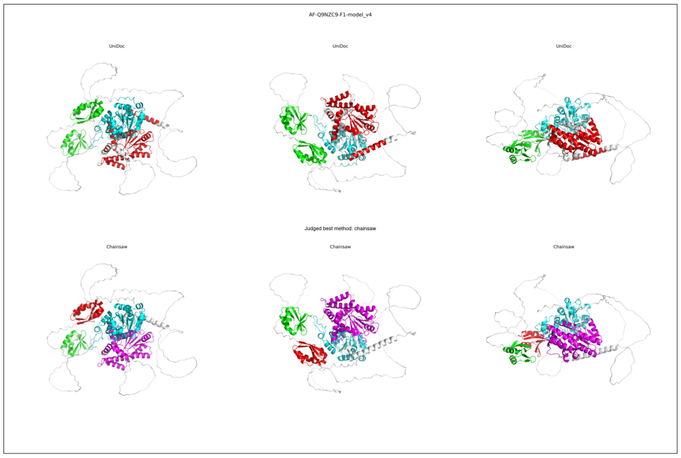
AF-Q9NZC9-F1-model_v4
UniDoc
UniDoc
UniDoc
Judged best method: chainsaw
Chainsaw
Chainsaw
Chainsaw

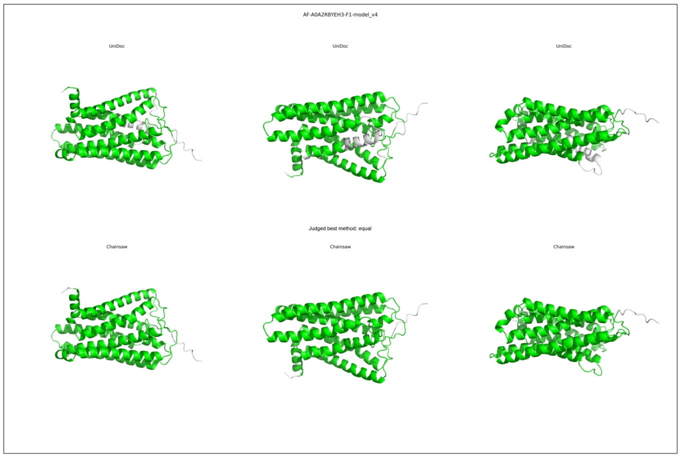
AF-A0A2R8YEH3-F1-model_v4
UniDoc
UniDoc
UniDoc
Judged best method: equal
Chainsaw
Chainsaw
Chainsaw

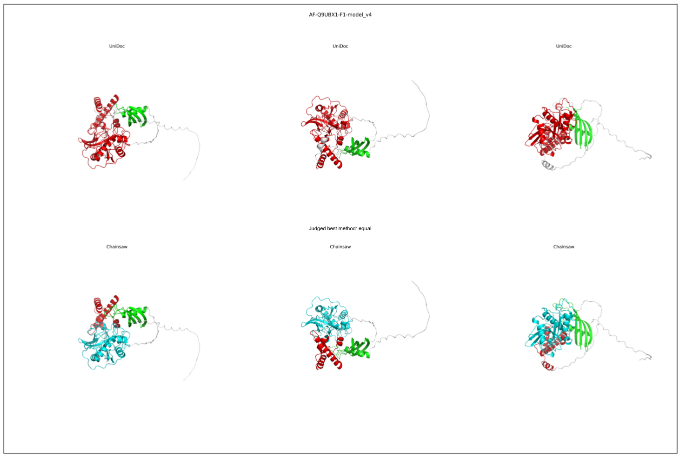
AF-Q9UBX1-F1-model_v4
UniDoc
UniDoc
UniDoc
Judged best method: equal
Chainsaw
Chainsaw
Chainsaw

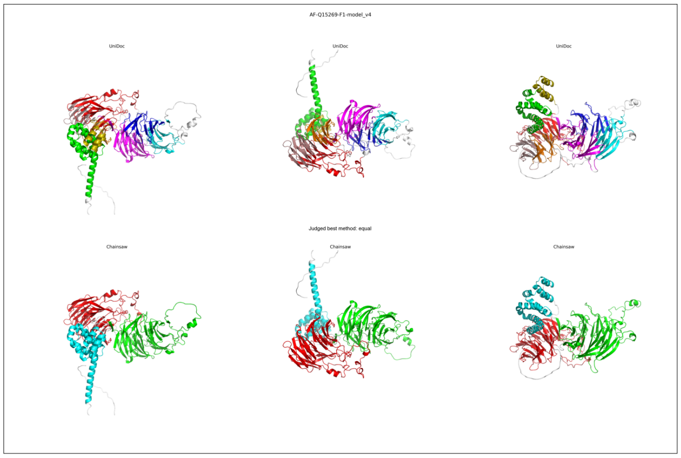
AF-Q15269-F1-model_v4
UniDoc
UniDoc
UniDoc
Judged best method: equal
Chainsaw
Chainsaw
Chainsaw

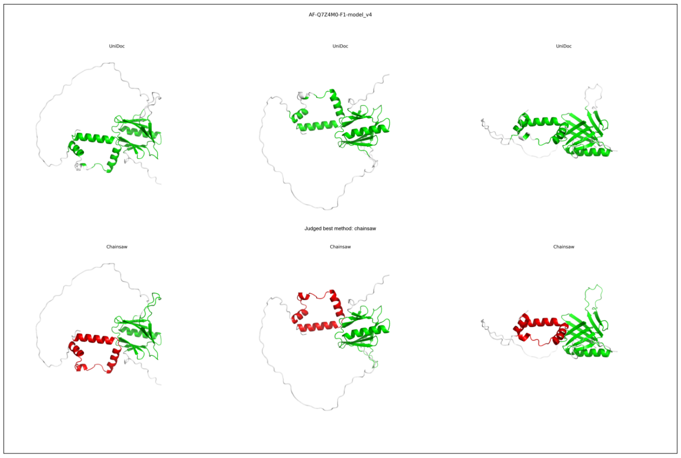
AF-Q7Z4M0-F1-model_v4
UniDoc
UniDoc
UniDoc
Judged best method: chainsaw
Chainsaw
Chainsaw
Chainsaw

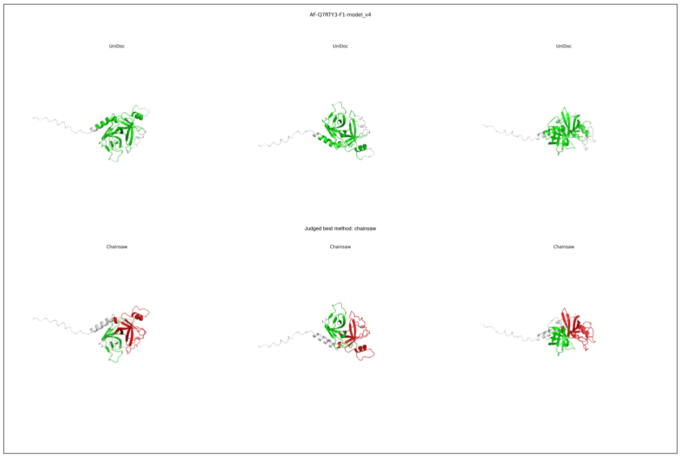
AF-Q7RTY3-F1-model_v4
UniDoc
UniDoc
UniDoc
Judged best method: chainsaw
Chainsaw
Chainsaw
Chainsaw

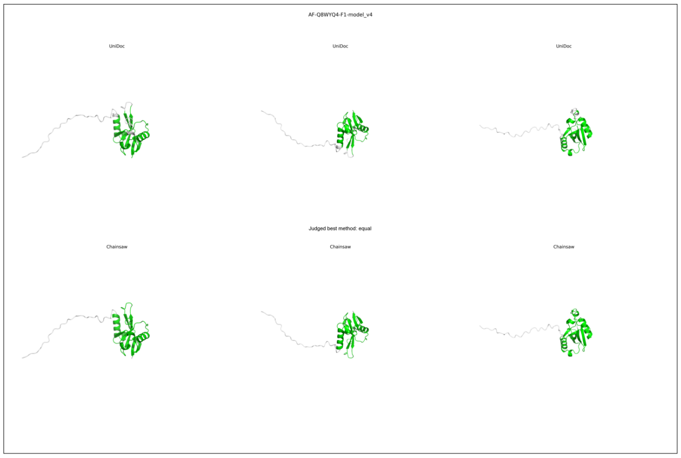
AF-Q8WYQ4-F1-model_v4
UniDoc
UniDoc
UniDoc
Judged best method: equal
Chainsaw
Chainsaw
Chainsaw

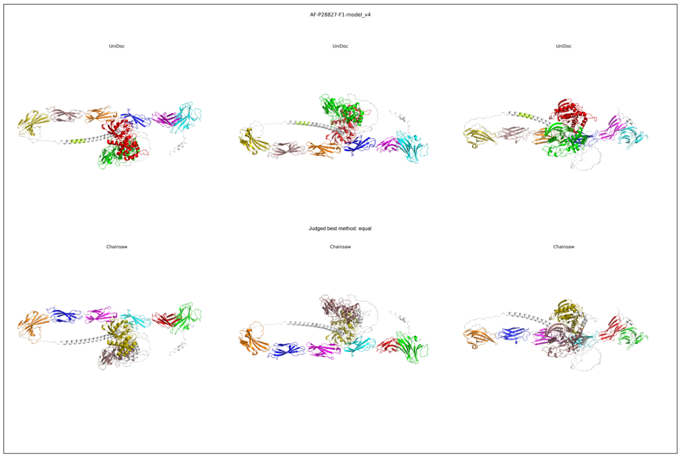
AF-P28827-F1-model_v4
UniDoc
UniDoc
UniDoc
Judged best method: equal
Chainsaw
Chainsaw
Chainsaw

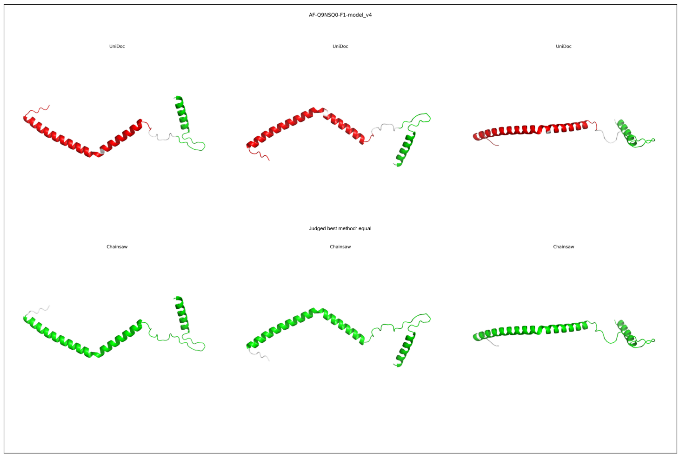
AF-Q9NSQ0-F1-model_v4
UniDoc
UniDoc
UniDoc
Judged best method: equal
Chainsaw
Chainsaw
Chainsaw

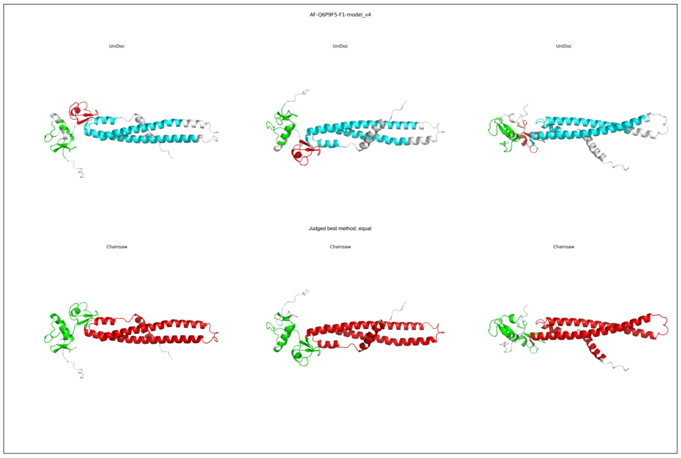
AF-Q6P9F5-F1-model_v4
UniDoc
UniDoc
UniDoc
Judged best method: equal
Chainsaw
Chainsaw
Chainsaw

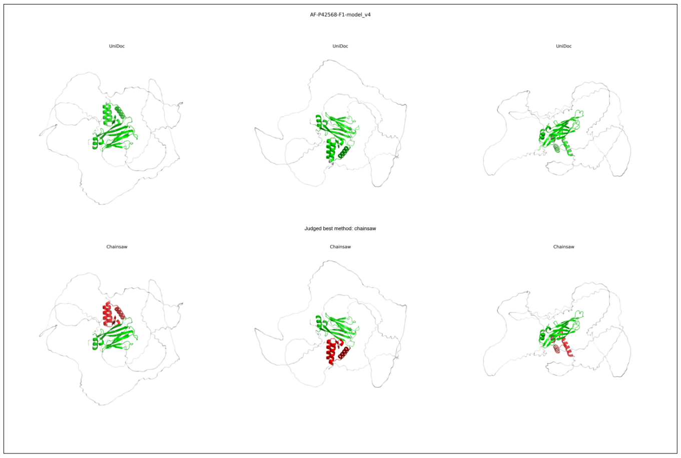
AF-P42568-F1-model_v4
UniDoc
UniDoc
UniDoc
Judged best method: chainsaw
Chainsaw
Chainsaw
Chainsaw

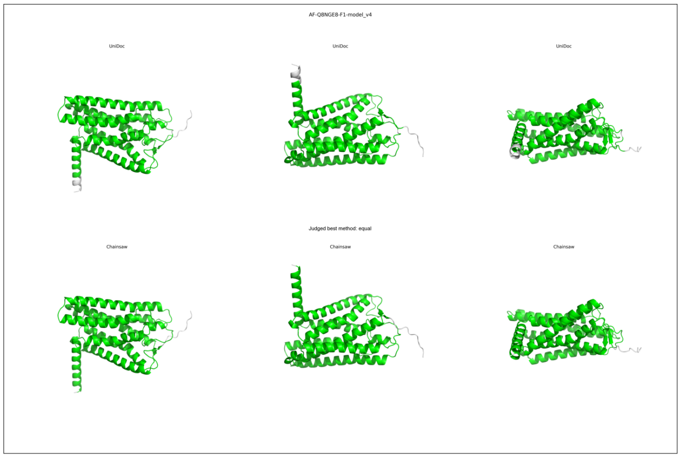
AF-Q8NGE8-F1-model_v4
UniDoc
UniDoc
UniDoc
Judged best method: equal
Chainsaw
Chainsaw
Chainsaw

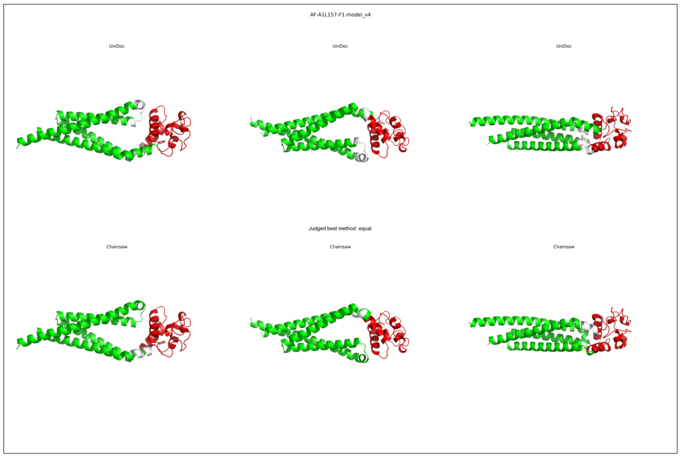
AF-A1L157-F1-model_v4
UniDoc
UniDoc
UniDoc
Judged best method: equal
Chainsaw
Chainsaw
Chainsaw

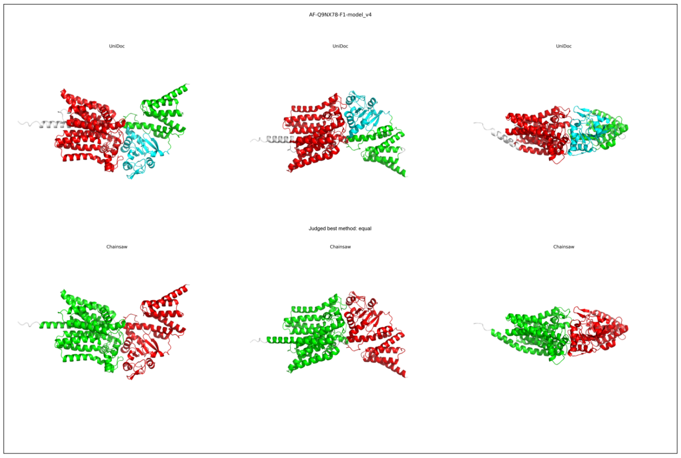
AF-Q9NX78-F1-model_v4
UniDoc
UniDoc
UniDoc
Judged best method: equal
Chainsaw
Chainsaw
Chainsaw

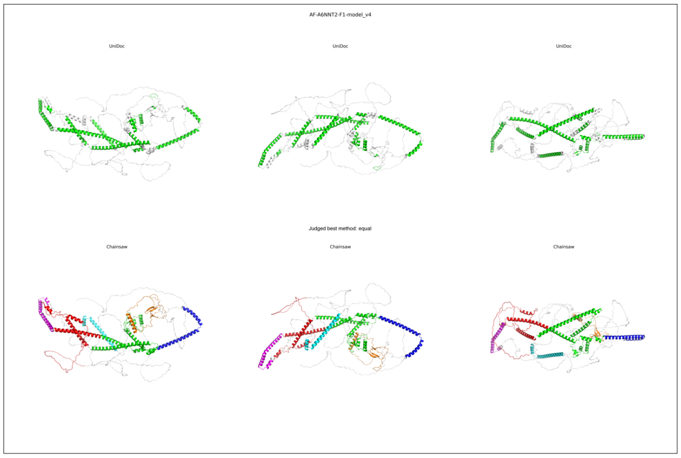
AF-A6NNT2-F1-model_v4
UniDoc
UniDoc
UniDoc
Judged best method: equal
Chainsaw
Chainsaw
Chainsaw

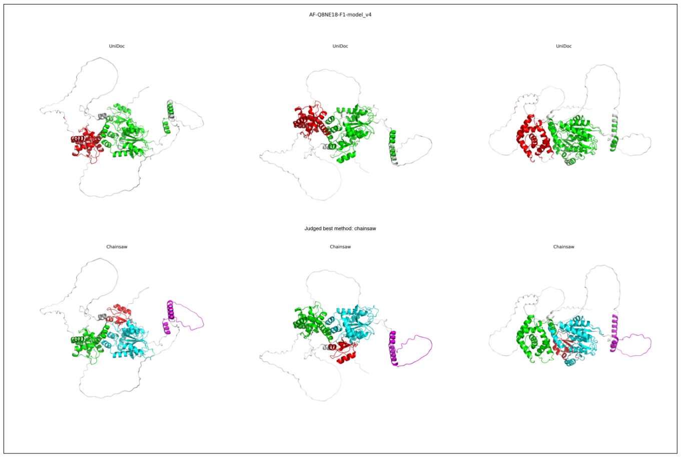
AF-Q8NE18-F1-model_v4
UniDoc
UniDoc
UniDoc
Judged best method: chainsaw
Chainsaw
Chainsaw
Chainsaw

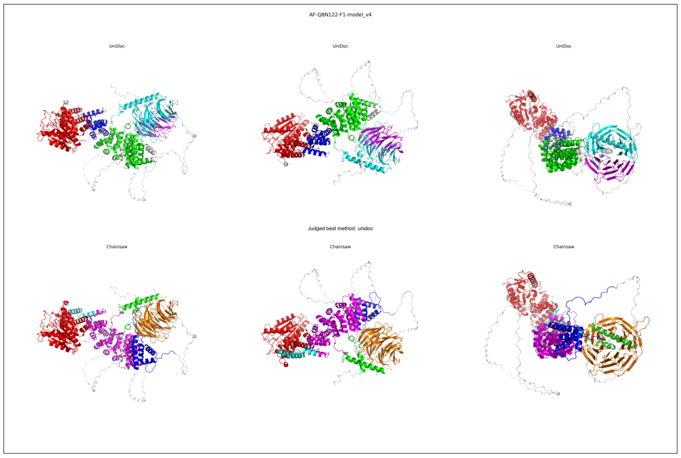
AF-Q8N122-F1-model_v4
UniDoc
UniDoc
UniDoc
Judged best method: unidoc
Chainsaw
Chainsaw
Chainsaw

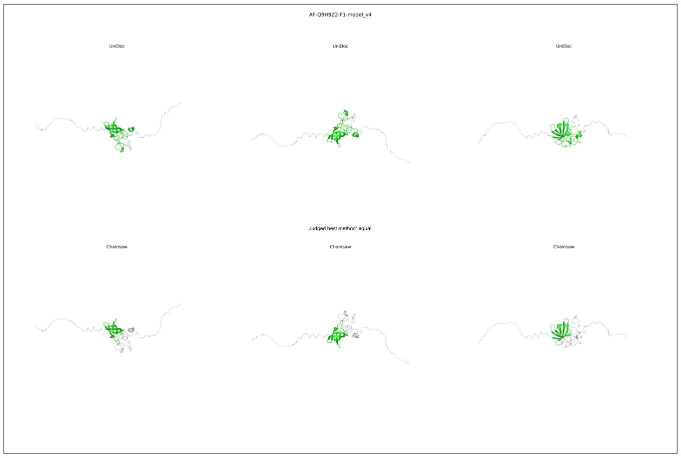
AF-Q9H9Z2-F1-model_v4
UniDoc
UniDoc
UniDoc
Judged best method: equal
Chainsaw
Chainsaw
Chainsaw

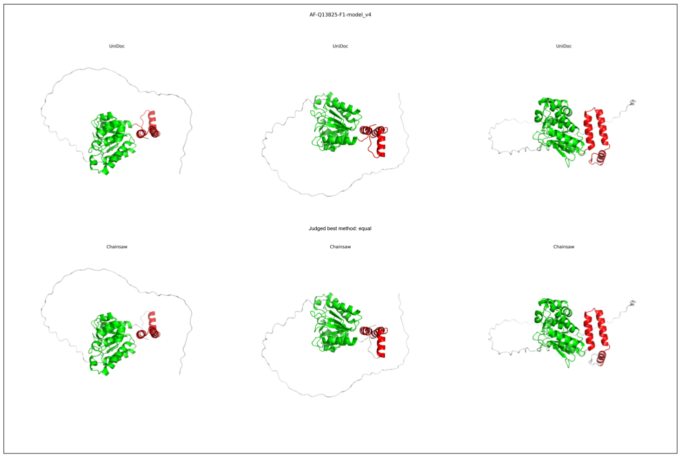
AF-Q13825-F1-model_v4
UniDoc
UniDoc
UniDoc
Judged best method: equal
Chainsaw
Chainsaw
Chainsaw

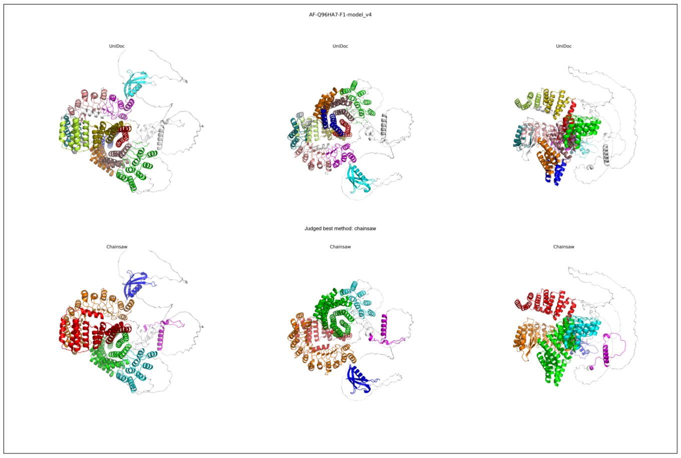
AF-Q96HA7-F1-model_v4
UniDoc
UniDoc
UniDoc
Judged best method: chainsaw
Chainsaw
Chainsaw
Chainsaw

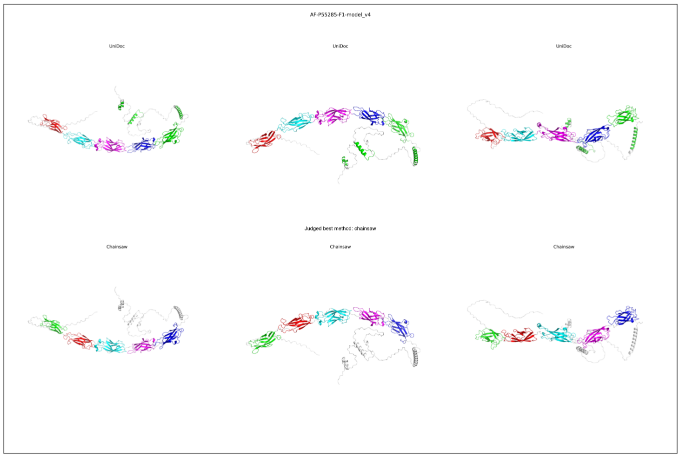
AF-P55285-F1-model_v4
UniDoc
UniDoc
UniDoc
Judged best method: chainsaw
Chainsaw
Chainsaw
Chainsaw

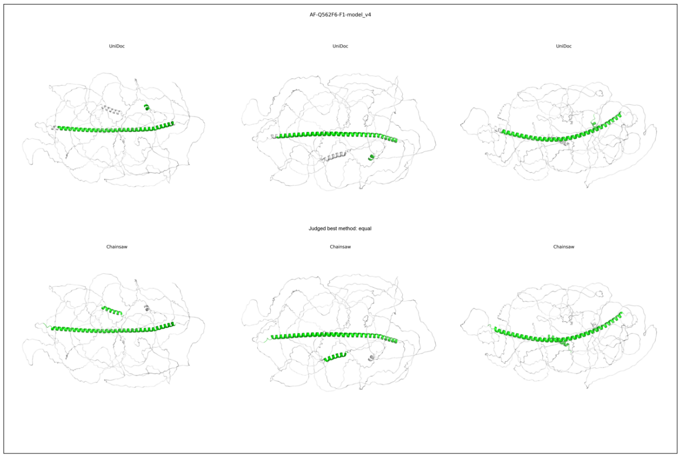
AF-Q562F6-F1-model_v4
UniDoc
UniDoc
UniDoc
Judged best method: equal
Chainsaw
Chainsaw
Chainsaw

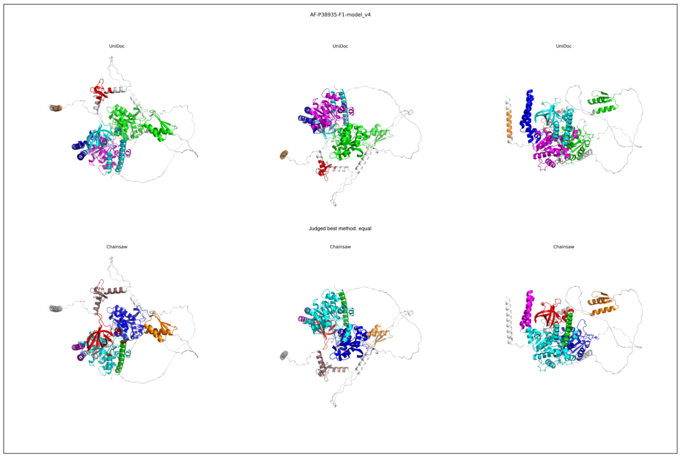
AF-P38935-F1-model_v4
UniDoc
UniDoc
UniDoc
Judged best method: equal
Chainsaw
Chainsaw
Chainsaw

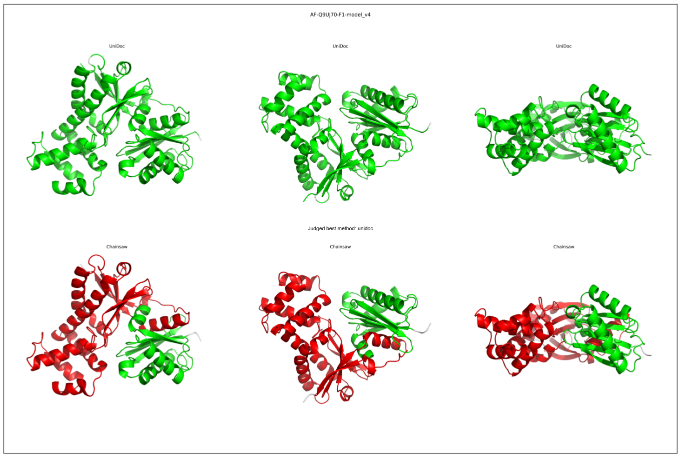
AF-Q9UJ70-F1-model_v4
UniDoc
UniDoc
UniDoc
Judged best method: unidoc
Chainsaw
Chainsaw
Chainsaw

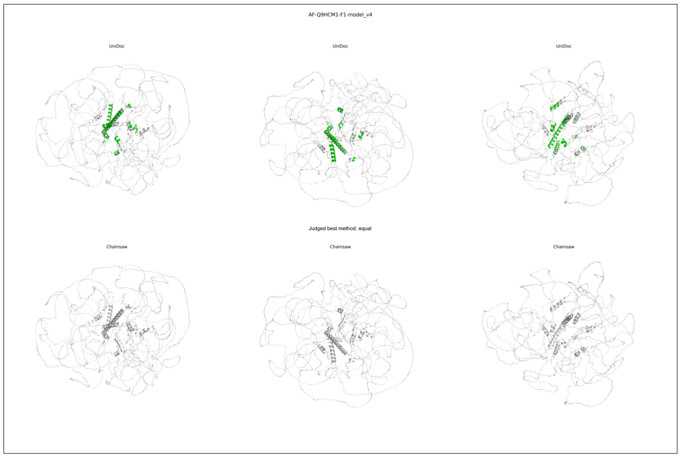
AF-Q9HCM1-F1-model_v4
UniDoc
UniDoc
UniDoc
Judged best method: equal
Chainsaw
Chainsaw
Chainsaw

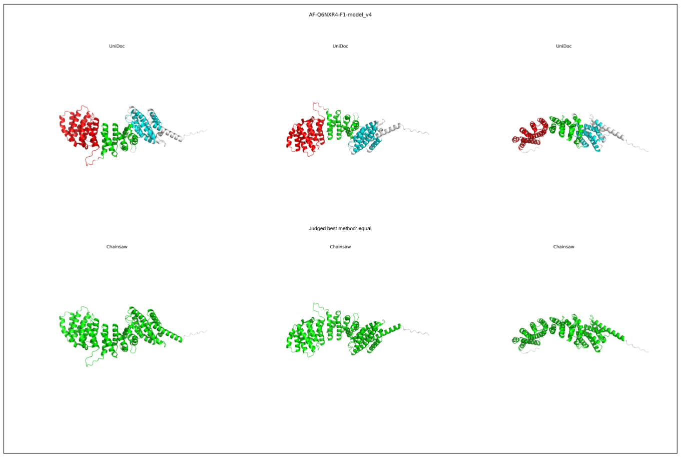
AF-Q6NXR4-F1-model_v4
UniDoc
UniDoc
UniDoc
Judged best method: equal
Chainsaw
Chainsaw
Chainsaw

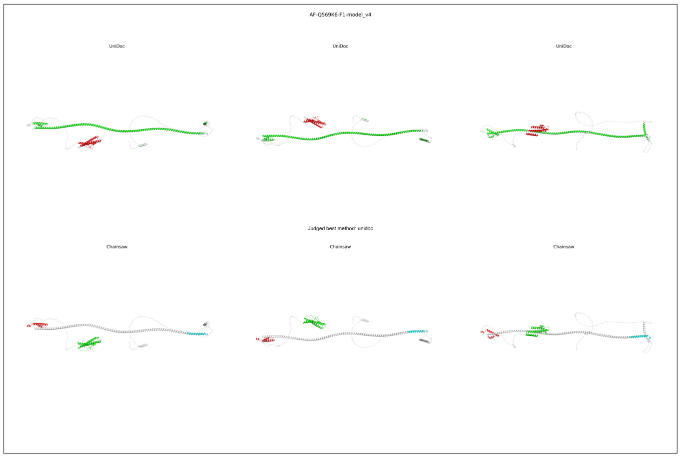
AF-Q569K6-F1-model_v4
UniDoc
UniDoc
UniDoc
Judged best method: unidoc
Chainsaw
Chainsaw
Chainsaw

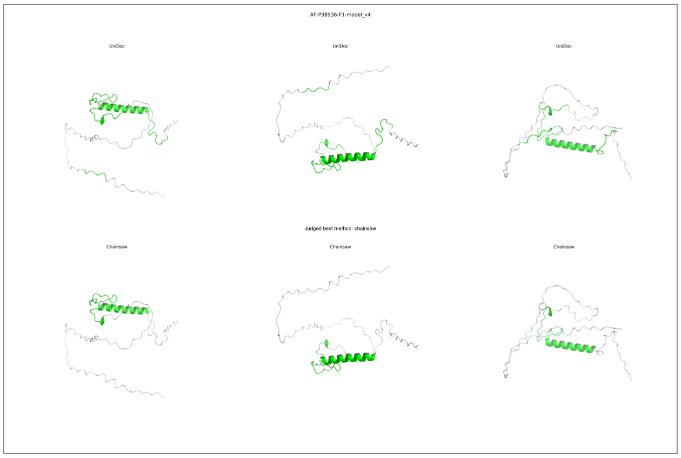
AF-P38936-F1-model_v4
UniDoc
UniDoc
UniDoc
Judged best method: chainsaw
Chainsaw
Chainsaw
Chainsaw

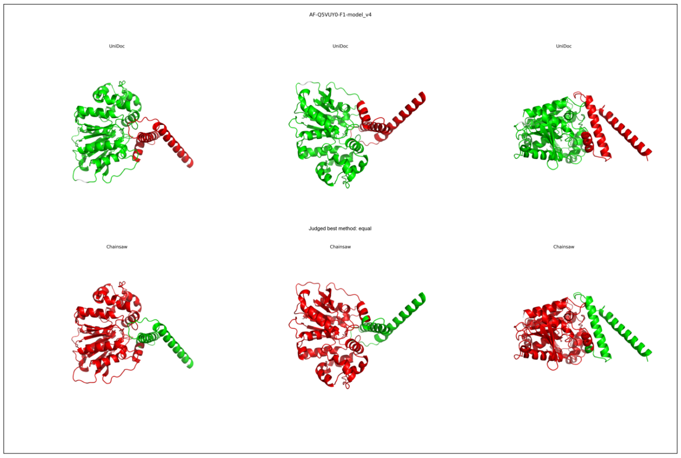
AF-Q5VUY0-F1-model_v4
UniDoc
UniDoc
UniDoc
Judged best method: equal
Chainsaw
Chainsaw
Chainsaw

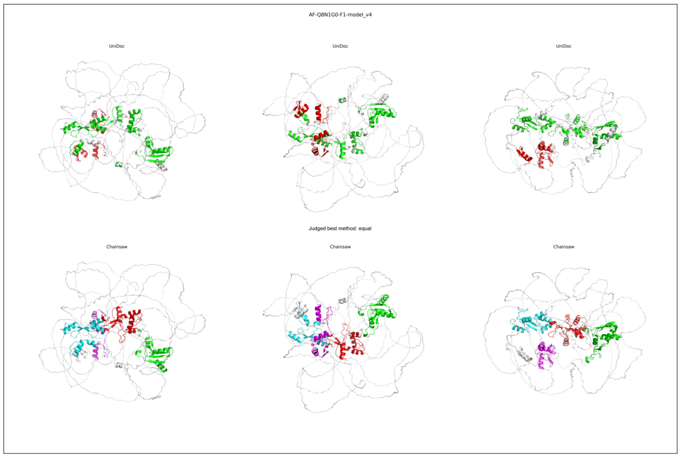
AF-Q8N1G0-F1-model_v4
UniDoc
UniDoc
UniDoc
Judged best method: equal
Chainsaw
Chainsaw
Chainsaw

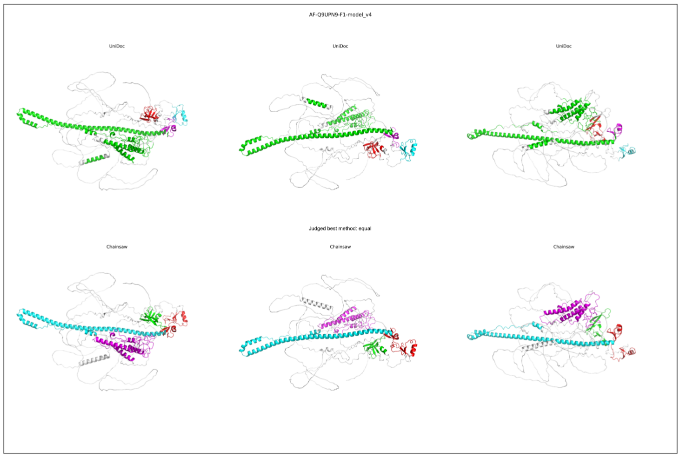
AF-Q9UPN9-F1-model_v4
UniDoc
UniDoc
UniDoc
Judged best method: equal
Chainsaw
Chainsaw
Chainsaw

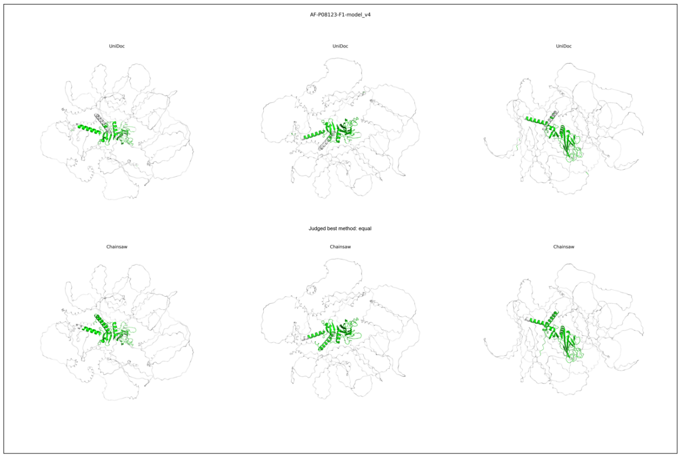
AF-P08123-F1-model_v4
UniDoc
UniDoc
UniDoc
Judged best method: equal
Chainsaw
Chainsaw
Chainsaw

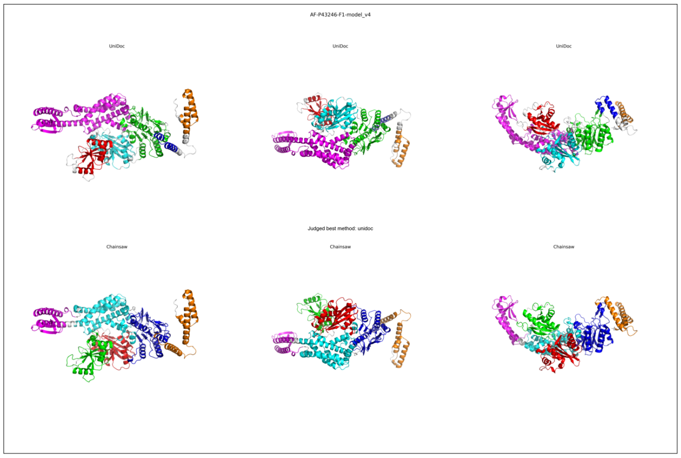
AF-P43246-F1-model_v4
UniDoc
UniDoc
UniDoc
Judged best method: unidoc
Chainsaw
Chainsaw
Chainsaw

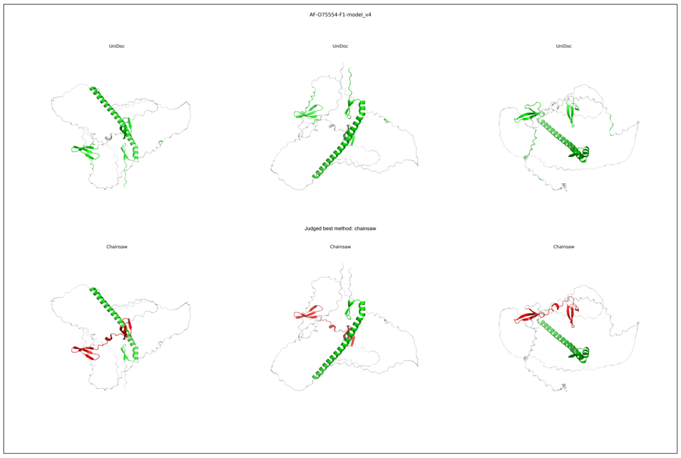
AF-O75554-F1-model_v4
UniDoc
UniDoc
UniDoc
Judged best method: chainsaw
Chainsaw
Chainsaw
Chainsaw

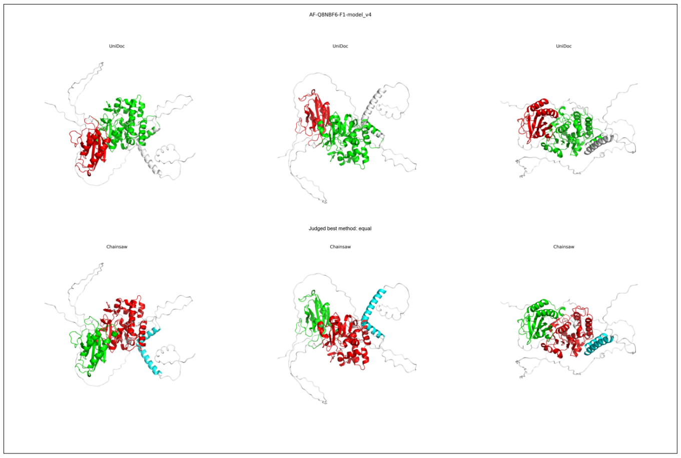
AF-Q8NBF6-F1-model_v4
UniDoc
UniDoc
UniDoc
Judged best method: equal
Chainsaw
Chainsaw
Chainsaw

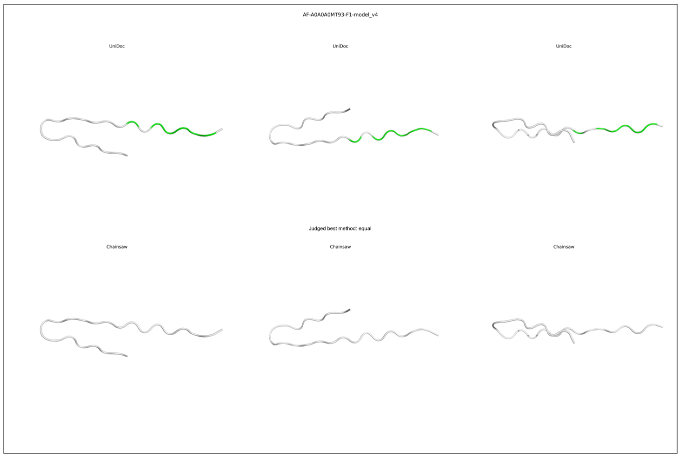
AF-A0A0A0MT93-F1-model_v4
UniDoc
UniDoc
UniDoc
Judged best method: equal
Chainsaw
Chainsaw
Chainsaw

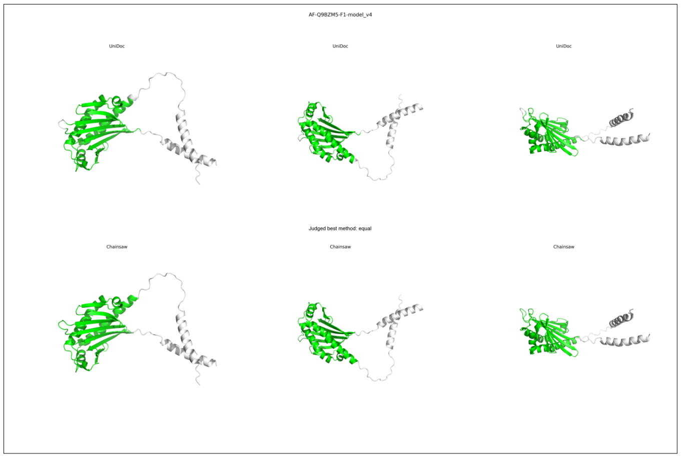
AF-Q9BZM5-F1-model_v4
UniDoc
UniDoc
UniDoc
Judged best method: equal
Chainsaw
Chainsaw
Chainsaw

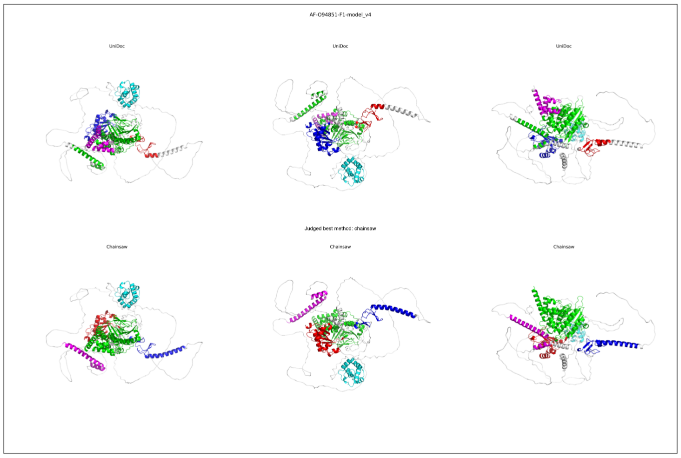
AF-O94851-F1-model_v4
UniDoc
UniDoc
UniDoc
Judged best method: chainsaw
Chainsaw
Chainsaw
Chainsaw

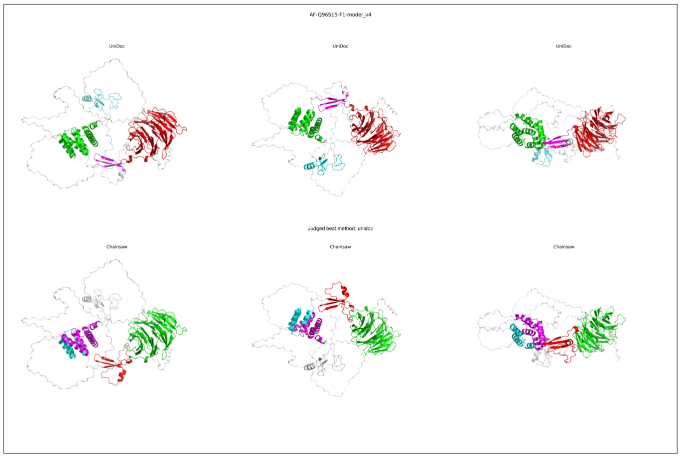
AF-Q96S15-F1-model_v4
UniDoc
UniDoc
UniDoc
Judged best method: unidoc
Chainsaw
Chainsaw
Chainsaw

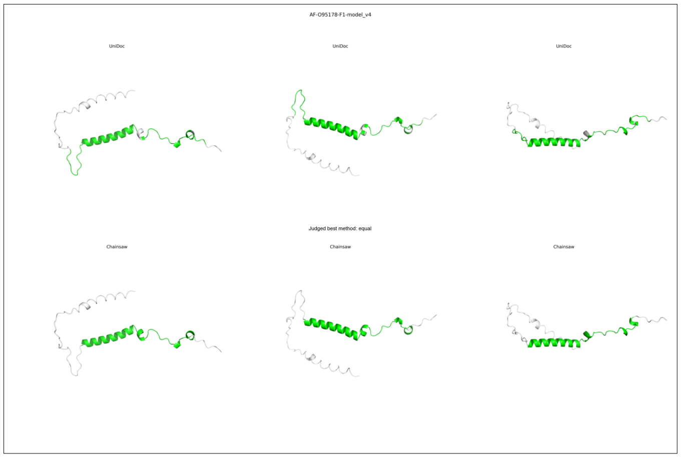
AF-O95178-F1-model_v4
UniDoc
UniDoc
UniDoc
Judged best method: equal
Chainsaw
Chainsaw
Chainsaw

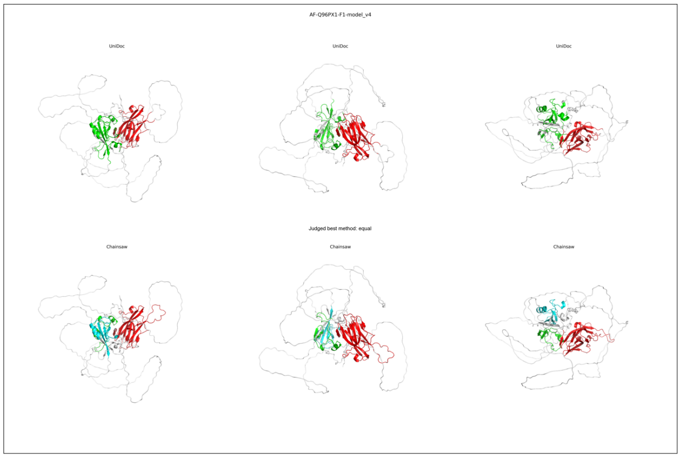
AF-Q96PX1-F1-model_v4
UniDoc
UniDoc
UniDoc
Judged best method: equal
Chainsaw
Chainsaw
Chainsaw

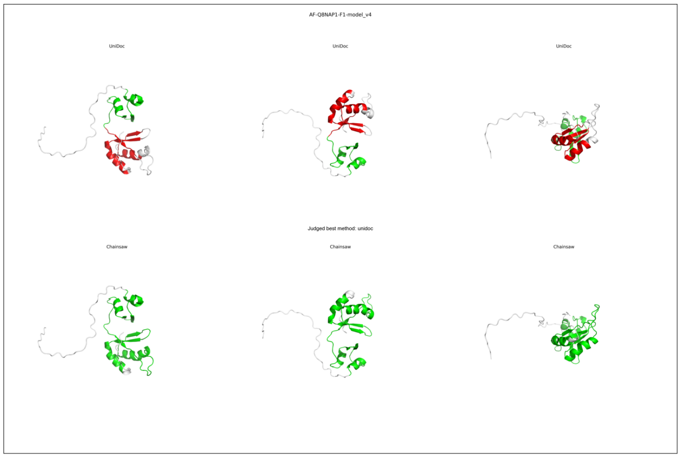
AF-Q8NAP1-F1-model_v4
UniDoc
UniDoc
UniDoc
Judged best method: unidoc
Chainsaw
Chainsaw
Chainsaw

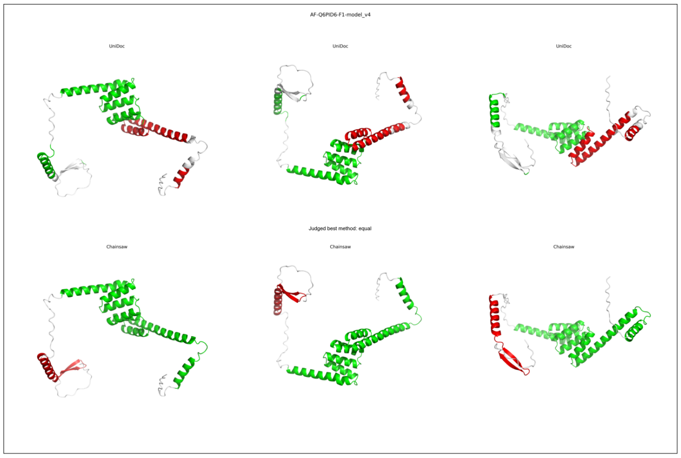
AF-Q6PID6-F1-model_v4
UniDoc
UniDoc
UniDoc
Judged best method: equal
Chainsaw
Chainsaw
Chainsaw

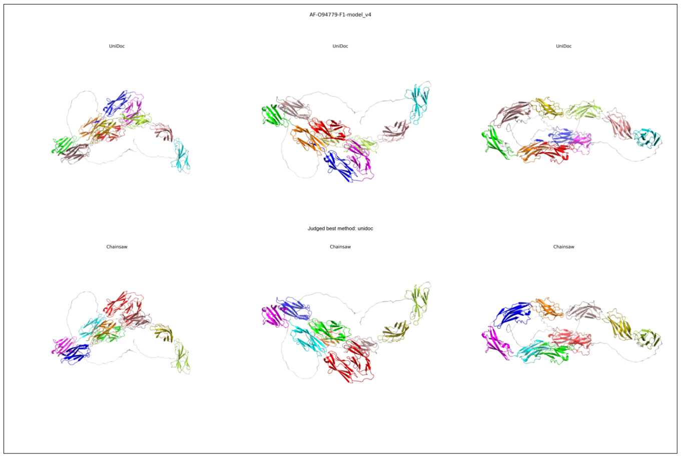
AF-O94779-F1-model_v4
UniDoc
UniDoc
UniDoc
Judged best method: unidoc
Chainsaw
Chainsaw
Chainsaw

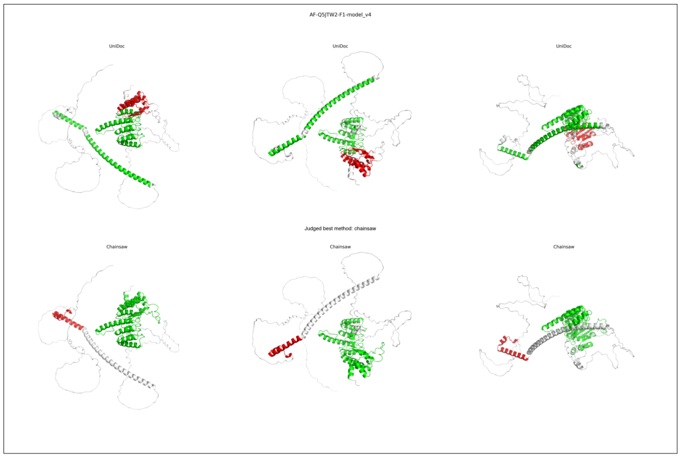
AF-Q5JTW2-F1-model_v4
UniDoc
UniDoc
UniDoc
Judged best method: chainsaw
Chainsaw
Chainsaw
Chainsaw

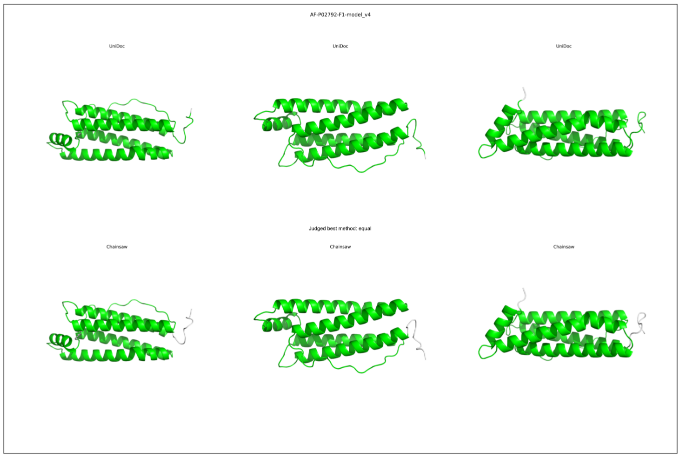
AF-P02792-F1-model_v4
UniDoc
UniDoc
UniDoc
Judged best method: equal
Chainsaw
Chainsaw
Chainsaw

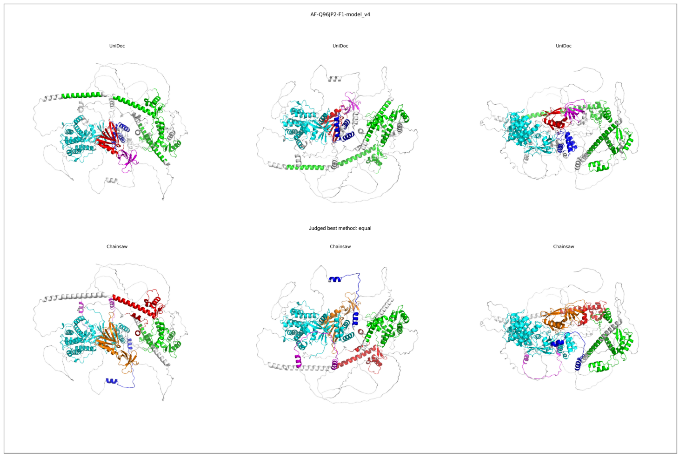
AF-Q96JP2-F1-model_v4
UniDoc
UniDoc
UniDoc
Judged best method: equal
Chainsaw
Chainsaw
Chainsaw

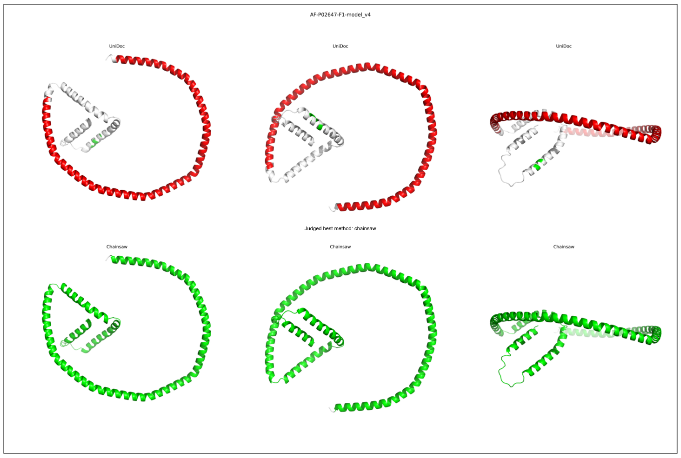
AF-P02647-F1-model_v4
UniDoc
UniDoc
UniDoc
Judged best method: chainsaw
Chainsaw
Chainsaw
Chainsaw

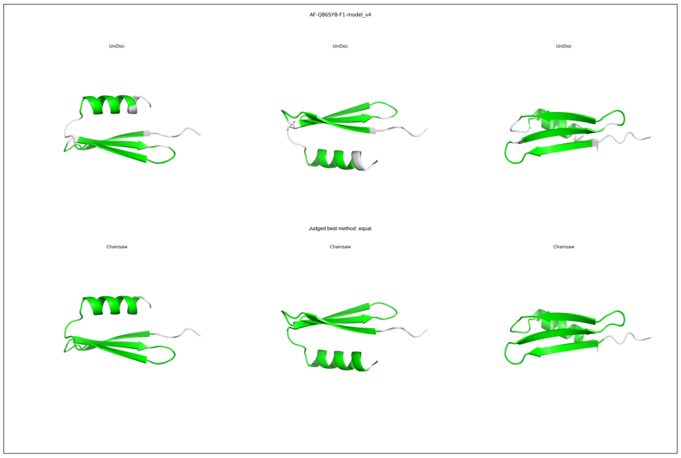
AF-Q86SY8-F1-model_v4
UniDoc
UniDoc
UniDoc
Judged best method: equal
Chainsaw
Chainsaw
Chainsaw

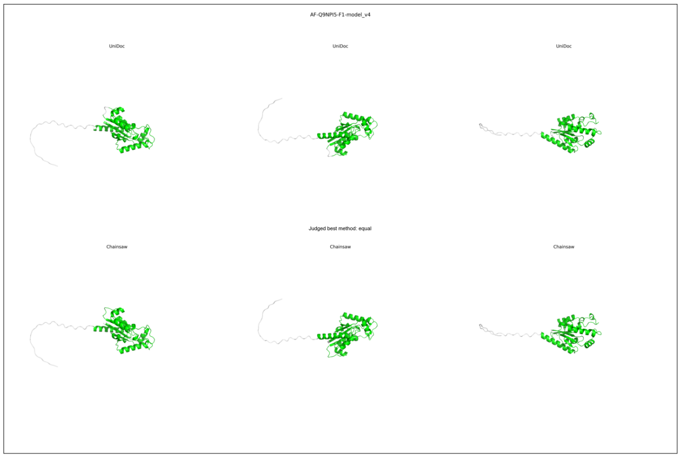
AF-Q9NPI5-F1-model_v4
UniDoc
UniDoc
UniDoc
Judged best method: equal
Chainsaw
Chainsaw
Chainsaw

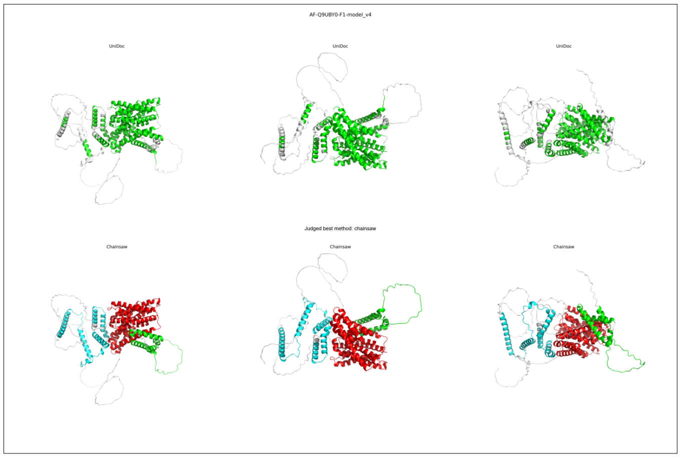
AF-Q9UBY0-F1-model_v4
UniDoc
UniDoc
UniDoc
Judged best method: chainsaw
Chainsaw
Chainsaw
Chainsaw

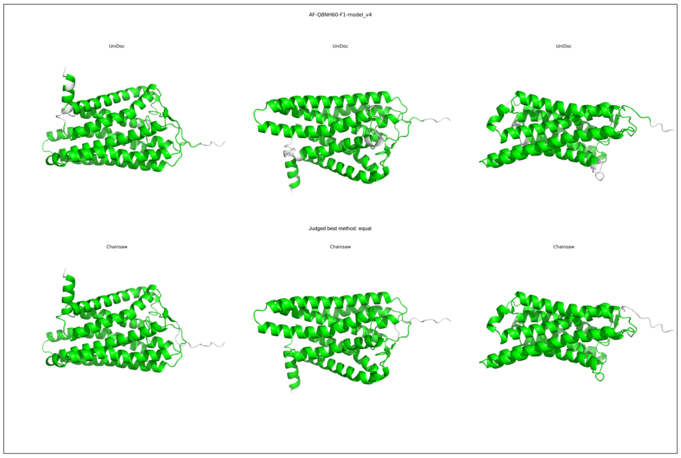
AF-Q8NH60-F1-model_v4
UniDoc
UniDoc
UniDoc
Judged best method: equal
Chainsaw
Chainsaw
Chainsaw

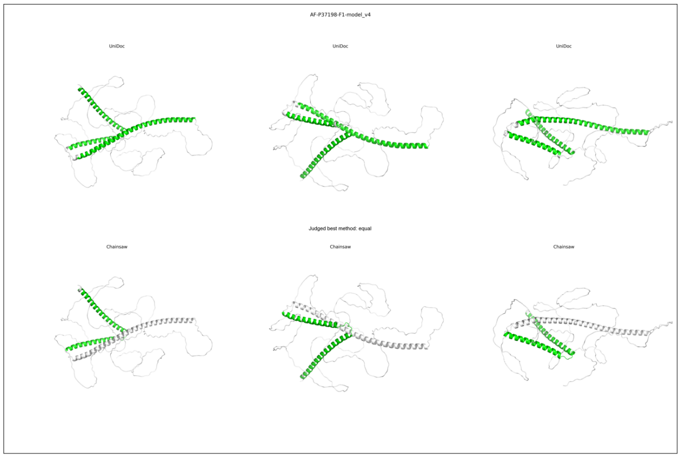
AF-P37198-F1-model_v4
UniDoc
UniDoc
UniDoc
Judged best method: equal
Chainsaw
Chainsaw
Chainsaw

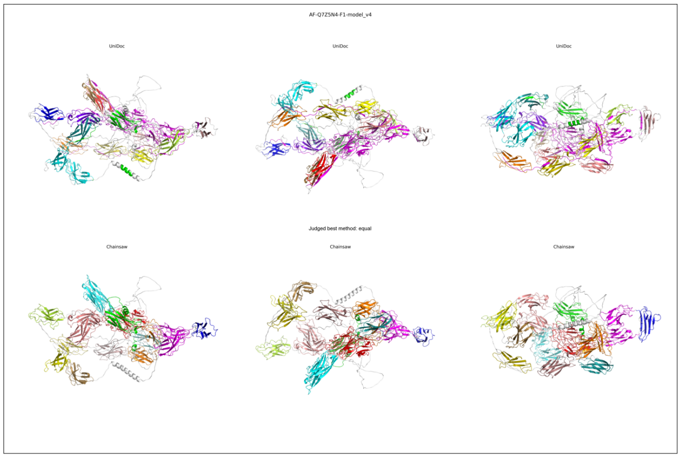
AF-Q7Z5N4-F1-model_v4
UniDoc
UniDoc
UniDoc
Judged best method: equal
Chainsaw
Chainsaw
Chainsaw

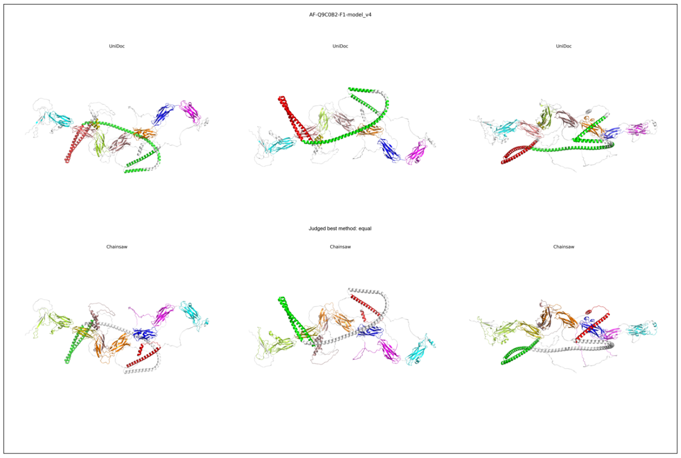
AF-Q9C0B2-F1-model_v4
UniDoc
UniDoc
UniDoc
Judged best method: equal
Chainsaw
Chainsaw
Chainsaw

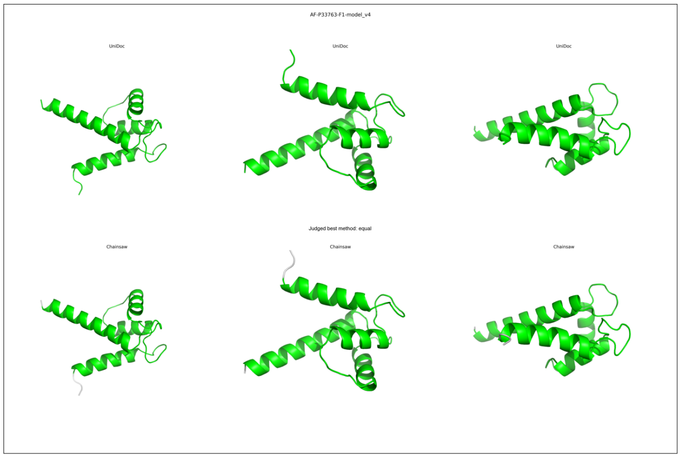
AF-P33763-F1-model_v4
UniDoc
UniDoc
UniDoc
Judged best method: equal
Chainsaw
Chainsaw
Chainsaw

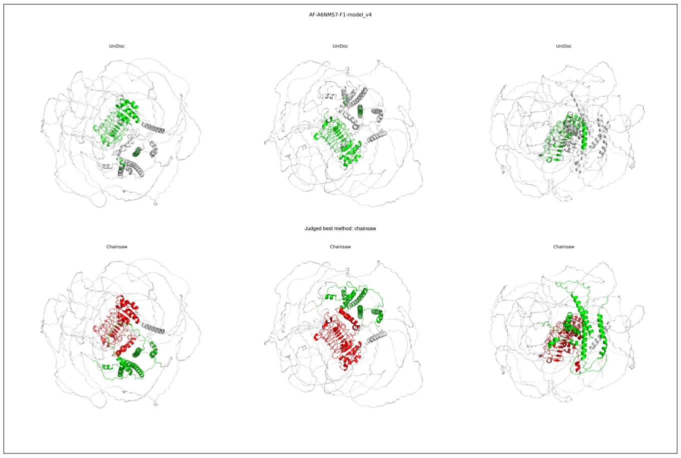
AF-A6NMS7-F1-model_v4
UniDoc
UniDoc
UniDoc
Judged best method: chainsaw
Chainsaw
Chainsaw
Chainsaw

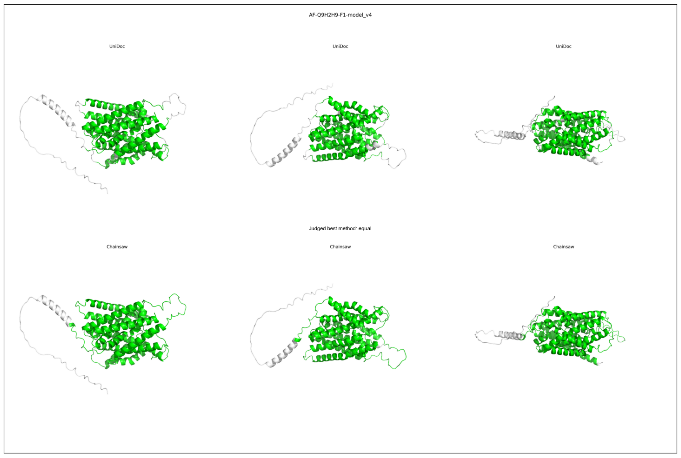
AF-Q9H2H9-F1-model_v4
UniDoc
UniDoc
UniDoc
Judged best method: equal
Chainsaw
Chainsaw
Chainsaw

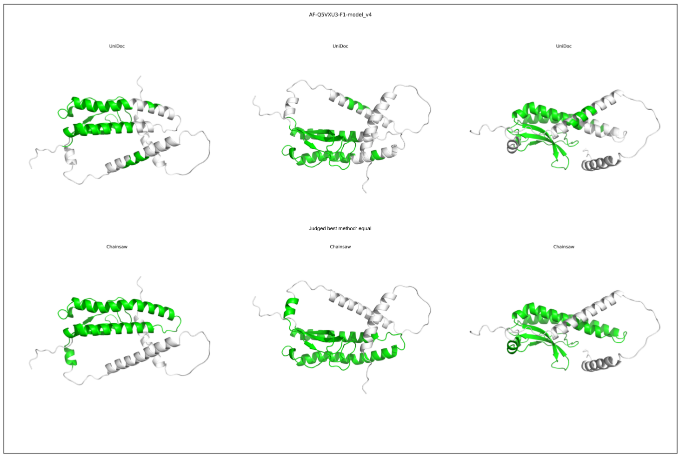
AF-Q5VXU3-F1-model_v4
UniDoc
UniDoc
UniDoc
Judged best method: equal
Chainsaw
Chainsaw
Chainsaw

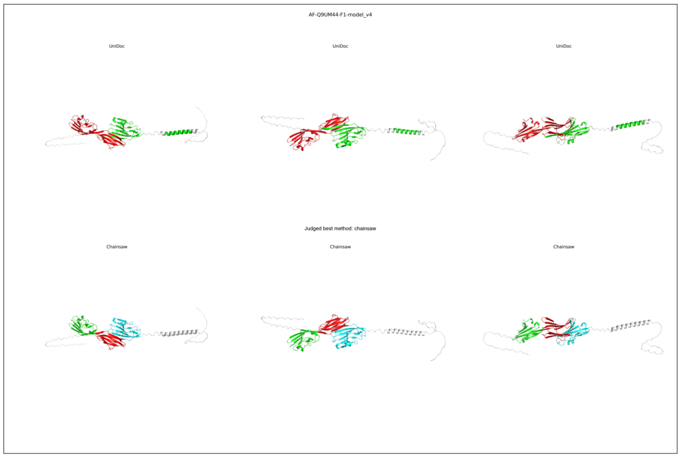
AF-Q9UM44-F1-model_v4
UniDoc
UniDoc
UniDoc
Judged best method: chainsaw
Chainsaw
Chainsaw
Chainsaw

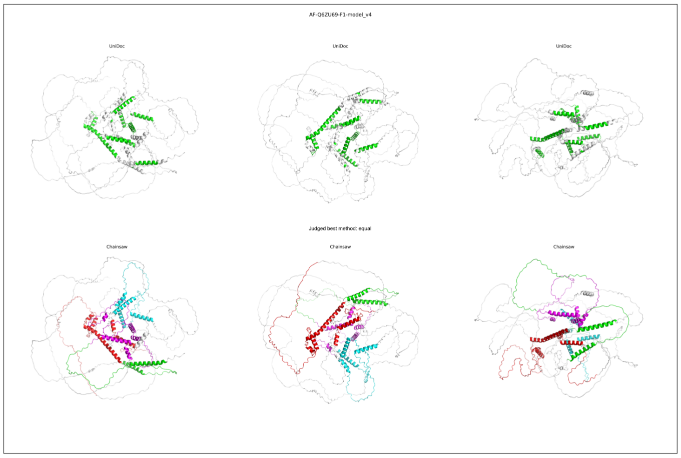
AF-Q6ZU69-F1-model_v4
UniDoc
UniDoc
UniDoc
Judged best method: equal
Chainsaw
Chainsaw
Chainsaw

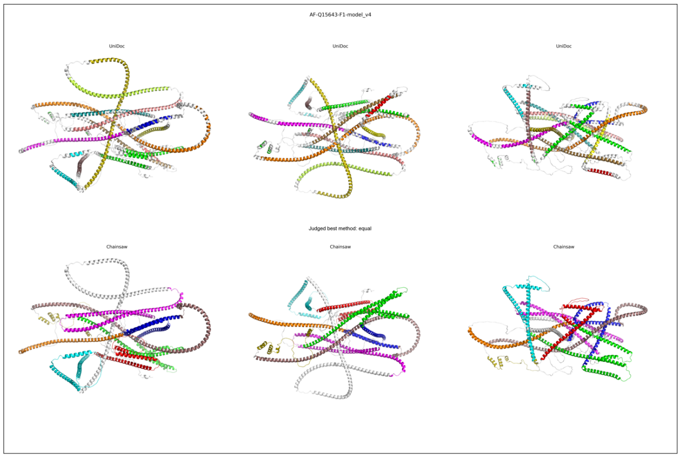
AF-Q15643-F1-model_v4
UniDoc
UniDoc
UniDoc
Judged best method: equal
Chainsaw
Chainsaw
Chainsaw

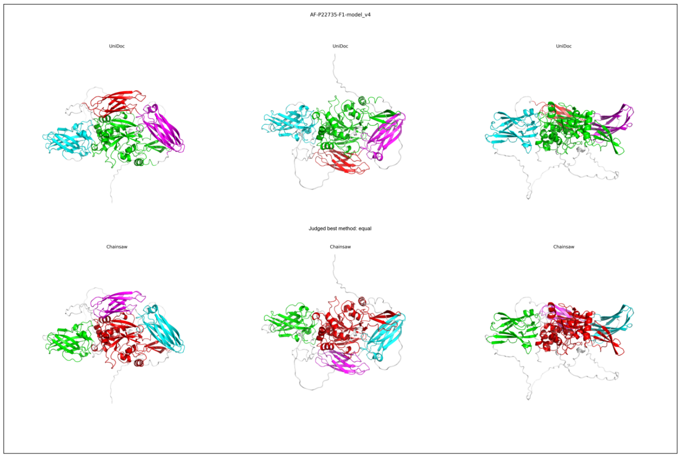
AF-P22735-F1-model_v4
UniDoc
UniDoc
UniDoc
Judged best method: equal
Chainsaw
Chainsaw
Chainsaw

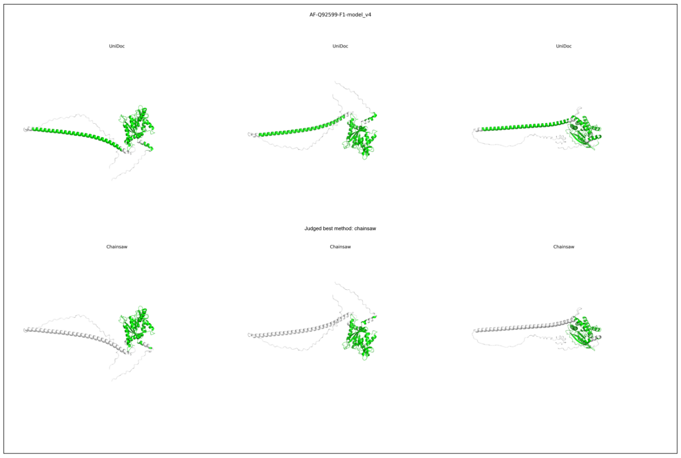
AF-Q92599-F1-model_v4
UniDoc
UniDoc
UniDoc
Judged best method: chainsaw
Chainsaw
Chainsaw
Chainsaw

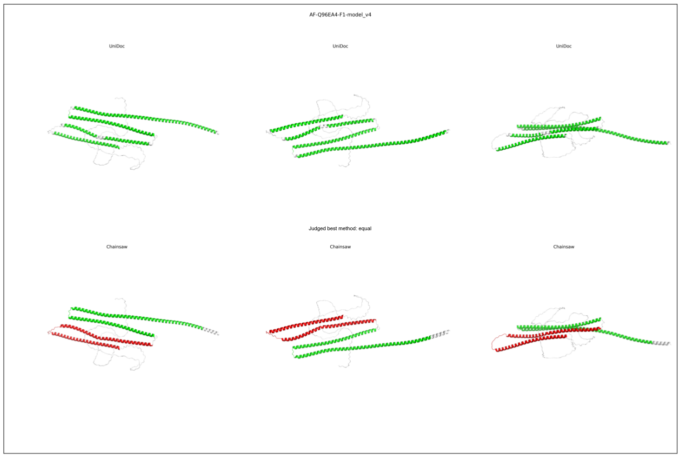
AF-Q96EA4-F1-model_v4
UniDoc
UniDoc
UniDoc
Judged best method: equal
Chainsaw
Chainsaw
Chainsaw

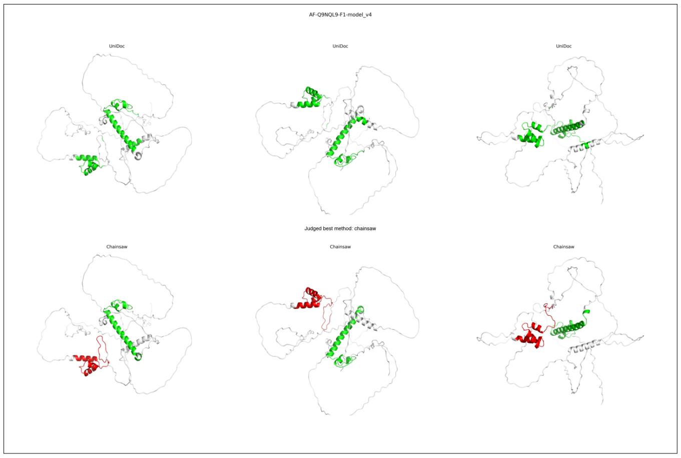
AF-Q9NQL9-F1-model_v4
UniDoc
UniDoc
UniDoc
Judged best method: chainsaw
Chainsaw
Chainsaw
Chainsaw

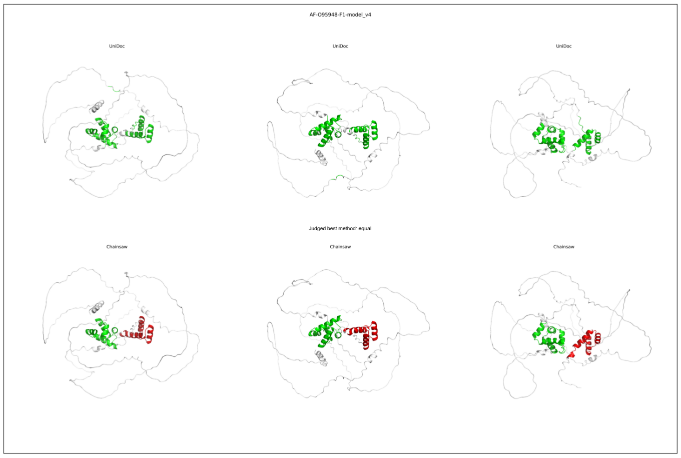
AF-O95948-F1-model_v4
UniDoc
UniDoc
UniDoc
Judged best method: equal
Chainsaw
Chainsaw
Chainsaw

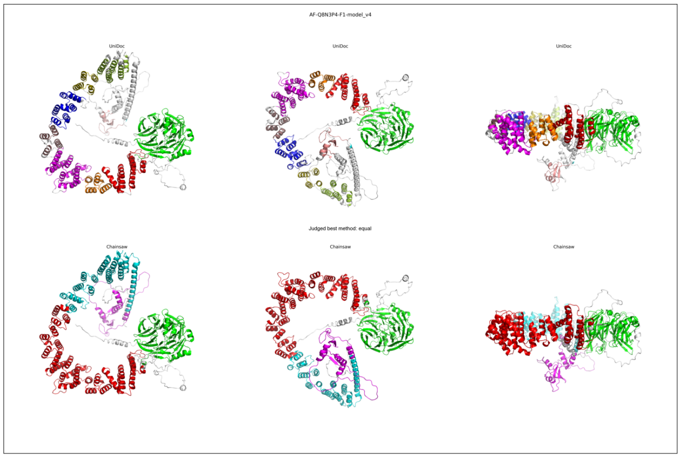
AF-Q8N3P4-F1-model_v4
UniDoc
UniDoc
UniDoc
Judged best method: equal
Chainsaw
Chainsaw
Chainsaw

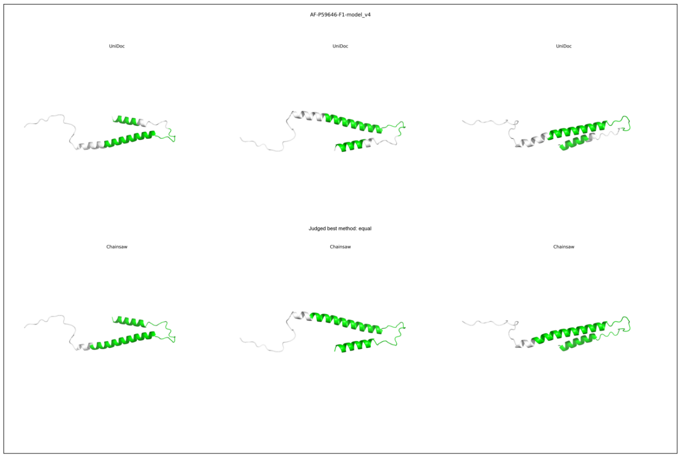
AF-P59646-F1-model_v4
UniDoc
UniDoc
UniDoc
Judged best method: equal
Chainsaw
Chainsaw
Chainsaw

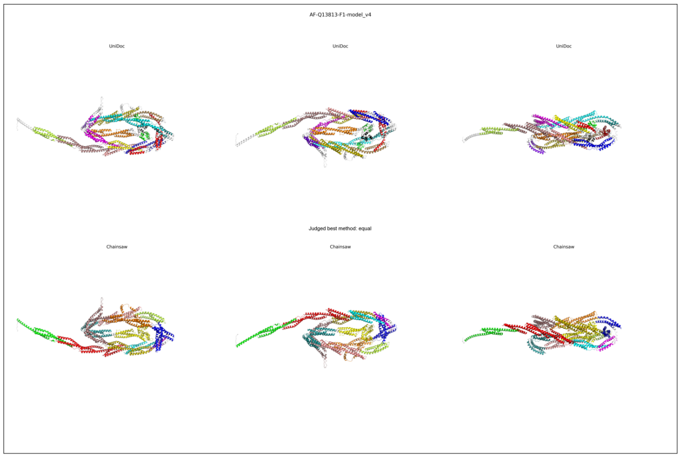
AF-Q13813-F1-model_v4
UniDoc
UniDoc
UniDoc
Judged best method: equal
Chainsaw
Chainsaw
Chainsaw

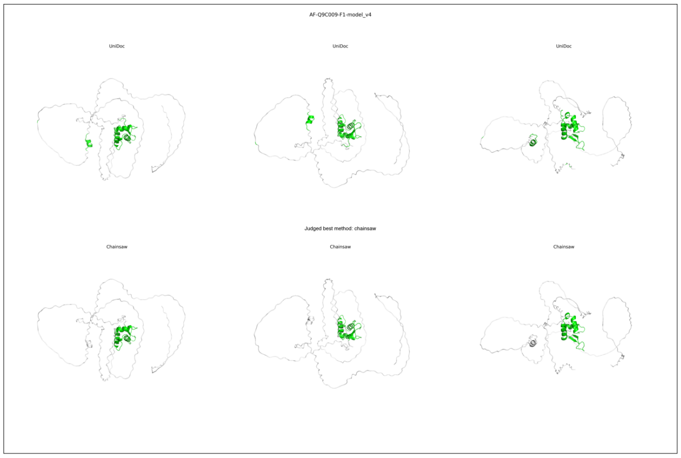
AF-Q9C009-F1-model_v4
UniDoc
UniDoc
UniDoc
Judged best method: chainsaw
Chainsaw
Chainsaw
Chainsaw

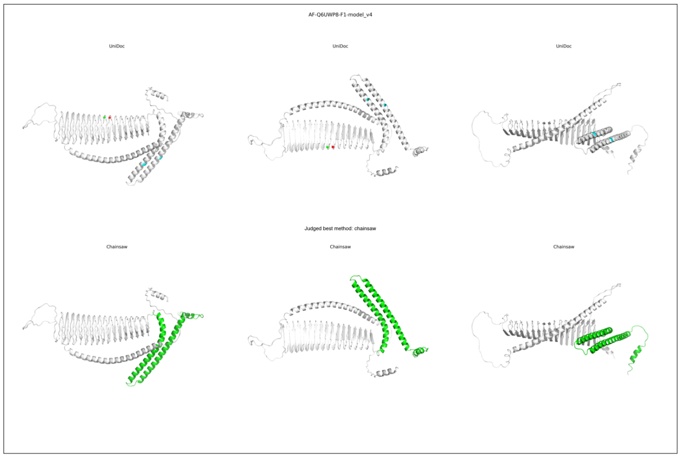
AF-Q6UWP8-F1-model_v4
UniDoc
UniDoc
UniDoc
Judged best method: chainsaw
Chainsaw
Chainsaw
Chainsaw

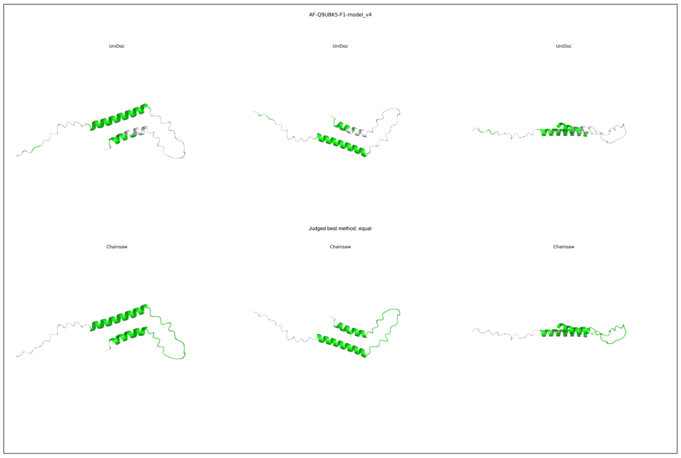
AF-Q9UBK5-F1-model_v4
UniDoc
UniDoc
UniDoc
Judged best method: equal
Chainsaw
Chainsaw
Chainsaw

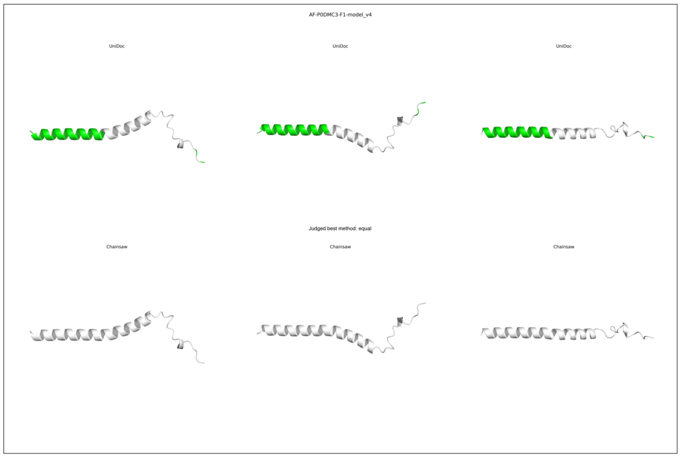
AF-P0DMC3-F1-model_v4
UniDoc
UniDoc
UniDoc
Judged best method: equal
Chainsaw
Chainsaw
Chainsaw

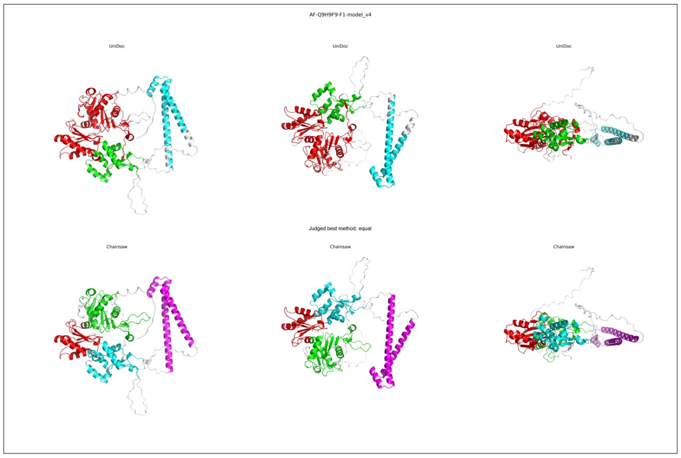
AF-Q9H9F9-F1-model_v4
UniDoc
UniDoc
UniDoc
Judged best method: equal
Chainsaw
Chainsaw
Chainsaw

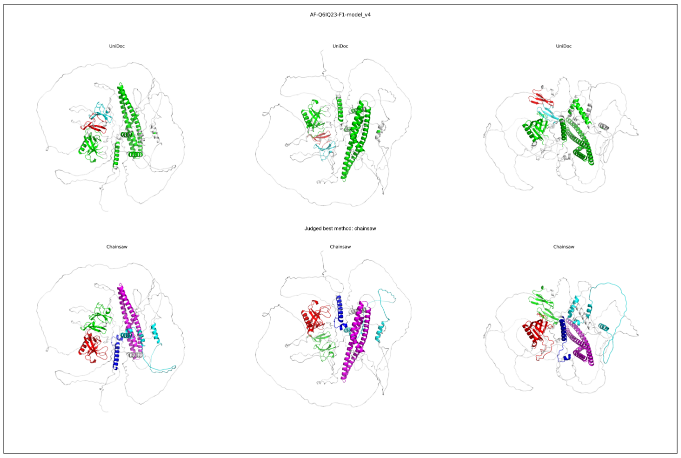
AF-Q6IQ23-F1-model_v4
UniDoc
UniDoc
UniDoc
Judged best method: chainsaw
Chainsaw
Chainsaw
Chainsaw

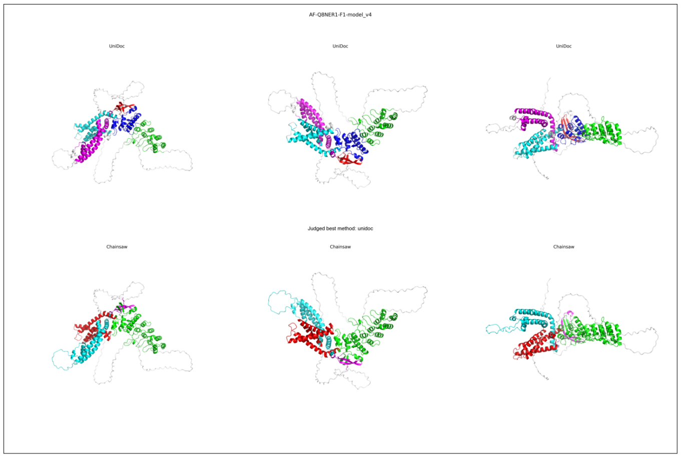
AF-Q8NER1-F1-model_v4
UniDoc
UniDoc
UniDoc
Judged best method: unidoc
Chainsaw
Chainsaw
Chainsaw

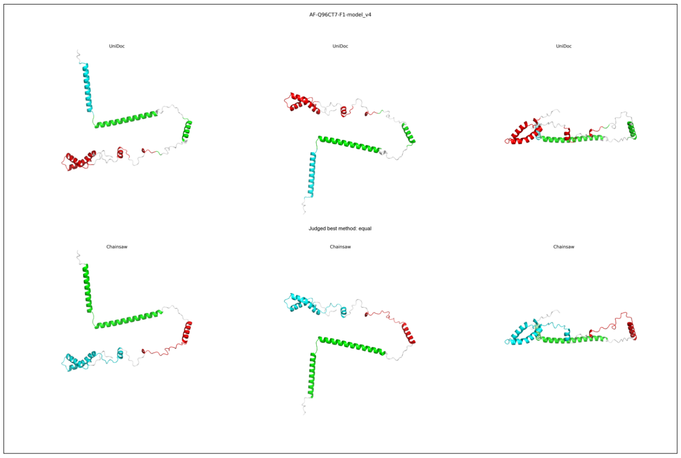
AF-Q96CT7-F1-model_v4
UniDoc
UniDoc
UniDoc
Judged best method: equal
Chainsaw
Chainsaw
Chainsaw

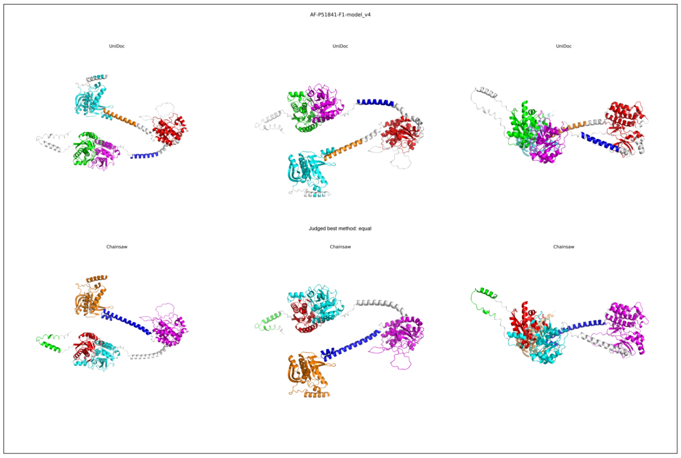
AF-P51841-F1-model_v4
UniDoc
UniDoc
UniDoc
Judged best method: equal
Chainsaw
Chainsaw
Chainsaw

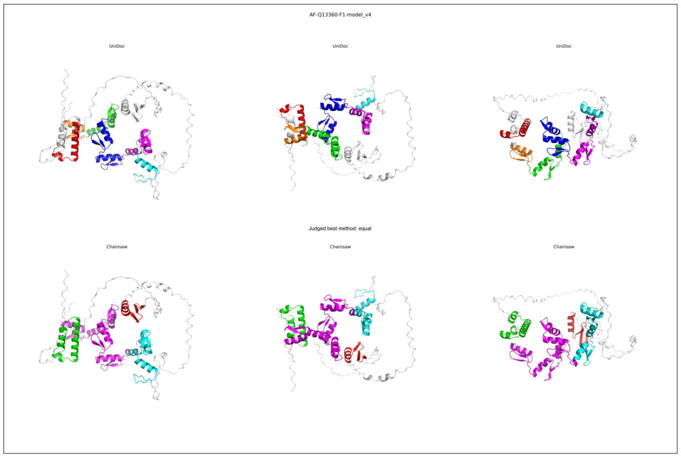
AF-Q13360-F1-model_v4
UniDoc
UniDoc
UniDoc
Judged best method: equal
Chainsaw
Chainsaw
Chainsaw

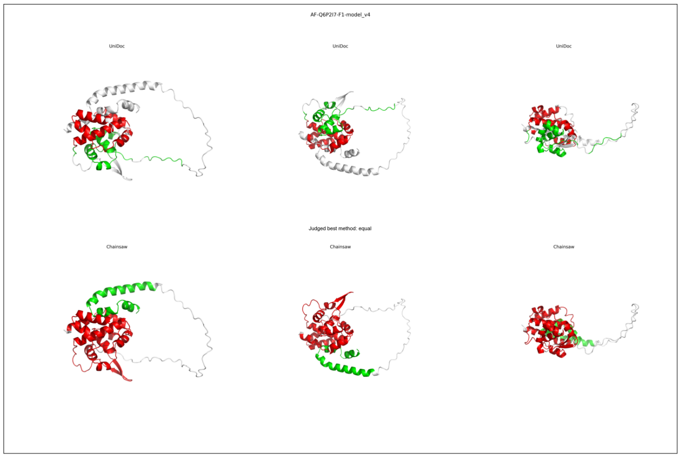
AF-Q6P2I7-F1-model_v4
UniDoc
UniDoc
UniDoc
Judged best method: equal
Chainsaw
Chainsaw
Chainsaw

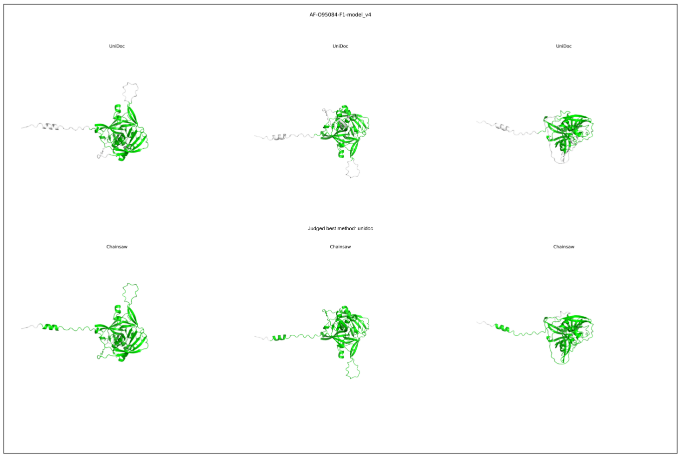
AF-O95084-F1-model_v4
UniDoc
UniDoc
UniDoc
Judged best method: unidoc
Chainsaw
Chainsaw
Chainsaw

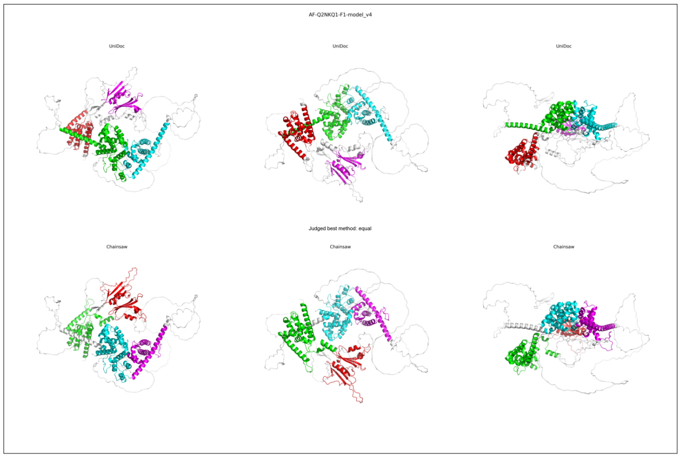
AF-Q2NKQ1-F1-model_v4
UniDoc
UniDoc
UniDoc
Judged best method: equal
Chainsaw
Chainsaw
Chainsaw

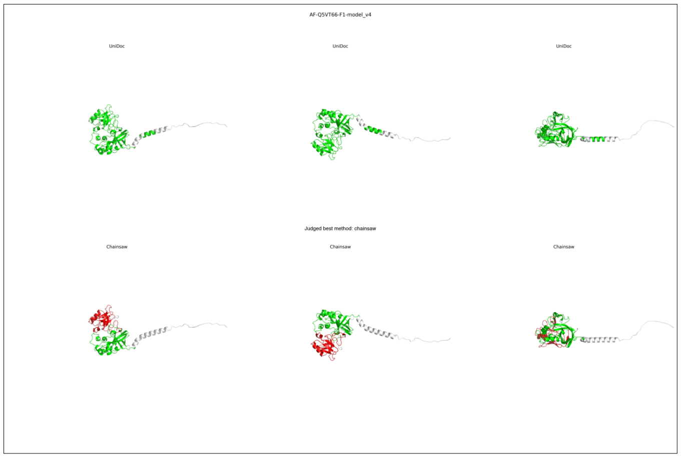
AF-Q5VT66-F1-model_v4
UniDoc
UniDoc
UniDoc
Judged best method: chainsaw
Chainsaw
Chainsaw
Chainsaw

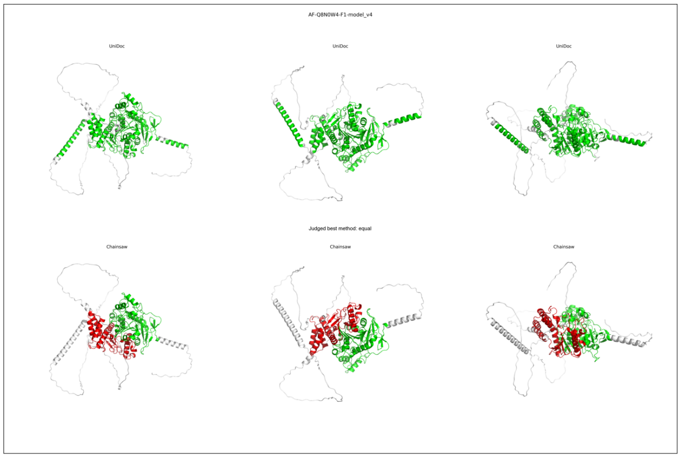
AF-Q8N0W4-F1-model_v4
UniDoc
UniDoc
UniDoc
Judged best method: equal
Chainsaw
Chainsaw
Chainsaw

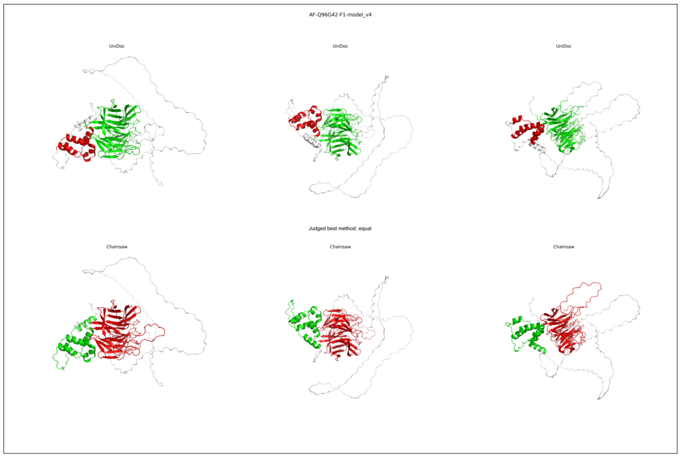
AF-Q96G42-F1-model_v4
UniDoc
UniDoc
UniDoc
Judged best method: equal
Chainsaw
Chainsaw
Chainsaw

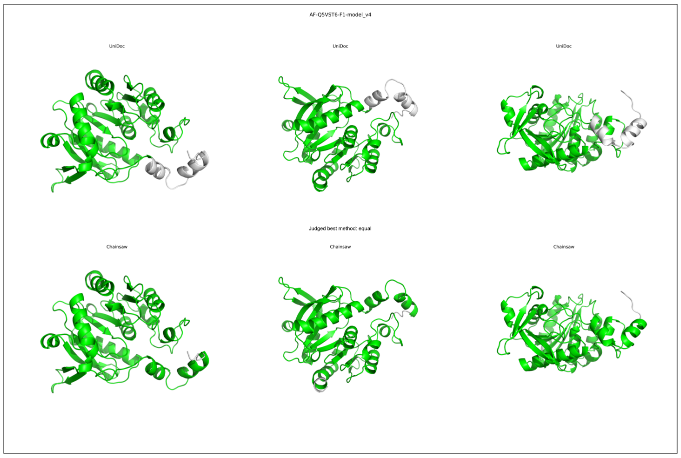
AF-Q5VST6-F1-model_v4
UniDoc
UniDoc
UniDoc
Judged best method: equal
Chainsaw
Chainsaw
Chainsaw

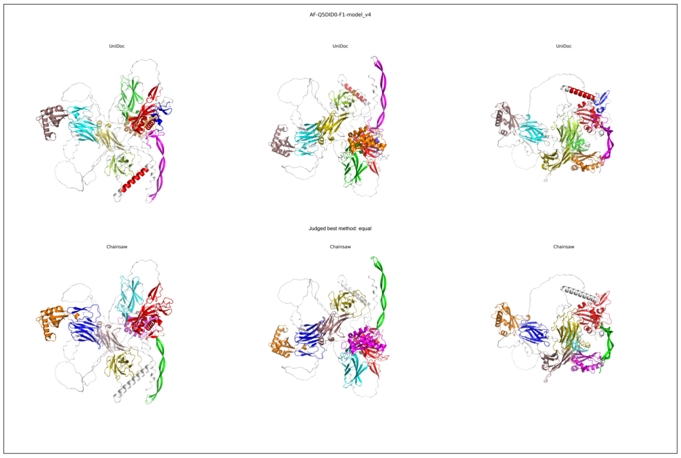
AF-Q5DID0-F1-model_v4
UniDoc
UniDoc
UniDoc
Judged best method: equal
Chainsaw
Chainsaw
Chainsaw

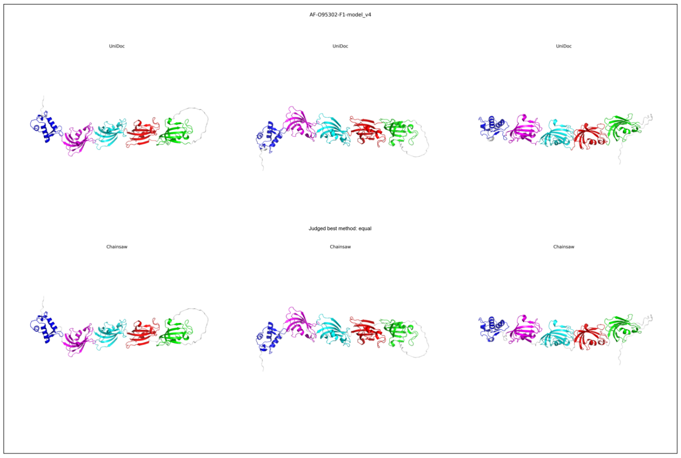
AF-O95302-F1-model_v4
UniDoc
UniDoc
UniDoc
Judged best method: equal
Chainsaw
Chainsaw
Chainsaw

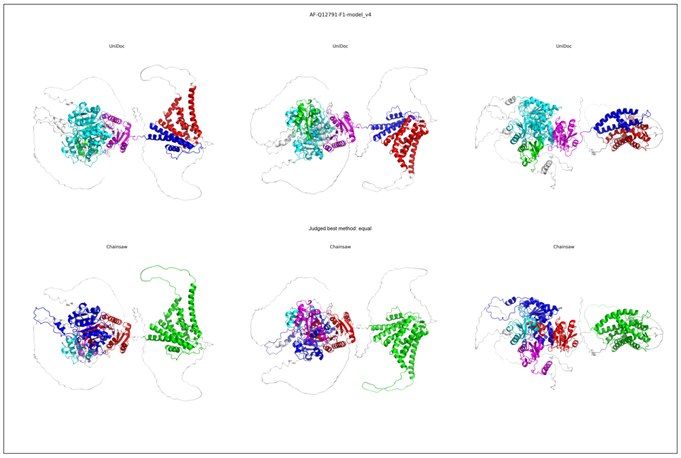
AF-Q12791-F1-model_v4
UniDoc
UniDoc
UniDoc
Judged best method: equal
Chainsaw
Chainsaw
Chainsaw

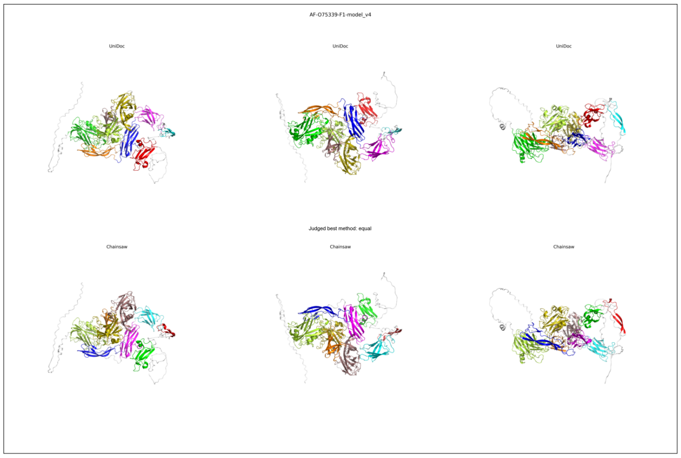
AF-O75339-F1-model_v4
UniDoc
UniDoc
UniDoc
Judged best method: equal
Chainsaw
Chainsaw
Chainsaw

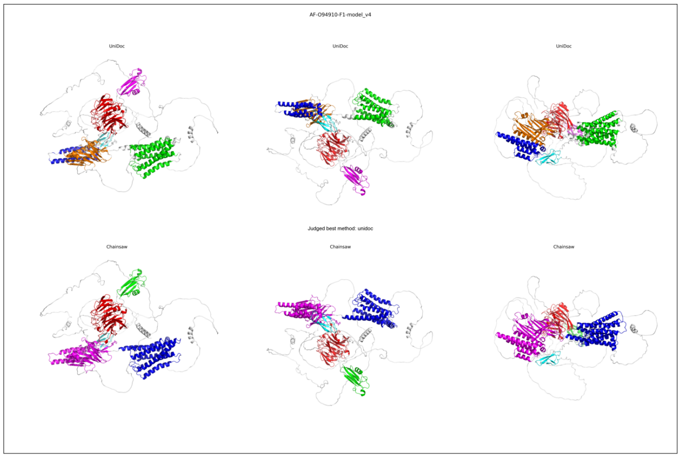
AF-O94910-F1-model_v4
UniDoc
UniDoc
UniDoc
Judged best method: unidoc
Chainsaw
Chainsaw
Chainsaw

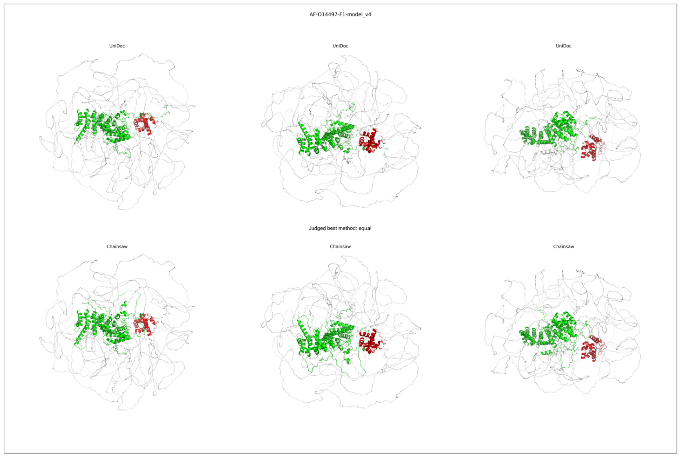
AF-O14497-F1-model_v4
UniDoc
UniDoc
UniDoc
Judged best method: equal
Chainsaw
Chainsaw
Chainsaw

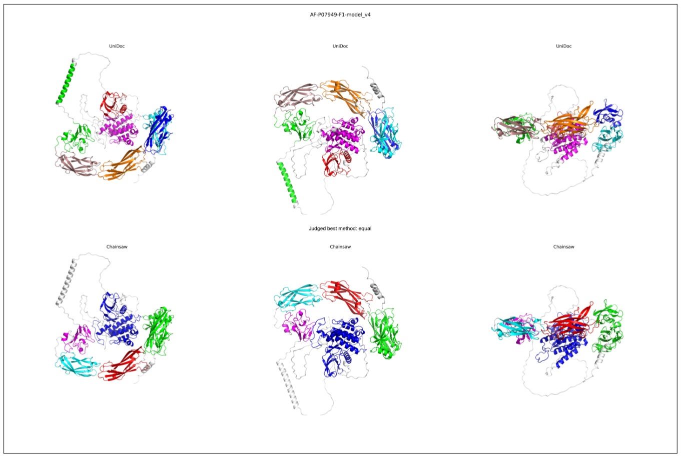
AF-P07949-F1-model_v4
UniDoc
UniDoc
UniDoc
Judged best method: equal
Chainsaw
Chainsaw
Chainsaw

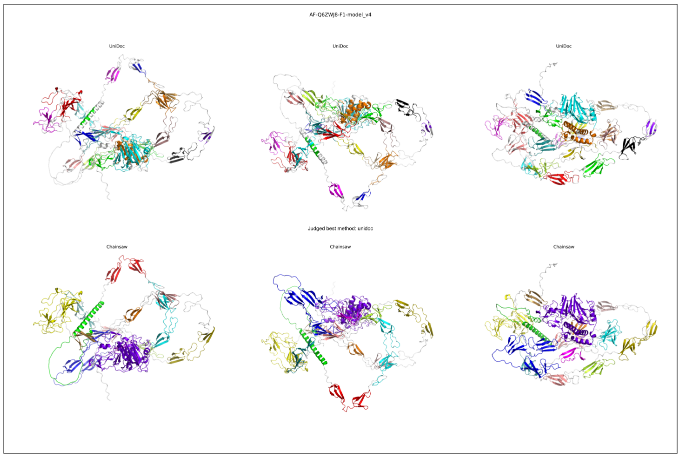
AF-Q6ZWJ8-F1-model_v4
UniDoc
UniDoc
UniDoc
Judged best method: unidoc
Chainsaw
Chainsaw
Chainsaw

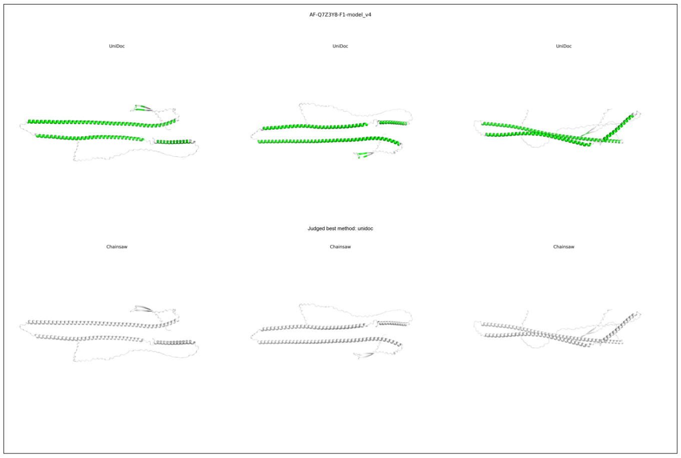
AF-Q7Z3Y8-F1-model_v4
UniDoc
UniDoc
UniDoc
Judged best method: unidoc
Chainsaw
Chainsaw
Chainsaw

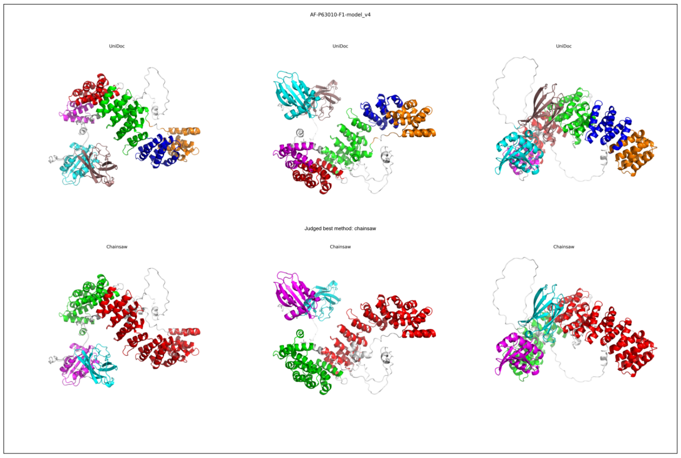
AF-P63010-F1-model_v4
UniDoc
UniDoc
UniDoc
Judged best method: chainsaw
Chainsaw
Chainsaw
Chainsaw

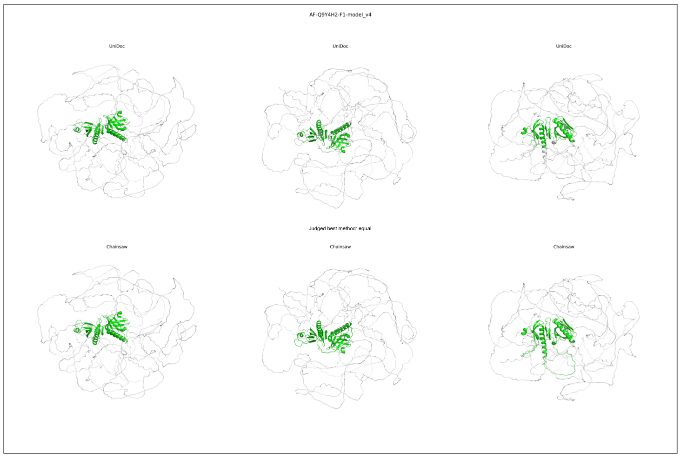
AF-Q9Y4H2-F1-model_v4
UniDoc
UniDoc
UniDoc
Judged best method: equal
Chainsaw
Chainsaw
Chainsaw

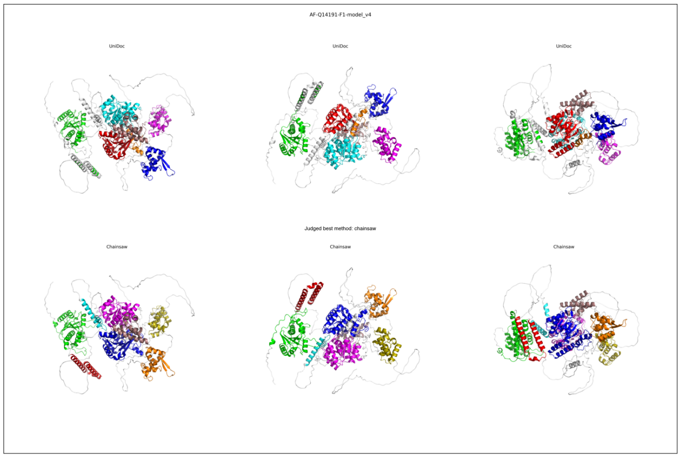
AF-Q14191-F1-model_v4
UniDoc
UniDoc
UniDoc
Judged best method: chainsaw
Chainsaw
Chainsaw
Chainsaw

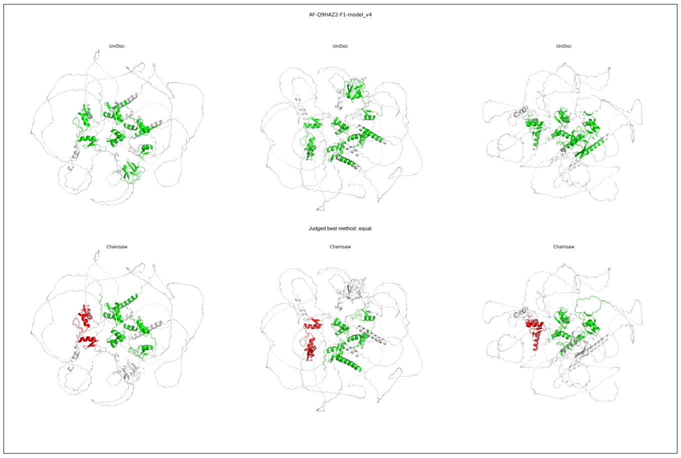
AF-Q9HAZ2-F1-model_v4
UniDoc
UniDoc
UniDoc
Judged best method: equal
Chainsaw
Chainsaw
Chainsaw

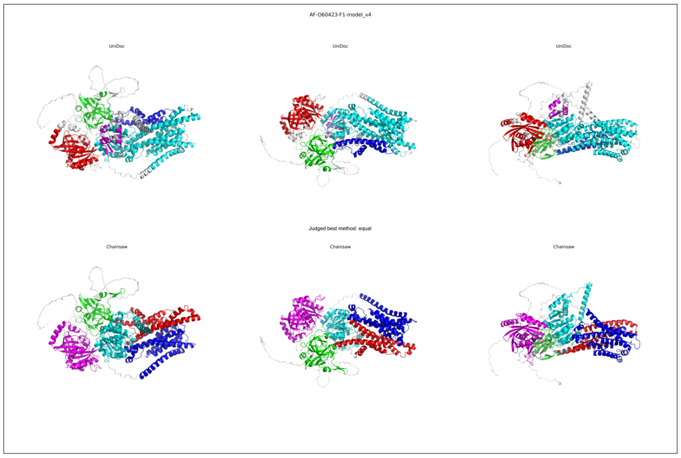
AF-O60423-F1-model_v4
UniDoc
UniDoc
UniDoc
Judged best method: equal
Chainsaw
Chainsaw
Chainsaw

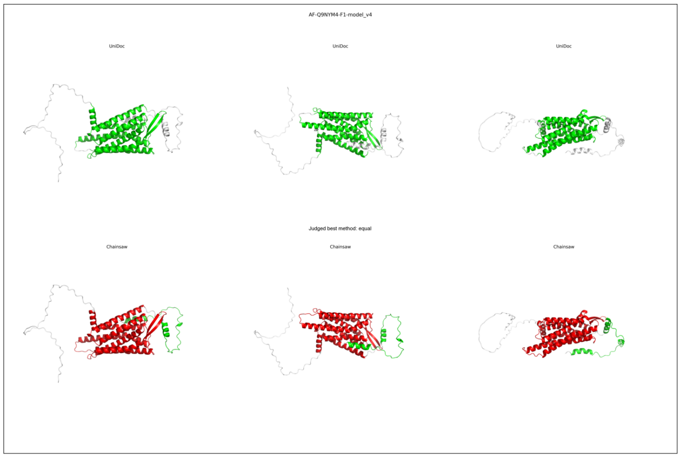
AF-Q9NYM4-F1-model_v4
UniDoc
UniDoc
UniDoc
Judged best method: equal
Chainsaw
Chainsaw
Chainsaw

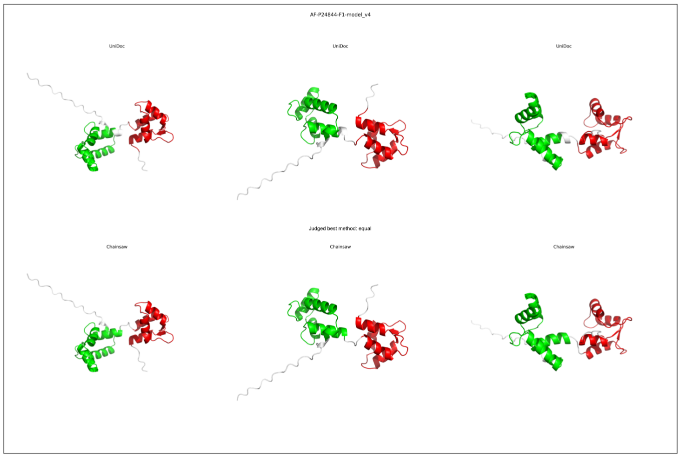
AF-P24844-F1-model_v4
UniDoc
UniDoc
UniDoc
Judged best method: equal
Chainsaw
Chainsaw
Chainsaw

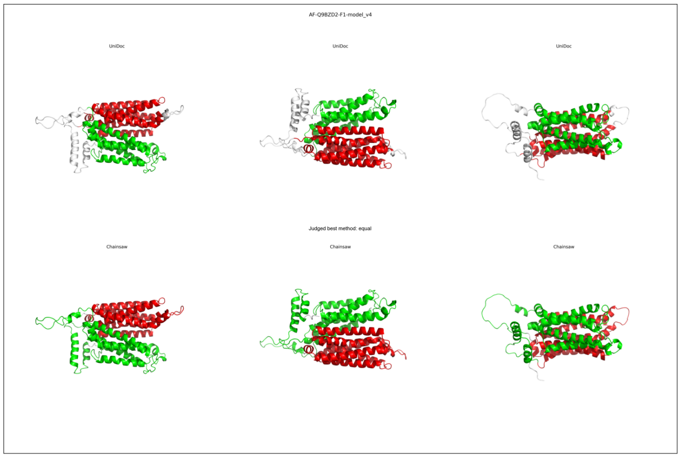
AF-Q9BZD2-F1-model_v4
UniDoc
UniDoc
UniDoc
Judged best method: equal
Chainsaw
Chainsaw
Chainsaw

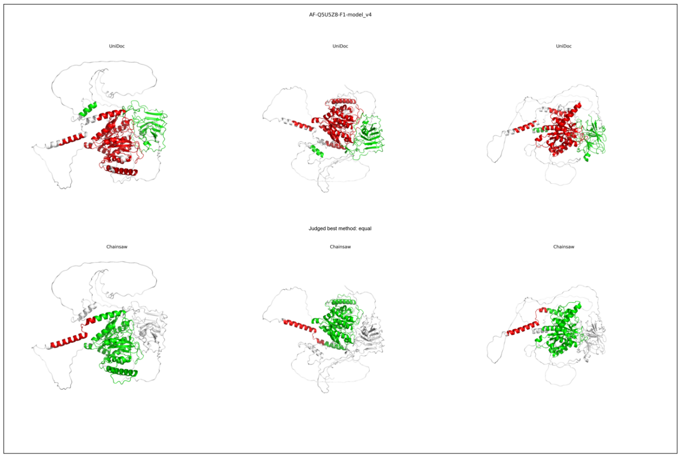
AF-Q5U5Z8-F1-model_v4
UniDoc
UniDoc
UniDoc
Judged best method: equal
Chainsaw
Chainsaw
Chainsaw

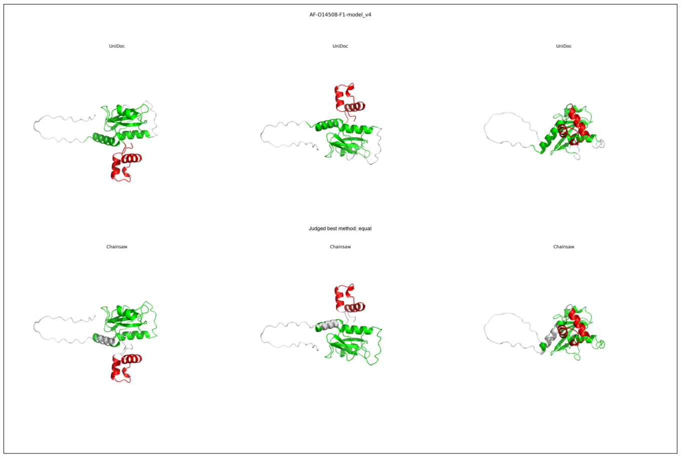
AF-O14508-F1-model_v4
UniDoc
UniDoc
UniDoc
Judged best method: equal
Chainsaw
Chainsaw
Chainsaw

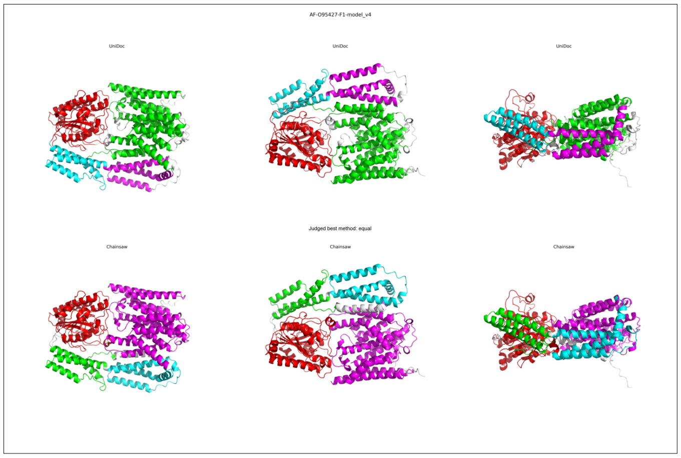
AF-O95427-F1-model_v4
UniDoc
UniDoc
UniDoc
Judged best method: equal
Chainsaw
Chainsaw
Chainsaw

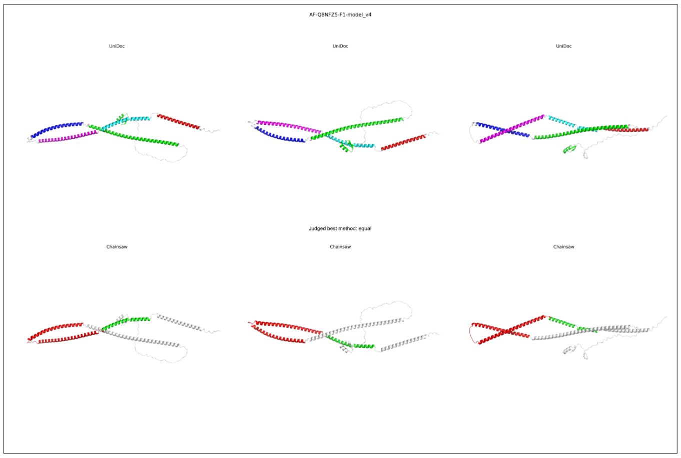
AF-Q8NFZ5-F1-model_v4
UniDoc
UniDoc
UniDoc
Judged best method: equal
Chainsaw
Chainsaw
Chainsaw

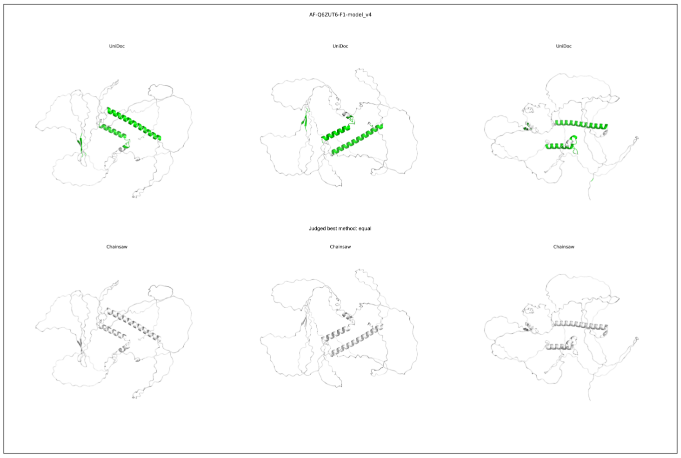
AF-Q6ZUT6-F1-model_v4
UniDoc
UniDoc
UniDoc
Judged best method: equal
Chainsaw
Chainsaw
Chainsaw

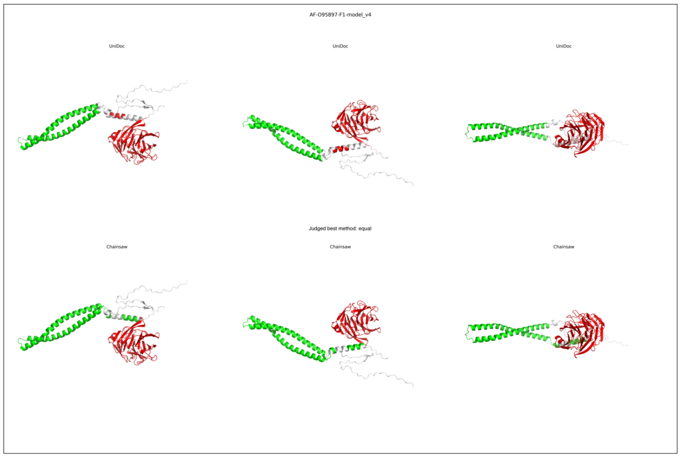
AF-O95897-F1-model_v4
UniDoc
UniDoc
UniDoc
Judged best method: equal
Chainsaw
Chainsaw
Chainsaw

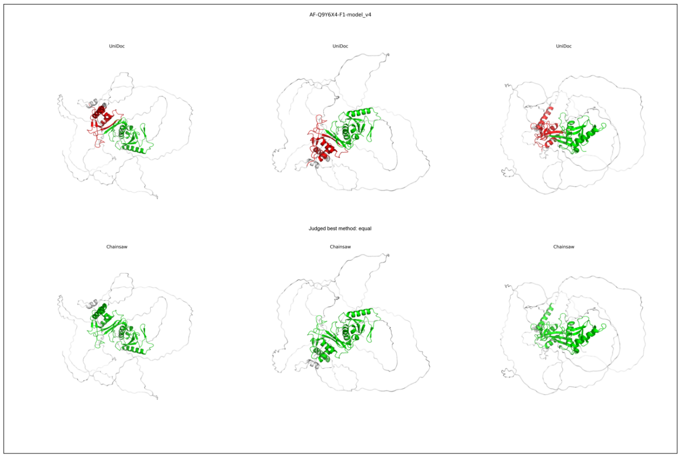
AF-Q9Y6X4-F1-model_v4
UniDoc
UniDoc
UniDoc
Judged best method: equal
Chainsaw
Chainsaw
Chainsaw

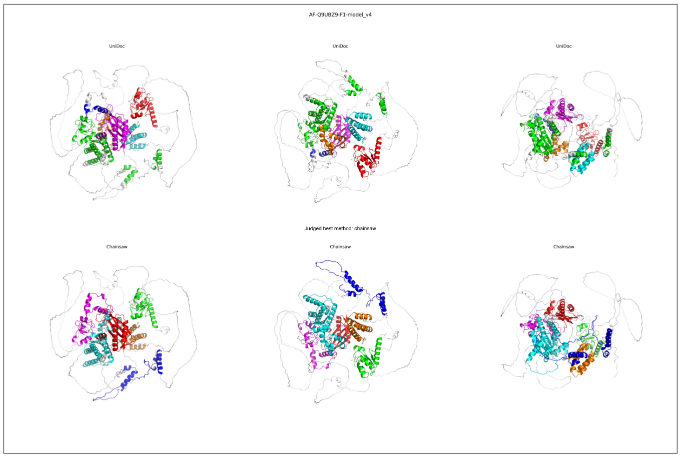
AF-Q9UBZ9-F1-model_v4
UniDoc
UniDoc
UniDoc
Judged best method: chainsaw
Chainsaw
Chainsaw
Chainsaw

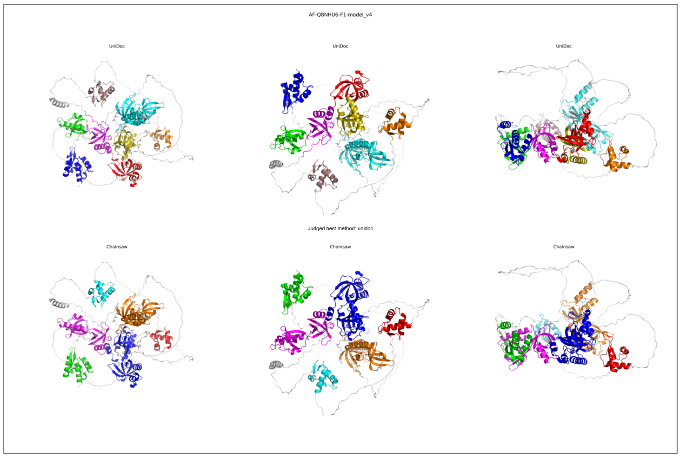
AF-Q8NHU6-F1-model_v4
UniDoc
UniDoc
UniDoc
Judged best method: unidoc
Chainsaw
Chainsaw
Chainsaw

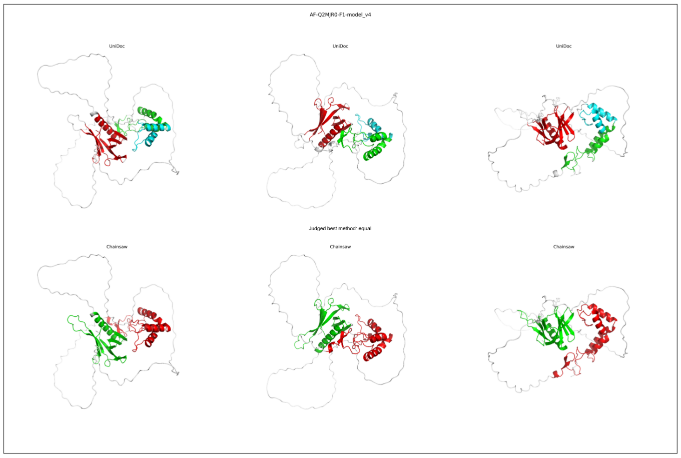
AF-Q2MJR0-F1-model_v4
UniDoc
UniDoc
UniDoc
Judged best method: equal
Chainsaw
Chainsaw
Chainsaw

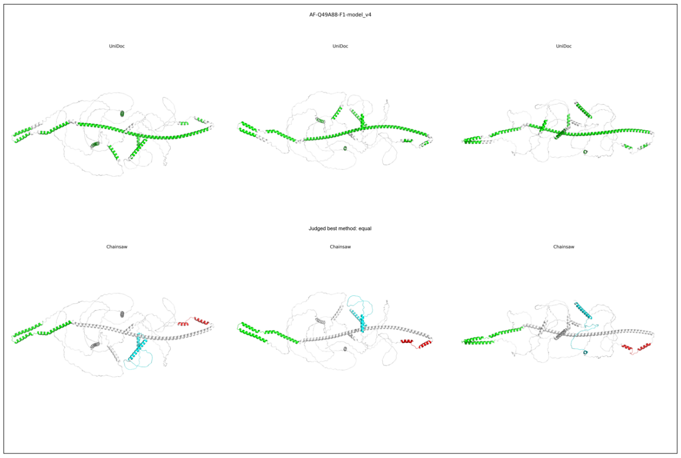
AF-Q49A88-F1-model_v4
UniDoc
UniDoc
UniDoc
Judged best method: equal
Chainsaw
Chainsaw
Chainsaw

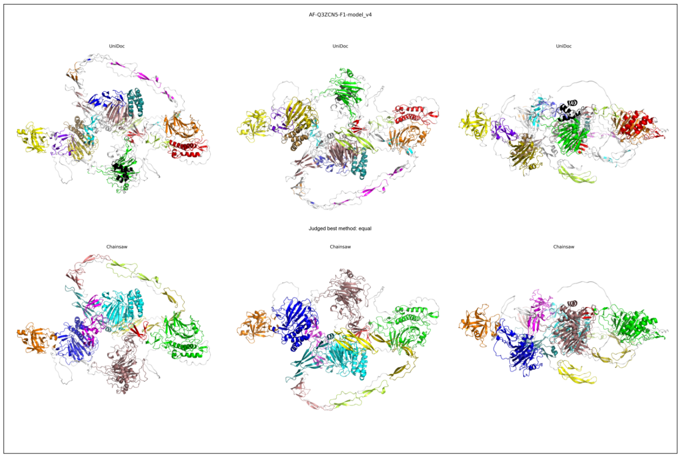
AF-Q3ZCN5-F1-model_v4
UniDoc
UniDoc
UniDoc
Judged best method: equal
Chainsaw
Chainsaw
Chainsaw

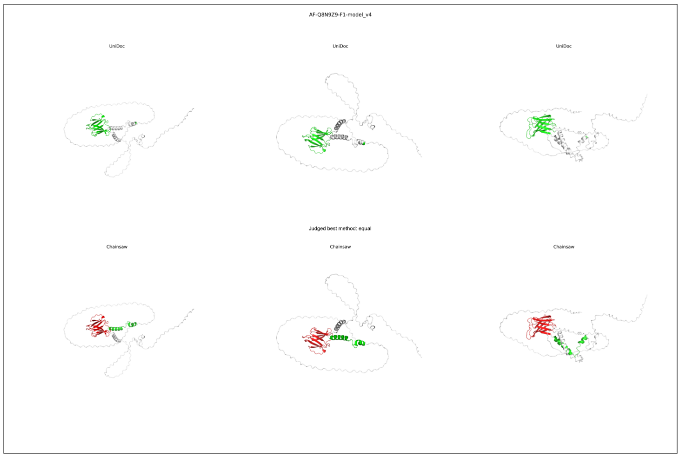
AF-Q8N9Z9-F1-model_v4
UniDoc
UniDoc
UniDoc
Judged best method: equal
Chainsaw
Chainsaw
Chainsaw

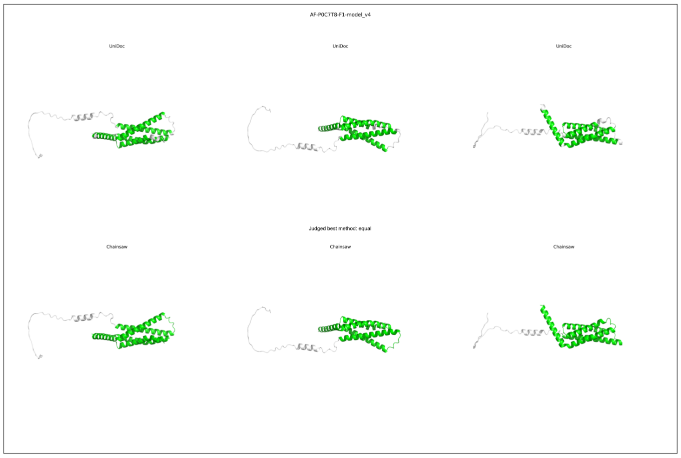
AF-P0C7T8-F1-model_v4
UniDoc
UniDoc
UniDoc
Judged best method: equal
Chainsaw
Chainsaw
Chainsaw

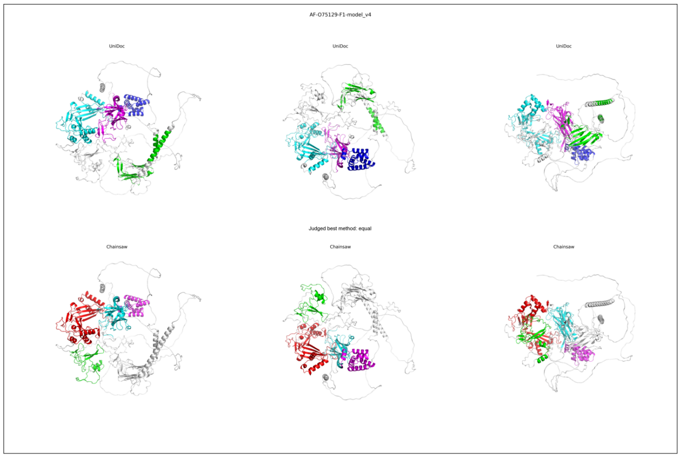
AF-O75129-F1-model_v4
UniDoc
UniDoc
UniDoc
Judged best method: equal
Chainsaw
Chainsaw
Chainsaw

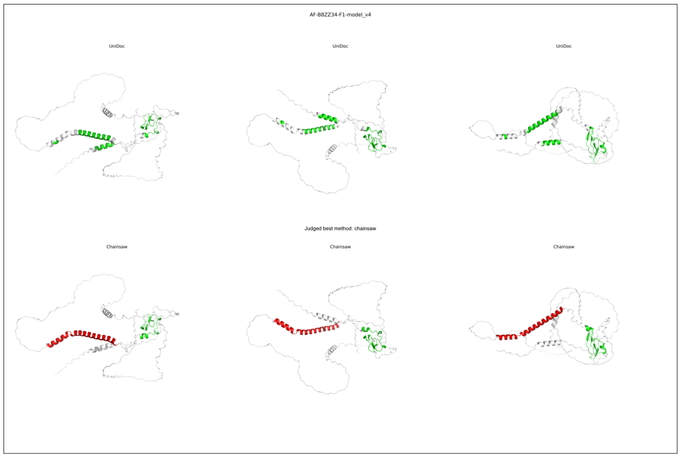
AF-B8ZZ34-F1-model_v4
UniDoc
UniDoc
UniDoc
Judged best method: chainsaw
Chainsaw
Chainsaw
Chainsaw

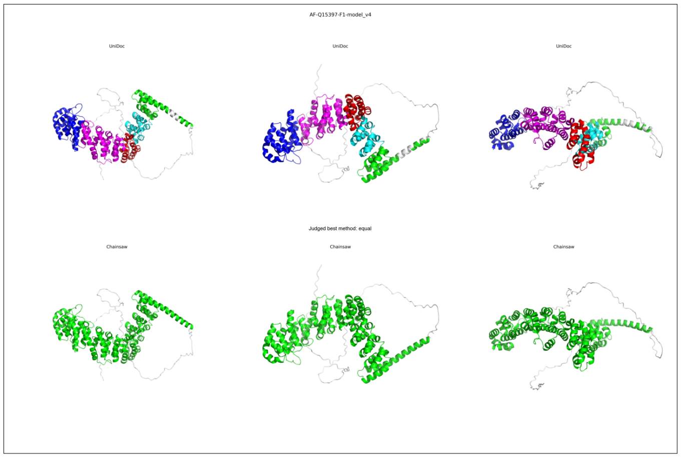
AF-Q15397-F1-model_v4
UniDoc
UniDoc
UniDoc
Judged best method: equal
Chainsaw
Chainsaw
Chainsaw

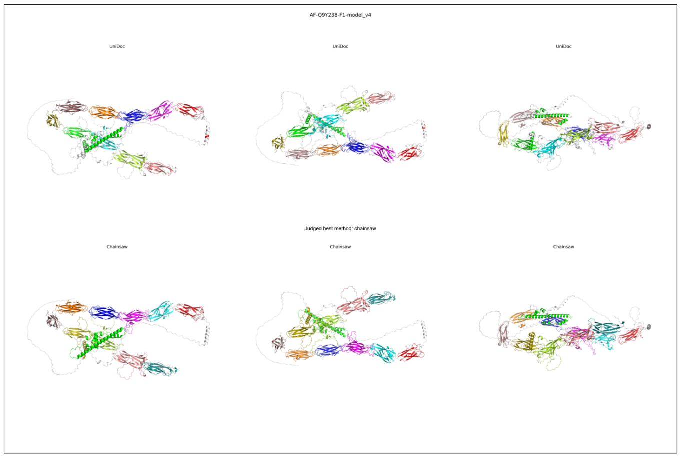
AF-Q9Y238-F1-model_v4
UniDoc
UniDoc
UniDoc
Judged best method: chainsaw
Chainsaw
Chainsaw
Chainsaw

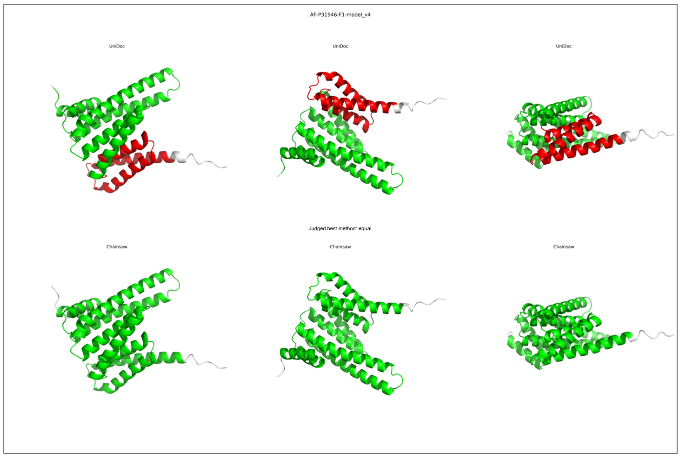
AF-P31946-F1-model_v4
UniDoc
UniDoc
UniDoc
Judged best method: equal
Chainsaw
Chainsaw
Chainsaw

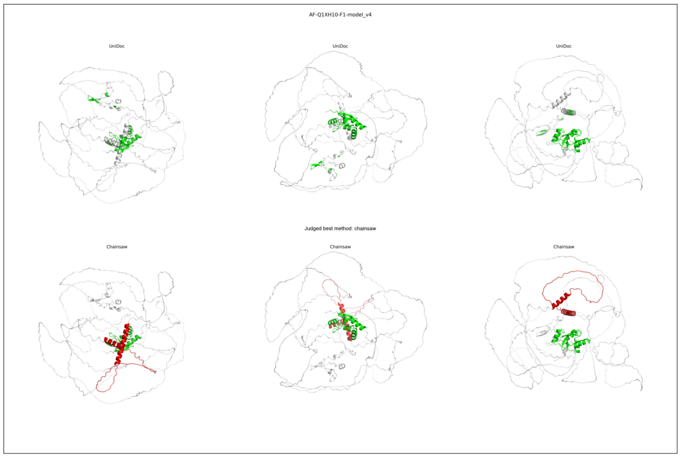
AF-Q1XH10-F1-model_v4
UniDoc
UniDoc
UniDoc
Judged best method: chainsaw
Chainsaw
Chainsaw
Chainsaw

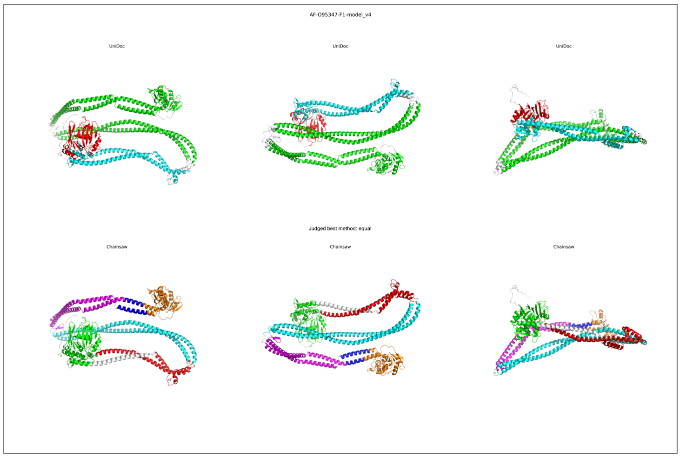
AF-O95347-F1-model_v4
UniDoc
UniDoc
UniDoc
Judged best method: equal
Chainsaw
Chainsaw
Chainsaw

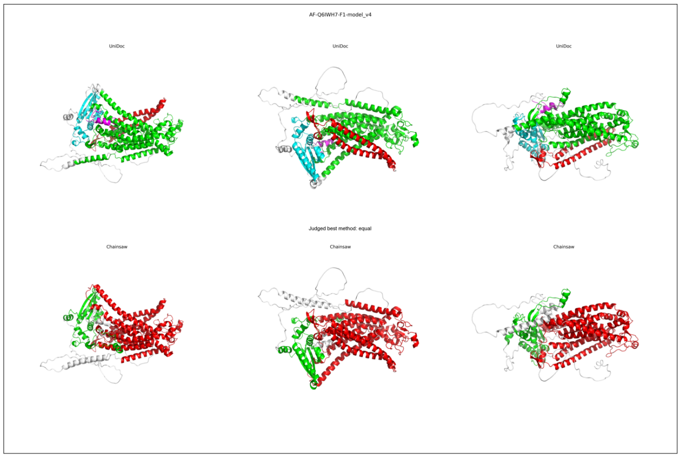
AF-Q6IWH7-F1-model_v4
UniDoc
UniDoc
UniDoc
Judged best method: equal
Chainsaw
Chainsaw
Chainsaw

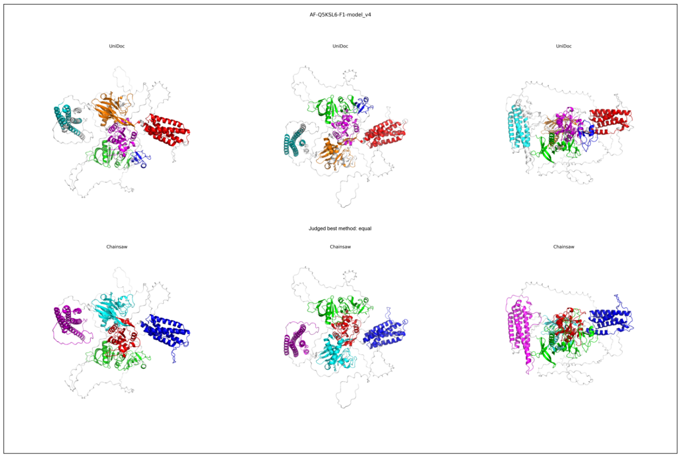
AF-Q5KSL6-F1-model_v4
UniDoc
UniDoc
UniDoc
Judged best method: equal
Chainsaw
Chainsaw
Chainsaw

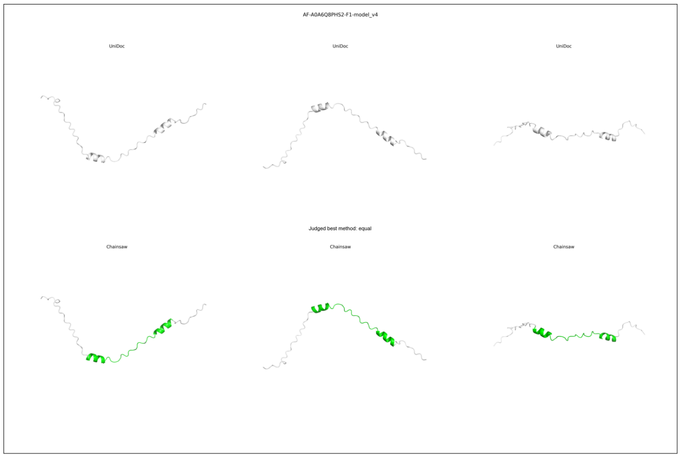
AF-A0A6Q8PHS2-F1-model_v4
UniDoc
UniDoc
UniDoc
Judged best method: equal
Chainsaw
Chainsaw
Chainsaw

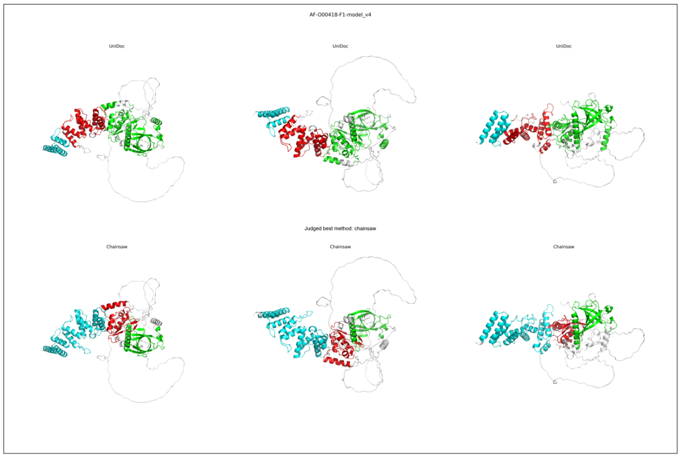
AF-O00418-F1-model_v4
UniDoc
UniDoc
UniDoc
Judged best method: chainsaw
Chainsaw
Chainsaw
Chainsaw

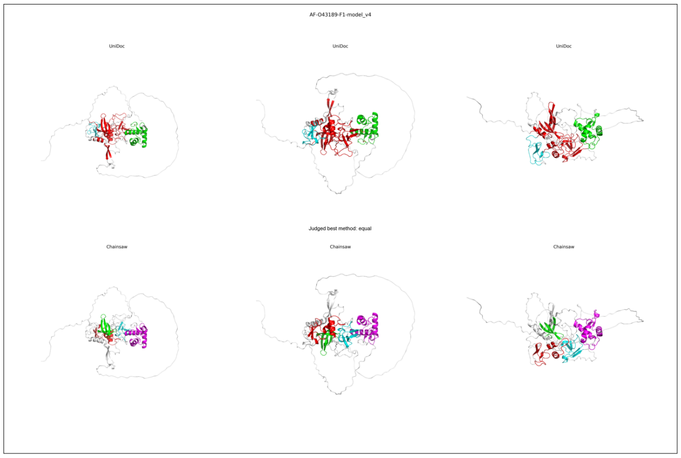
AF-O43189-F1-model_v4
UniDoc
UniDoc
UniDoc
Judged best method: equal
Chainsaw
Chainsaw
Chainsaw

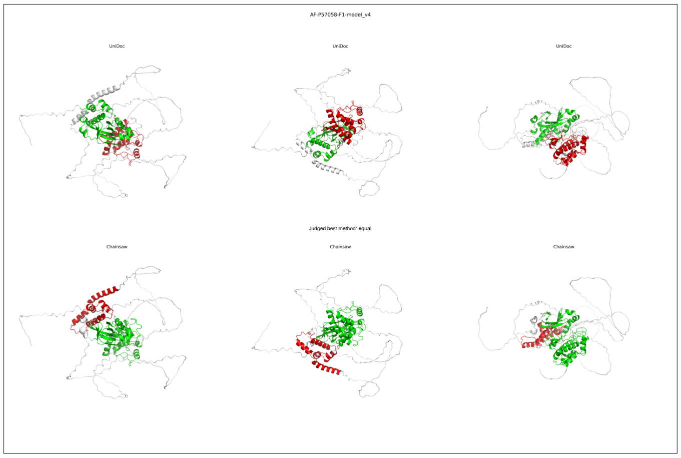
AF-P57058-F1-model_v4
UniDoc
UniDoc
UniDoc
Judged best method: equal
Chainsaw
Chainsaw
Chainsaw

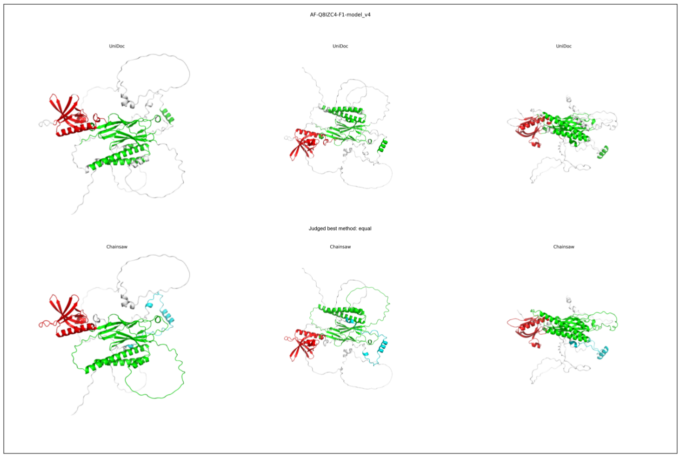
AF-Q8IZC4-F1-model_v4
UniDoc
UniDoc
UniDoc
Judged best method: equal
Chainsaw
Chainsaw
Chainsaw

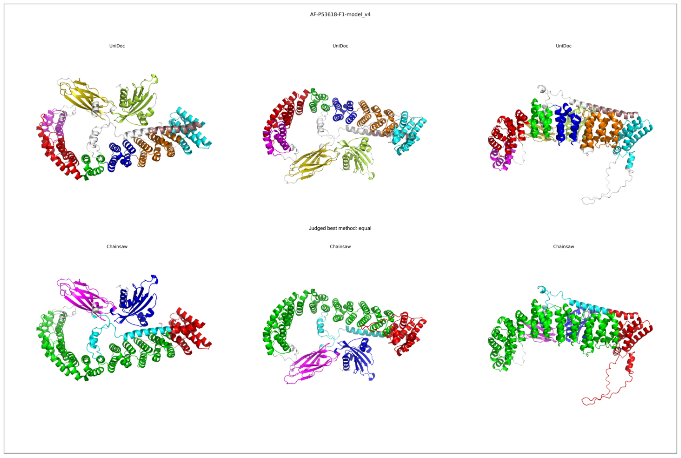
AF-P53618-F1-model_v4
UniDoc
UniDoc
UniDoc
Judged best method: equal
Chainsaw
Chainsaw
Chainsaw

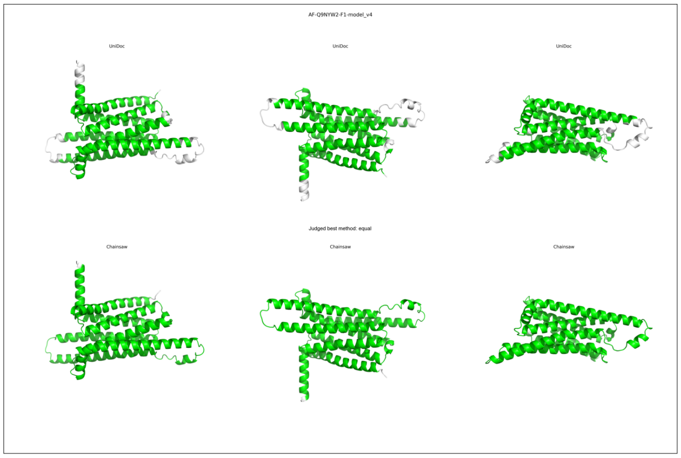
AF-Q9NYW2-F1-model_v4
UniDoc
UniDoc
UniDoc
Judged best method: equal
Chainsaw
Chainsaw
Chainsaw

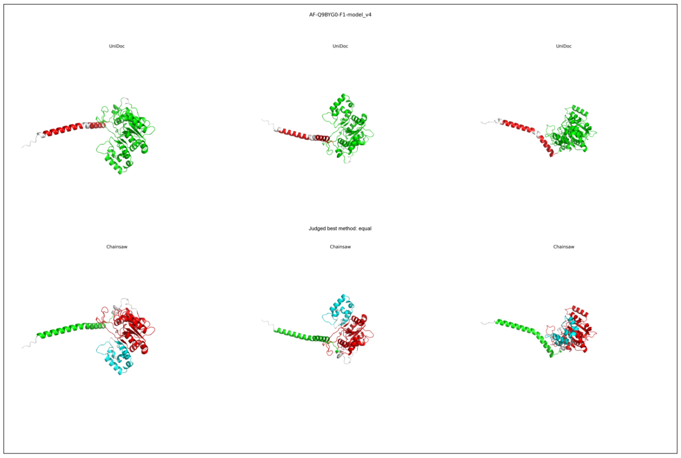
AF-Q9BYG0-F1-model_v4
UniDoc
UniDoc
UniDoc
Judged best method: equal
Chainsaw
Chainsaw
Chainsaw

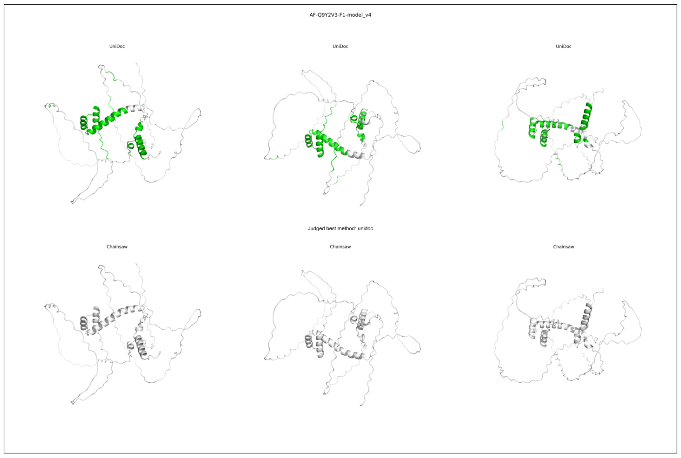
AF-Q9Y2V3-F1-model_v4
UniDoc
UniDoc
UniDoc
Judged best method: unidoc
Chainsaw
Chainsaw
Chainsaw

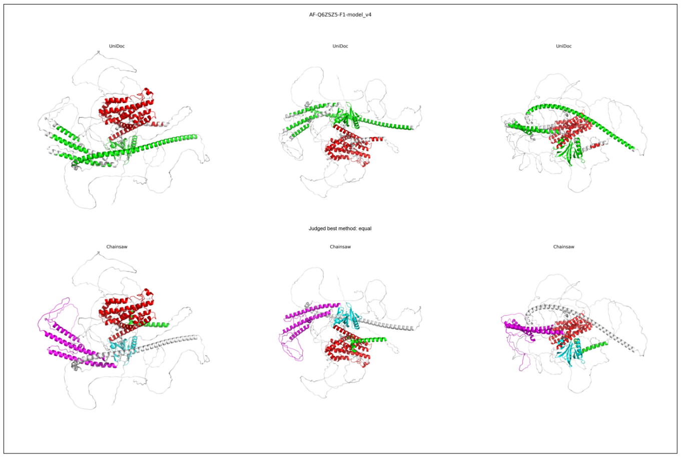
AF-Q6ZSZ5-F1-model_v4
UniDoc
UniDoc
UniDoc
Judged best method: equal
Chainsaw
Chainsaw
Chainsaw

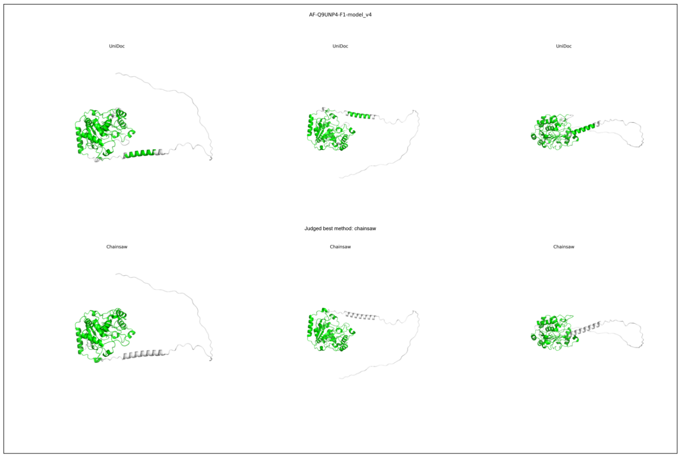
AF-Q9UNP4-F1-model_v4
UniDoc
UniDoc
UniDoc
Judged best method: chainsaw
Chainsaw
Chainsaw
Chainsaw

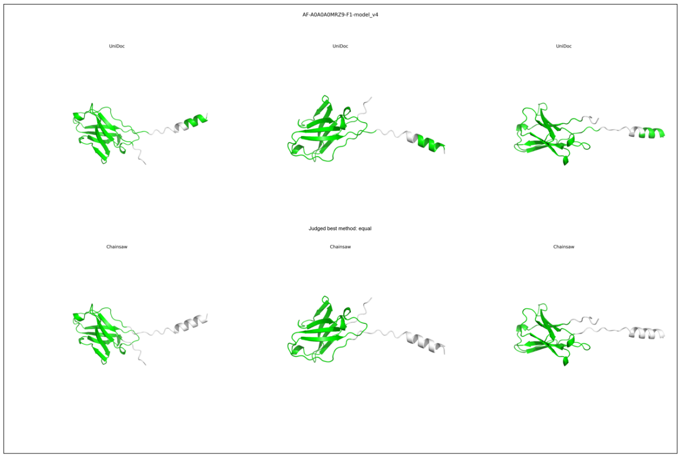
AF-A0A0A0MRZ9-F1-model_v4
UniDoc
UniDoc
UniDoc
Judged best method: equal
Chainsaw
Chainsaw
Chainsaw

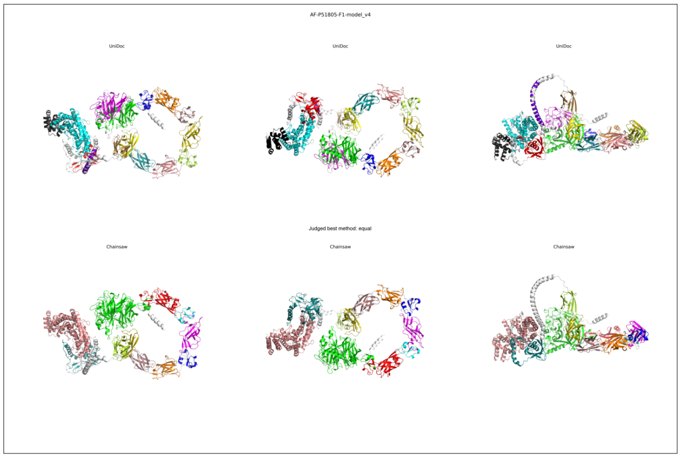
AF-P51805-F1-model_v4
UniDoc
UniDoc
UniDoc
Judged best method: equal
Chainsaw
Chainsaw
Chainsaw

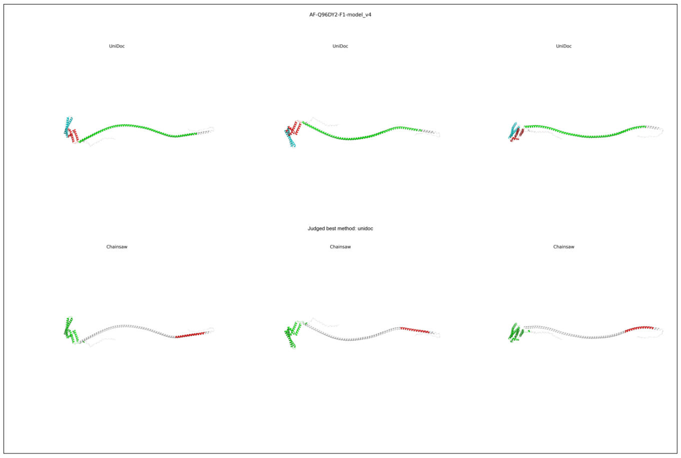
AF-Q96DY2-F1-model_v4
UniDoc
UniDoc
UniDoc
Judged best method: unidoc
Chainsaw
Chainsaw
Chainsaw

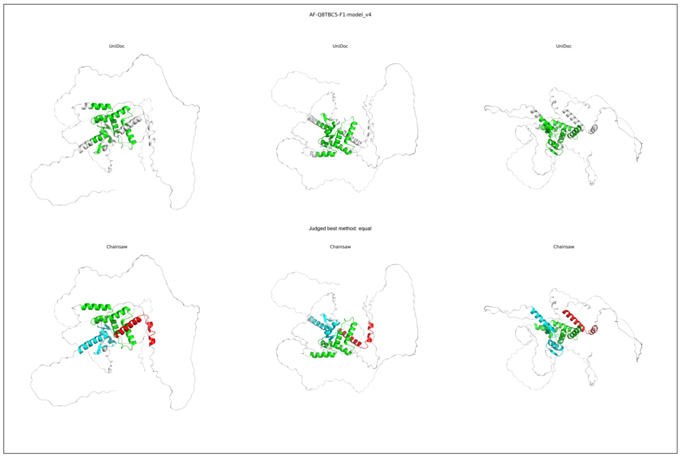
AF-Q8TBC5-F1-model_v4
UniDoc
UniDoc
UniDoc
Judged best method: equal
Chainsaw
Chainsaw
Chainsaw

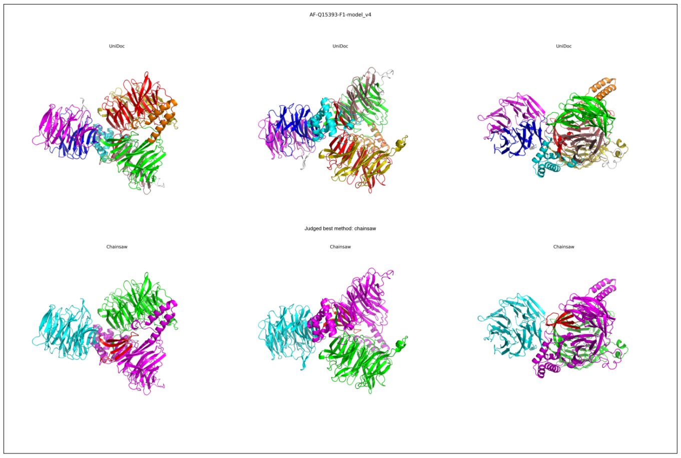
AF-Q15393-F1-model_v4
UniDoc
UniDoc
UniDoc
Judged best method: chainsaw
Chainsaw
Chainsaw
Chainsaw

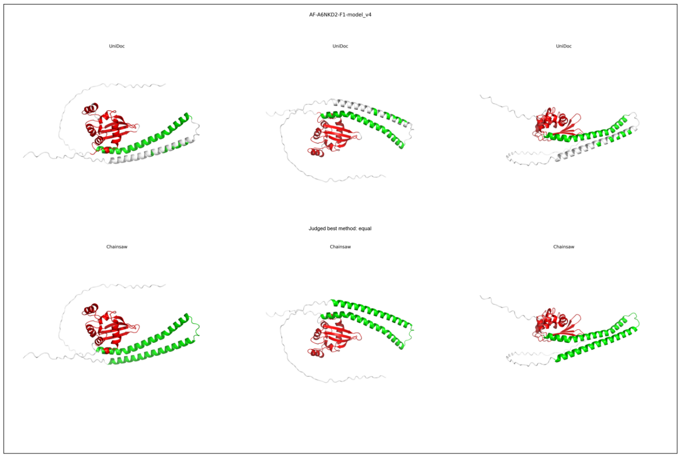
AF-A6NKD2-F1-model_v4
UniDoc
UniDoc
UniDoc
Judged best method: equal
Chainsaw
Chainsaw
Chainsaw

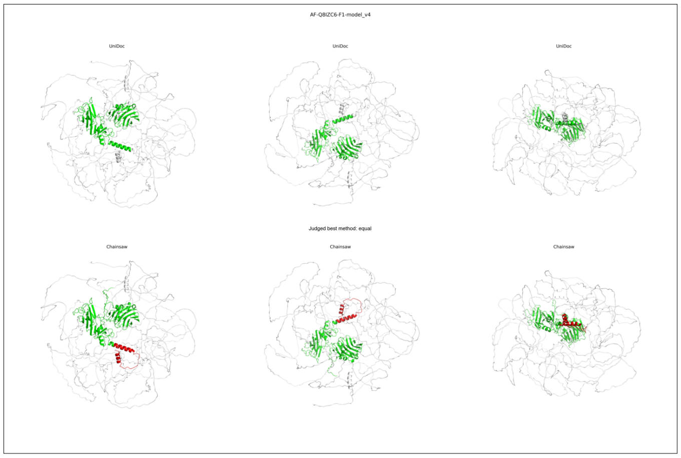
AF-Q8IZC6-F1-model_v4
UniDoc
UniDoc
UniDoc
Judged best method: equal
Chainsaw
Chainsaw
Chainsaw

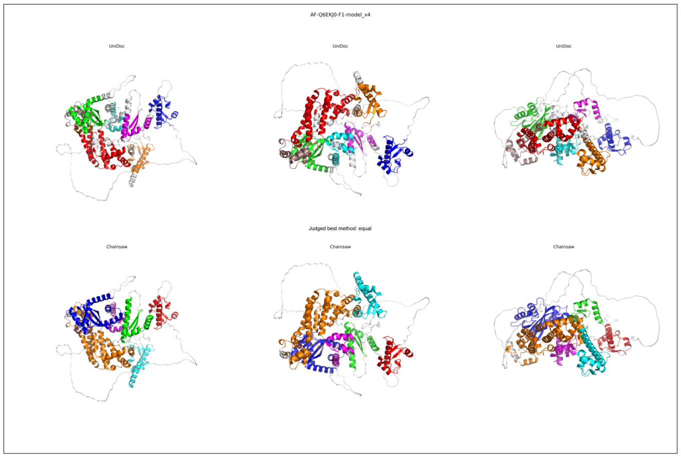
AF-Q6EKJ0-F1-model_v4
UniDoc
UniDoc
UniDoc
Judged best method: equal
Chainsaw
Chainsaw
Chainsaw

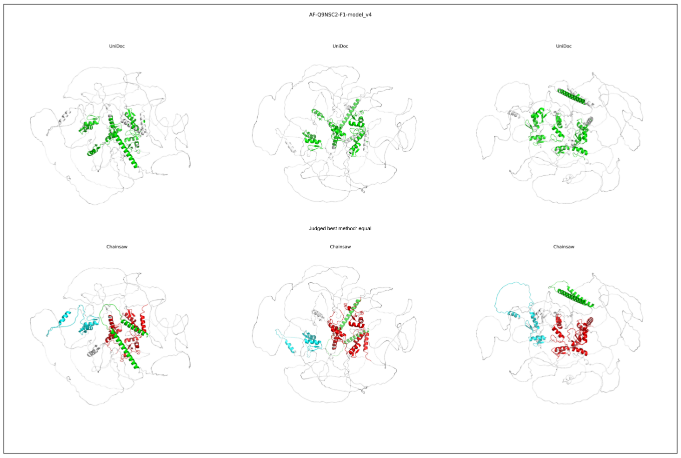
AF-Q9NSC2-F1-model_v4
UniDoc
UniDoc
UniDoc
Judged best method: equal
Chainsaw
Chainsaw
Chainsaw

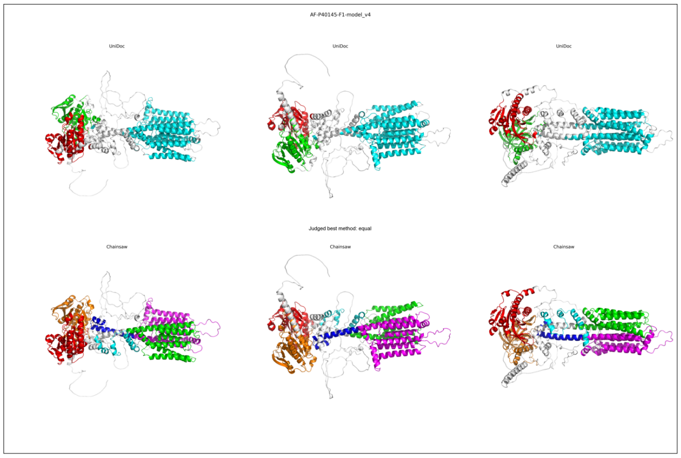
AF-P40145-F1-model_v4
UniDoc
UniDoc
UniDoc
Judged best method: equal
Chainsaw
Chainsaw
Chainsaw

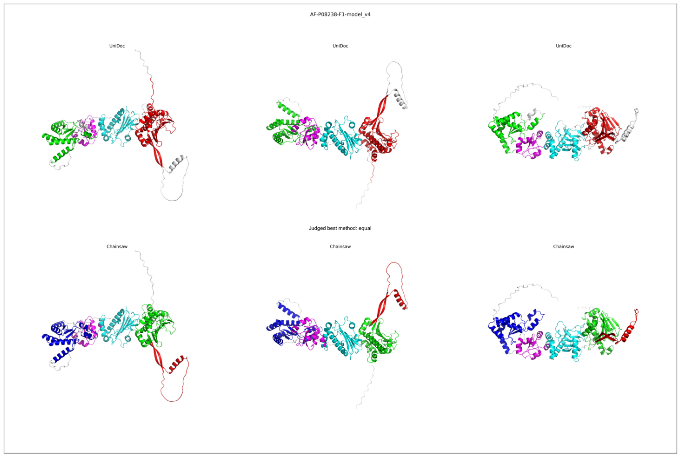
AF-P08238-F1-model_v4
UniDoc
UniDoc
UniDoc
Judged best method: equal
Chainsaw
Chainsaw
Chainsaw

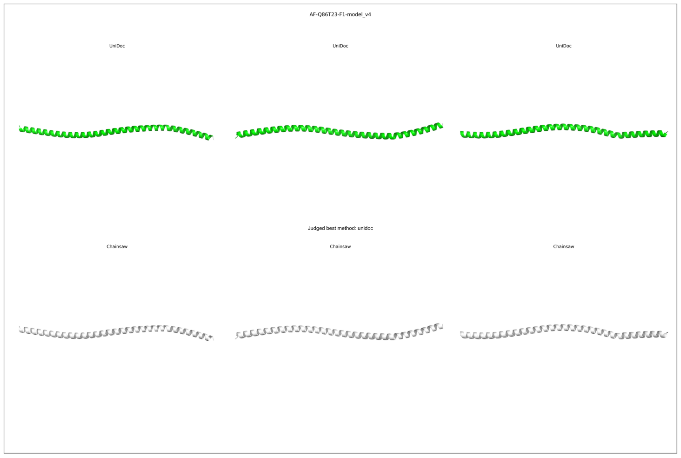
AF-Q86T23-F1-model_v4
UniDoc
UniDoc
UniDoc
Judged best method: unidoc
Chainsaw
Chainsaw
Chainsaw

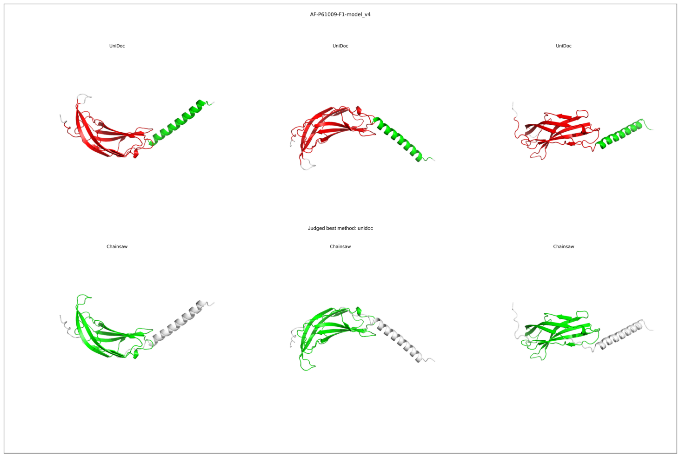
AF-P61009-F1-model_v4
UniDoc
UniDoc
UniDoc
Judged best method: unidoc
Chainsaw
Chainsaw
Chainsaw

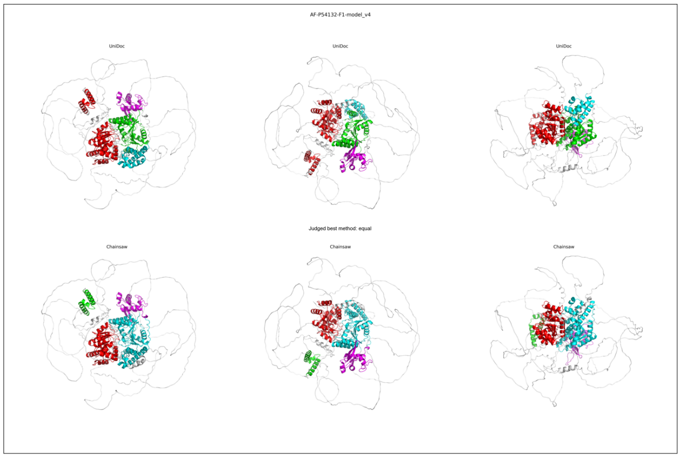
AF-P54132-F1-model_v4
UniDoc
UniDoc
UniDoc
Judged best method: equal
Chainsaw
Chainsaw
Chainsaw

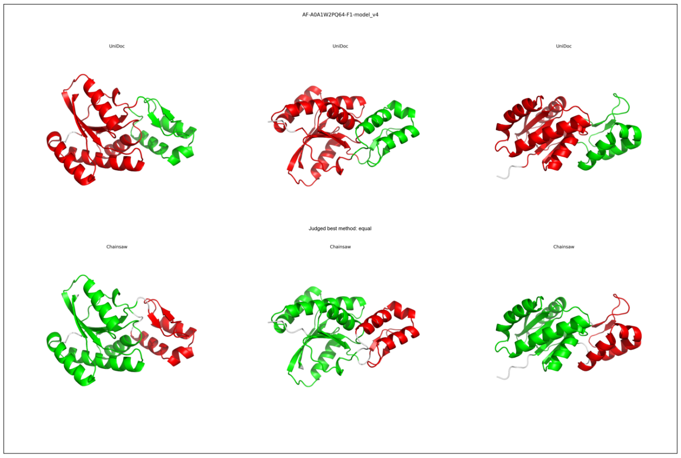
AF-A0A1W2PQ64-F1-model_v4
UniDoc
UniDoc
UniDoc
Judged best method: equal
Chainsaw
Chainsaw
Chainsaw

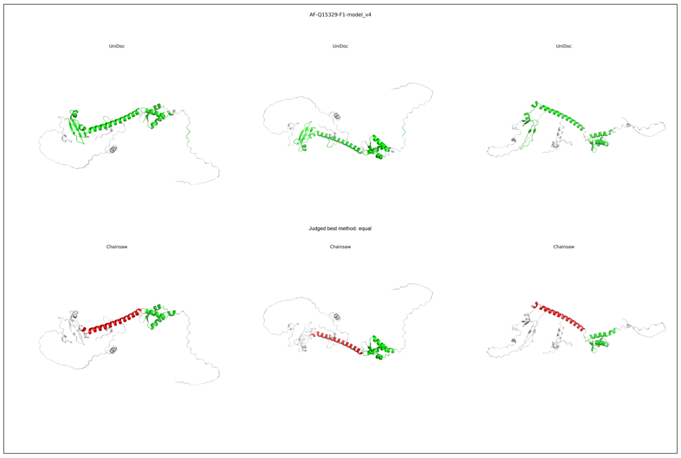
AF-Q15329-F1-model_v4
UniDoc
UniDoc
UniDoc
Judged best method: equal
Chainsaw
Chainsaw
Chainsaw

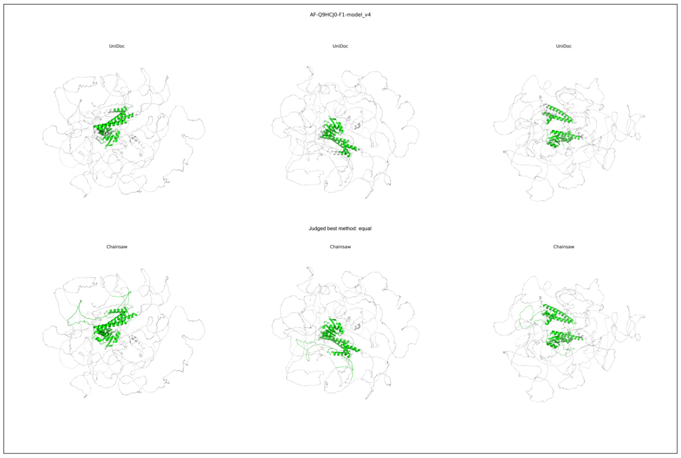
AF-Q9HCJ0-F1-model_v4
UniDoc
UniDoc
UniDoc
Judged best method: equal
Chainsaw
Chainsaw
Chainsaw

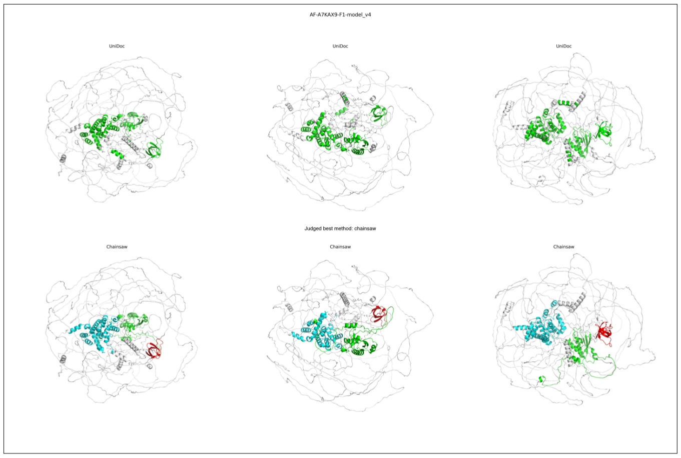
AF-A7KAX9-F1-model_v4
UniDoc
UniDoc
UniDoc
Judged best method: chainsaw
Chainsaw
Chainsaw
Chainsaw

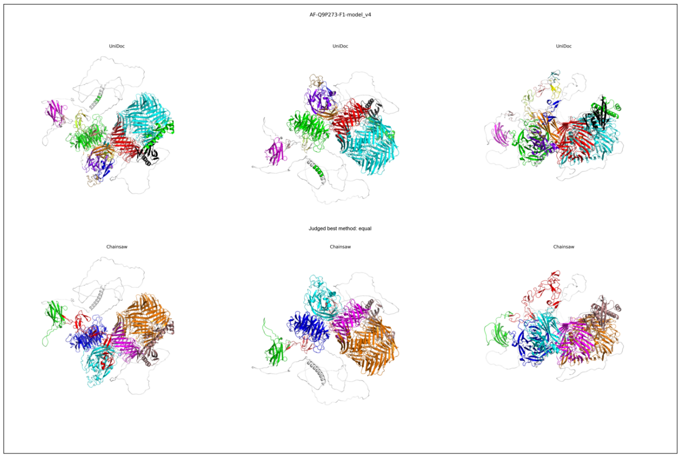
AF-Q9P273-F1-model_v4
UniDoc
UniDoc
UniDoc
Judged best method: equal
Chainsaw
Chainsaw
Chainsaw
